# Concurrent origins of the genetic code and the homochirality of life, and the origin and evolution of biodiversity. Part II: Technical appendix

**DOI:** 10.1101/017988

**Authors:** Dirson Jian Li

## Abstract

Here is Part II of my two-part series paper, which provides technical details and evidence for Part I of this paper (see “Concurrent origins of the genetic code and the homochirality of life, and the origin and evolution of biodiversity. Part I: Observations and explanations” on bioRxiv).

## 0 Overview

The origins of the genetic code, the homochirality of life and the three domains of life are the critical events at the beginning of life. Most direct evidence for these primordial events has been erased; we have to turn to circumstantial evidence in the genome sequences to reproduce the early history of life. In Part I, a roadmap theory is proposed to explain the above events. It indicates that the genetic code and the homochirality of life originated concurrently, which consequently determined the origin and evolution of the biodiversity. Evidence is the final arbiter for this theory. Here in Part II, based on biological and geological data, plenty of evidence has been provided in detail from different perspectives.

Section 1 explains the roadmap for the evolution of the genetic code and the origin of the homochirality of life (Fig 1, 2), which includes initiation, expansion and the ending of the roadmap (Fig 1A, 1B, 2B); the origin of tRNAs (Fig 2A); cooperative recruitments of codons and amino acids (Fig 2B); roadmap symmetry and codon degeneracy (Fig 2C); chiral symmetry broken at the origin of life (Fig 2D); and a summary of evidences that support the roadmap. Section 2 explains the origin of the three domains of life (Fig 3), which includes the relationship between the evolution of the genetic code and the tree of life based on the genomic codon distributions (Fig 3A, S3a5); the origin order of Virus, Bacteria, Archaea and Eukarya according to the recruitment order of codons (Fig 3C); construction of the evolution space; and construction of the tree of life based on the roadmap and genomic codon distributions (Fig 3D, S3d9). And Section 3 explains the Phanerozoic biodiversity curve qualitatively and quantitatively (Fig 4), which includes the split and reconstruction scheme for the Phanerozoic biodiversity curve (Fig 4A, 4B, 4C); the evolution of genome size (Fig S4a2, S4a7); the Phanerozoic climatic curve (Fig S4b2); the Phanerozoic eustatic curve (Fig S4b3); tectonic cause of mass extinctions (Fig 4B); and explanation of the declining origination rate and extinction rate (Fig 4D). Some new nomenclatures are defined when necessary in this paper, which are listed in the Index.

There are 14 subfigures plus 53 supplementary subfigures to explain the origin and evolution of life based on biological and geological data. Numbering rules are as follows: Appendix Section 1−3 correspond to Section 1−3 in the report respectively, and the supplementary subfigures Fig S*αβγ* (*α* = 1, 2, 3, 4; *β* = a, b, c, d; *γ* = 1, 2, 3, …) correspond to subfigures Fig *αβ* (*α* = 1, 2, 3, 4; *β* = A, B, C, D; with corresponding ‘*αβ*’) in the report respectively.

> Section 1 (Appendix Section 1) : Fig 1, 2 (Fig S1, S2)
>
> Fig 1A (Fig S1a1) ; Fig 1B,
>
> Fig 2A (Fig S2a1-3) ; Fig 2B (Fig S2b1-5),
>
> Fig 2C (Fig S2c1-3) ; Fig 2D (Fig S2d1);
>
> Section 2 (Appendix Section 2) : Fig 3 (Fig S3)
>
> Fig 3A (Fig S3a1-6) ; Fig 3B (Fig S3b1-3),
>
> Fig 3C (Fig S3c1-2) ; Fig 3D (Fig S3d1-10);
>
> Section 3 (Appendix Section 3) : Fig 4 (Fig S4)
>
> Fig 4A (Fig S4a1-8) ; Fig 4B (Fig S4b1-7),
>
> Fig 4C (Fig S4c1) ; Fig 4D (Fig S4d1-3).

## 1 The roadmap for the evolution of the genetic code and the origin of homochirality of life

### 1.1 Initiation of the roadmap

The evolution of the genetic code can be divided into three stages: the initiation stage, the expansion stage and the ending stage.

In the beginning, there was an *R* (*R* denotes purine) single-stranded DNA *Poly G* (Fig 1A, 1B #1). By complementary base pairing, a *YR* (*Y* denotes pyrimidine) double-stranded DNA *Poly C* · *Poly G* formed. And by base triplex *CG* * *G*, a *YR* * *R*1 triple-stranded DNA *Poly C* · *Poly G* * *Poly G* formed (Fig 1A, 1B #1). The third *R*1 strand *Poly G* separated out of this *YR* * *R*1 triple-stranded DNA, which then formed a new *Y*1*R*1 double-stranded DNA *Poly C* · *Poly G*. So far, there was only initial codon pair *GGG* · *CCC* (Fig 1A, 1B #1).

Starting from this initial #1 codon pair and by the base substitutions throughout the four hierarchies, the base pairs from #2 to #32 recruited so that the whole genetic code table is fulfilled (Fig 1). In this paper, the technical process is illustrated in the roadmap for the evolution of the genetic code (or the roadmap for short) (Fig 1A). The relative stabilities of all the 16 possible base triplexes are listed from weak to strong as follows [1] [2]:

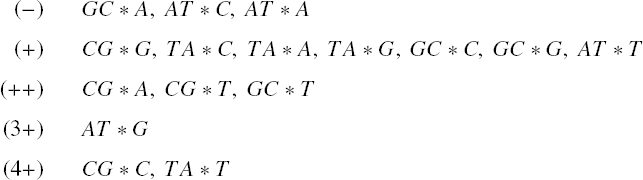

The relative stability of base triplexes played a crucial role in the base substitutions along the roadmap.

A weaker base triplex tends to be substituted spontaneously by a stronger one, in the course of repeated combinations and separations of strands for the triple-stranded DNA. Only three kinds of substitutions of base triplexes are practically required in the roadmap: substitution of (+) *CG* * *G* by (++) *CG* * *A*, with the transition from *G* to *A* in the third *R* strand; substitution of (+) *CG* * *G* by (4+) *CG* * *C*, with the transversion from *G* to *C* in the third *R* strand; substitution of (+) *GC* * *C* by (++) *GC* * *T*, with the transition from *C* to *T* in the third *R* strand (Fig 1A, S1a1). The transition from *G* to *A* is of the most common substitution in the roadmap (Fig 1A). All the 8 codon pairs in *Route* 0 were formed by the transition from *G* to *A*. Considering the dislocation of bases in *Poly C* · *Poly G* * *Poly G*, only one transversion from *G* to *C* was enough to blazed a new path other than *Route* 0 by bringing the starting codon pairs *GGC* ·*GCC*, *GCG* ·*CGC* and *CGG* ·*CCG* for *Route* 1, *Route* 2 and *Route* 3 at positions #2, #7 and #10 in the roadmap, respectively (Fig 1A). Considering the dislocation of bases similarly, only one transition from *C* to *T* was enough to blaze a new path so as to form the remaining codon pairs by substitutions at positions #6, #19 and #12 in the roadmap, respectively (Fig 1A). The length of the genetic code are not greater than 3 bases because such supposed genetic code cannot be fulfilled by the demand of increasing stability of the base triplexes; namely the unavoidable instabilities of the base triplexes prohibited the k-base genetic code (*k* * 3) in practice.

In the initiation stage of the roadmap, the codon pairs from #1 to #6 were recruited by the roadmap, which constituted the initial subset of the genetic code:

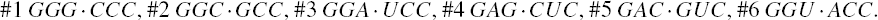

And in this stage were recruited the earliest 9 amino acids in order: 1*Gly*, 2*Ala*, 3*Glu*, 4*Asp*, 5*Val*, 6*Pro*, 7*S er*, 8*Leu*, 9*T hr*, all of which belong to Phase *I* amino acids [3] [4]. Although the initial subset is small, the two essential features of the roadmap, pair connection and route duality, had taken shape in this initial stage (Fig 1A, 2B).

Pair connection is an essential feature of the roadmap. The connected codon pairs in the roadmap generally encode a common amino acid (Fig 1A, 2C). For instance, the pair connection #1−*Gly*−#2 indicates that both *GGG* in #1 and *GGC* in #2 encode the common amino acid *Gly*. Pair connections reveal the close relationship between the recruitment of the codons and the recruitment of the amino acids.

Route duality is another essential feature of the roadmap, which shows the relationship of pair connections between different routes (Fig 1A, 2C). For instance, the route duality

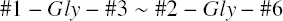

indicates that the pair connection #1 − *Gly* − #3 in *Route* 0 and the pair connection #2 − *Gly* − #6 in *Route* 1 are dual, which encode the common amino acid *Gly*. Route dualities generally exist between *Route* 0 and *Route* 3, or between *Route* 1 and *Route* 2 (Fig 2C).

### 1.2 Expansion of the genetic code along the roadmap

The genetic codes evolved along the four routes *Route* 0 − 3 respectively; and the 8 codon pairs in each route evolved in the order of the four hierarchies *Hierarchy* 1 − 4 respectively (Fig 1A). The roadmap can be divided into two groups: the early hierarchies *Hierarchy* 1 − 2 and the late hierarchies *Hierarchy* 3 − 4. It can also be divided into two groups: the initial route *Route* 0 (all-purine codons with all-pyrimidine codons) and the expanded routes *Route* 1 − 3 (Fig 1A, S3d2). These groupings will be used to explain the origin of the three domains in Subsection 2.3.

The genetic codes expanded spontaneously from the initial subset in the expansion stage of the roadmap. Each of the 6 codon pairs in the initial subset expanded to three codon pairs, respectively, by route dualities.

Details are as follows. The codon pair #2 in the initial subset expanded to the three continual codon pairs #7, #8 and #9 by route duality

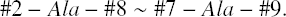

The codon pair #1 in the initial subset expanded to the three continual codon pairs #10, #11 and #12 by route duality

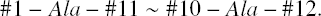

The codon pair #3 in the initial subset expanded to the three continual codon pairs #13, #14 and #15 by route duality

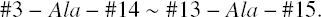

The codon pair #6 in the initial subset expanded to the three continual codon pairs #16, #17 and #18 by route duality

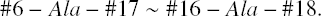

The codon pair #5 in the initial subset expanded to the three codon pairs #19, #24 and #26 by route duality

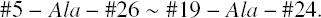

The codon pair #4 in the initial subset expanded to the three codon pairs #20, #25 and #27 by route duality

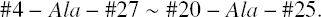

According to the base substitutions in the roadmap, the recruitment order of the codon pairs from #1 to #32 is as follows (Fig 1A):

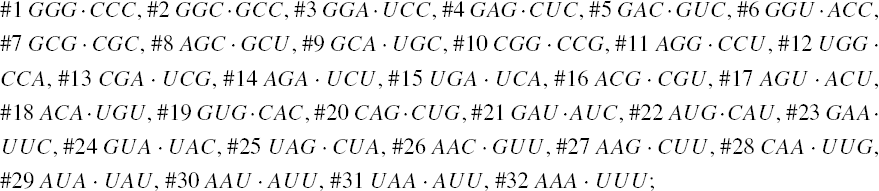

and the recruitment order of the amino acids from *No*.1 to *No*.20 is as follows (Fig 1A):

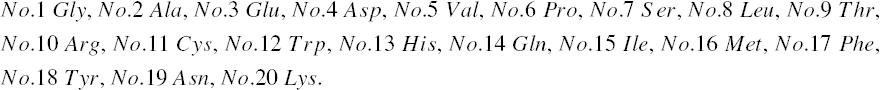

The recruitment order of the codon pairs and the recruitment order of the amino acids are intricately well organised and coherent, according to the subtle roadmap (Fig 1A, 2B). In the initiation stage, firstly, the amino acid *No*.1 was recruited with the codon pair #1, remaining a vacant position. Subsequently, *No*.1 and *No*.2 were recruited with the codon pair #2; *No*.1 was recruited with the codon pair #3, remaining a vacant position; *No*.3 was recruited with the codon pair #4, remaining a vacant position; *No*.4 and *No*.5 were recruited with the codon pair #5; *No*.6 filled up the vacant position of #1; *No*.7 filled up the vacant position of #3; *No*.8 filled up the vacant position of #4; *No*.1 and *No*.9 were recruited with the codon pair #6 (Fig 2B). Thus the framework of the genetic code had been established at the end of the initiation stage. Hereafter, it was always proper to recruit two amino acids that had been already recruited or were about to be recruited next, such that no more positions remained vacant. Details are as follows. *No*.2 and *No*.10 amino acids were recruited with the codon pair #7; and subsequently, *No*.2 and *No*.7 were recruited with #8; *No*.2 and *No*.11 were recruited with #9; *No*.6 and *No*.10 were recruited with #10; *No*.6 and *No*.10 were recruited with #11; *No*.6 and *No*.12 were recruited with #12; *No*.7 and *No*.10 were recruited with #13; *No*.7 and *No*.10 were recruited with #14; *No*.7 and *stop* were recruited with #15; *No*.9 and *No*.10 were recruited with #16; *No*.7 and *No*.9 were recruited with #17; *No*.9 and *No*.11 were recruited with #18; *No*.5 and *No*.13 were recruited with #19; *No*.8 and *No*.14 were recruited with #20; *No*.4 and *No*.15 were recruited with #21; *No*.13 and *No*.16 were recruited with #22; *No*.3 and *No*.17 were recruited with #23; *No*.5 and *No*.18 were recruited with #24; *No*.8 and *stop* were recruited with #25; *No*.5 and *No*.19 were recruited with #26; *No*.8 and *No*.20 were recruited with #27; *No*.8 and *No*.14 were recruited with #28; *No*.15 and *No*.18 were recruited with #29; *No*.15 and *No*.19 were recruited with #30; *No*.8 and *stop* were recruited with #31; *No*.17 and *No*.20 were recruited with #32 (Fig 2B).

Take for example from #1 to #29, the evolution of the genetic code along the roadmap can described in details as follows (Fig 1A, 1B). Starting from the position #1 (Fig 1B #1), an *R* single-stranded DNA brought about a *YR* double-stranded DNA; next, the *YR* double-stranded DNA brought about a *YR* \**R*1 triple-stranded DNA (the number 1 denotes #1, similar below); next, an *R*1 single-stranded DNA departed from the *YR* * *R*1 triple-stranded DNA; next, the *R*1 single-stranded DNA brought about a *R*1*Y*1 double-stranded DNA. Thus, the codon pair *GGG*·*CCC* were achieved at #1. At the beginning of #7 (Fig 1B #7), the *R*1*Y*1 double-stranded DNA was renamed as *Y*1*R*1 double-stranded DNA, where the 180? rotation did not change the right-handed helix; next, the *Y*1*R*1 double-stranded DNA brought about a *Y*1*R*1 * *R*7 triple-stranded DNA, through the transversion from *G* to *C*, where the stability (+) of *CG* * *G* increased to the stability (4+) of *CG* * *C*; next, an *R*7 single-stranded DNA departed from the *Y*1*R*1\**R*7 triple-stranded DNA; next, the *R*7 single-stranded DNA brought about a *R*7*Y*7 double-stranded DNA. Thus, the codon pair *GCG* · *CGC* were achieved at #7. The case of #19 is similar to #7 (Fig 1B #19); the codon pair *GT G* · *CAC* were achieved through the transition from *C* to *T*, where the stability (+) of *GC* \**C* increased to the stability (2+) of *GC* \**T*. The case of #24 is also similar to #7 (Fig 1B #24); the codon pair *GT A* · *T AC* were achieved through the common transition from *G* to *A*, where the stability (+) of *CG* * *G* increased to the stability (2+) of *CG* * *A*. At the position #29 (Fig 1B #29), the codon pair *GCG* · *CGC* in *Y*24*R*24 are non-palindromic in consideration that both *GCG* and *CGC* do not read the same backwards as forwards. In this case, a reverse operation is necessary so that the obtained codon pair *CAT* · *AT G* in *y*24*r*24 read reversely the same as the codon pair *T AC* · *GT A* in *Y*24*R*24. The process from *y*24*r*24 to *R*29*Y*29 is still similar to the case of #7; the codon pair *AT A* · *T AT* were achieved through the transition from *G* to *A*, where the stability (+) of *CG* * *G* increased to the stability (2+) of *CG* * *A*. Other processes in the roadmap are similar to the above example (Fig 1A, 1B). The reverse operation is unnecessary for the cases #2, #7, #10, #11, #3, #4, #16, #9, #19, #27, #23, #22, #24 after palindromic codon pairs and the last one #32 (Fig 1A), whereas the reverse operation is necessary for the remaining cases #5, #6, #8, #12, #13, #14, #15, #17, #18, #20, #21, #25, #26, #28, #29, #30, #31 (Fig 1A).

### 1.3 The ending of the roadmap

Nowadays the genetic code table had been expanded from the 6 codon pairs in the initial subset to the 6 + 18 codon pairs by route duality; the remaining 8 codon pairs were recruited into the genetic code table in the ending stage of the roadmap (Fig 1A, 2B). There were 2 codon pairs remained in each of the four routes *Route* 0 − 3 respectively. They satisfied pair connections as follows: #23 − *Phe* − #32, #21 − *Ile* − #30, #22 − *Met*/*Ile* − #29, #28 − *Leu* − #31 (Fig 2B). Two of them satisfied route duality (Fig 2B):

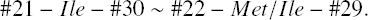

The last two stop codons appeared in the pair connection #25 − *stop* − #31 (Fig 1A, 2B). When the last two amino acids were recruited through the base pairs #26 − *Asn* − #30 and #27 − *Lys* − #32, the codon *U AG* at #25 had to be selected as a stop codon. Due to no participation of corresponding tRNA, the codon *U AA* at #31 was selected as the last stop codon.

The non-standard codons also satisfy codon pairs and route dualities in the roadmap (Fig 1A). The codon pairs pertaining to non-standard codons are as follows: #11 − *Arg* (*S er*, *stop*) − #14, #4 − *Leu* (*T hr*) − #27 in *Route* 0; none in *Route* 1; #22 − (*Met*) − #29 in *Route* 2; #20 − *Leu* (*T hr*, *Gln*) − #25, #12 − (*T r p*) − #15, #25 − *stop* (*Gln*)/*Leu* − #31, #28 − *Leu* (*Gln*) − #31 in *Route* 3. Majority of non-standard codons appear in the last *Route* 3 (Fig 1A). Route dualities of non-standard codons exist between *Route* 0 and *Route* 3 (Fig 1A):

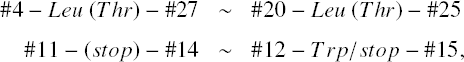

where the first stop codon *UGA* at #15 is dual to the non-standard stop codons in *Route* 0.

The stabilities of base triplexes played crucial roles in the evolution of the genetic code. It had been emphasised that the roadmap followed the strict rule that the stabilities of base triplexes increase gradually (Fig S1a1). Also note that the roadmap had tried its best to avoid the unstable triplex DNA. Among the 16 possible base triplexes, there are three relatively unstable base triplexes: *GC* * *A*, *AT* * *C* and *AT* * *A*. So the statistical ratio of instability for the base triplexes is 3/16. However, the ratio of instability for the base triplexes in the roadmap is much smaller. There are 49 triplex DNAs through #1 to #32 in the roadmap, which involve 3 × 49 = 147 base triplexes (Fig 1A). The relatively unstable base triplexes *GC* * *A* and *AT* * *C* have not appeared in the roadmap; only the relatively unstable base triplex *AT* * *A* has appeared inevitably for 7 times in the reverse operations so as to fulfil all the permutations of 64 codons (Fig 1A). The ratio of instability 7/147 in the roadmap is much smaller than the ratio of instability 3/16 by the statistical requirement. When the relatively unstable *AT* * *A* appears at the positions #15, #17, #21, #25, #29, #30 and #31, both stabilities of the other two base triplexes in the triplex DNA are (4+) (Fig 1A), which compensates the instability of the triplex DNA to some extent. The amino acid *Ile*, whose degeneracy uniquely is three, occupied three positions #21, #29 and #30 among those 7 positions. And the three stop codons occupied other three neighbour positions #15, #25 and #31 (Fig 1A). The route dualities played significant roles in the expansion stage (Fig 1A, 2B). The hence remnant codons were chosen as the stop codons. The first stop codon *UGA* appeared at the position #15 where the relatively unstable *AT* * *A* appeared firstly (Fig 1A), hence it is possible that the instability might be helpful to achieve the function of stop codons. The stop codon appeared as early as the midway of the evolution of the genetic code (Fig 1A, 2B), which indicates that the genetic code had been taken shape around the midway to promote the formation of the primitive life. Not until the fulfilment of the genetic code, did the translation efficiency increase notably by recognising all the 64 codons.

### 1.4 Origin of tRNAs

The roadmap illustrates the coevolution of the genetic code and the amino acids, in which tRNAs played mediate roles. The recruitment and expansion of tRNAs underlay pair connections and route dualities. This subsection explains the origin of tRNAs according to the roadmap.

There are 16 branch nodes and 16 leaf nodes in the roadmap (Fig 1A, 2A). The leaf nodes are as follows: #3, #14, #27, #32 in Route 0; #8, #17, #26, #30 in Route 1; #9, #18, #22, #29 in Route 2; #13, #15, #28, #31 in *Route* 3. And the branch nodes are as follows: #1, #4, #11, #23 in *Route* 0; #2, #5, #6, #21 in *Route* 1; #7, #16, #19, #24 in *Route* 2; #10, #12, #20, #25 in *Route* 3. For each branch node, the third strand of the triplex nucleic acids is of DNA (Fig 1A). Nevertheless the third strand is an RNA strand (Fig 2A). Namely the first two DNA strands are combined by a third RNA strand. Such triplex nucleic acids are of two types: the triplex nucleic acid *YR* * *Y*_*t*_ and the triplex nucleic acid *YR* * *R*_*t*_ (Fig 2A), where the subscript *t* denotes tRNA. After departing from the triplex nucleic acids, the combination of the two complementary single RNA strands *Y*_*t*_ and *R*_*t*_ fold into a cloverleaf-shaped tRNA [5] [6] [7] [8] [9], which acquired a certain anticodon corresponding to its position in the roadmap (Fig S2a3).

For example, the tRNA *t*1 can form from the two triplex nucleic acids *Y*1*R*1 * *Y*_*t*_1 and *Y*1*R*1 * *R*_*t*_1 in the position #1(Fig 2A #1). The triple code *CCC* in the strand *Y*_*t*_1 is palindromic, while the triple code *GGG* in the strand *R*_*t*_1 is palindromic. The two complementary strands *Y*_*t*_1 and *R*_*t*_1 can combine into a cloverleaf-shaped tRNA *t*1 by joining together, complementary pairing and folding (Fig S2a3). Thus anticodon arm of *t*1 contains the anticodon *CCC*, which corresponds to *Gly*; namely *t*1 transports *Gly*. As another example, the codon in the position #11 is not palindromic, where two tRNAs *t*7 and *t*10 are folded respectively (Fig 2A #11). Two complementary single RNA strands *Y*_*t*_11 and *R*_*t*_11 form by reverse cooperation. They combine into the tRNA *t*7 by joining together, complementary pairing and folding, which contains the anticodon *GGA* and carry *S er* (Fig S2a3). Furthermore, another two complementary single RNA strands *Y*_*t*_11 and *R*_*t*_11 can form. They combine into the tRNA *t*10, which contains the anticodon *CCU* and carry *Arg* (Fig S2a3).

There are 4 palindromic codons: #1 *CCC* · *GGG*, #4 *CUC* · *GAG*, #7 *CGC* · *GCG*, #19 *CAC* · *GUG* among the 16 branch nodes in the roadmap (Fig S2a3). Accordingly there are 12 non-palindromic codons among the branch nodes in the positions #2, #5, #6, #10, #11, #12, #16, #20, #21, #23, #24 and #25. Due to the bijection between *Route* 1 and *Route* 3 in the sense of reverse relationship (Fig 1A, 2A), the complementary single RNA strands obtained in the two routes are same to each other (Fig 2A). Thus, there are totally 4 + (12 − 4) × 2 = 20 pairs (4 palindromic codons, and the 12 non-palindromic codons minus 4 identities between *Route* 1 and *Route* 3) of complementary single RNA strands, which combine into 20 *tRNA*s respectively (Fig S2a1, S2a3).

The anticodons of tRNAs such obtained may originate from either the triplet code in *Y*_*t*_ or the triplet code in *R*_*t*_ of the 20 possible pairs of triplet codes in the branch nodes (Fig S2a3). An assignment scheme can be obtained so that 20 anticodons chosen from the 20 pairs of possible triplet codes can be assigned to the 20 canonical amino acids respectively (Fig S2a2), which is as follows in details (Fig 2A, S2a2, S2a3). Note that these tRNAs are named after the series number of the recruitment order of the amino acids in the roadmap. There are 6 tRNAs (*t*1:1*Gly*, *t*3:3*Glu*, *t*7:7*S er*, *t*10:10*Arg*, *t*17:17*Phe*, *t*20:20*Lys*) from *Route* 0, 2 tRNAs (*t*4:4*Asp*, *t*19:19*Asn*) from *Route* 1, 6 tRNAs (*t*2:2*Ala*, *t*9:9*T hr*, *t*11:11*Cys*, *t*13:13*His*, *t*16:16*Met*, *t*18:18*T yr*) from *Route* 2, and 6 tRNAs (*t*5:5*Val*, *t*6:6*Pro*, *t*8:8*Leu*, *t*12:12*T r p*, *t*14:14*Gln*, *t*15:15*Ile*) from *Route* 3. The anticodons of *t*4 and *t*5 originated respectively from *Y*_*t*_5 and *R*_*t*_20 of the same pairs *Y*_*t*_5*R*_*t*_5 or *Y*_*t*_20*R*_*t*_20 (Fig 2A). The tRNA *t*19 originated from the combination of *Y*_*t*_30 and *R*_*t*_30, whose anticodon *AUU* came from a transition from *C* to *U* (Fig 2A, S2a2).

There are 20 canonical amino acids encoded by the degenerate codons. The above assignment scheme has provided an argument to account for the number of the canonical amino acids (Fig 2A, S2a2, S2a3). It will be explained, in Subsection 2.3, that not only the 64 codons result from the 64 permutations of the 4 bases, but also the 20 canonical amino acids relate to the 20 combinations of the 4 bases (Fig S3d1). The genomic codon distributions can be divided into 20 classes according to the 20 combinations of the 4 bases (Fig S3d1). Given the base compositions *p*(*i*), *i* = *G*, *C*, *A*, *T*, the products *p*(*i*) * *p*(*j*) * *p*(*k*), *i*, *j*, *k* = *G*, *C*, *A*, *T* are equal for each of the 20 combinations. The 20 combinations of the 4 bases can be divided into 4 groups < *G* >, < *C* >, < *A* >, < *T* > (Fig S3d1). *Hierarchy* 1 and *Hierarchy* 2 corresponds < *G* > and < *C* >; *Hierarchy* 3 and *Hierarchy* 4 corresponds to < *A* > and < *T* >. Their positions in the roadmap are

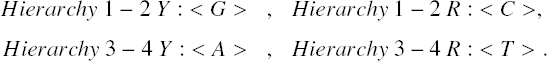

*Route* 0 and *Route* 1 − 3 are distinguished in each of the four groups. Take < *G* > for example, < *G*, *G*, *G* > an < *G*, *G*, *A* > belong to *Route* 0, while < *G*, *G*, *C* >, < *G*, *G*, *T* > and < *G*, *C*, *A* > belong to *Route* 1 − 3. The 20 tRNAs transfer 20 amino acids, the number of which is equal to the number of the 20 classes of genomic codon distributions. This is helpful to improve the translation efficiency, because it is easier for a tRNA to recognise a certain class of genomic codon distributions with the same codon interval distance.

The wobble pairing rules can be explained by the theory on the origin of tRNAs. The transition from *C* to *T* occurred in the position #6, which resulted in the wobble pairing rule *G* : *U or C* (Fig 2A). Taking *y*2*r*2 as a template, *y*_*t*_2 with *GCC* is formed by the triplex base paring, while *r*_*t*_2 with *GGC* and 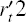 with *GGU* are formed, where the transition from *C* to *U* occurred in the formation of 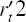. The complementary strands *y*_*t*_2 and 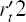 combine into a tRNA with anticodon *GCC*, where *G* at the first position of the anticodon of the tRNA is paired with *U* at the third position of the triple code of an additional single strand 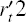. It implies that the wobble pairing rule *G* : *U* had been established as early as the end of the initiation stage of the roadmap. The transition from *C* to *T* occurred in the position #12, which resulted in the wobble pairing rule *U* : *G or A* (Fig 2A). Taking *y*10*r*10 as a template, *y*_*t*_10 with *CCG* is formed by the triplex base paring, and *r*_*t*_10 with *CGG* and 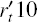 with *UGG* are also formed, where the transition from *C* to *U* occurred in the formation of 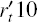. The complementary strands *y*_*t*_10 and 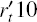 combine into a tRNA with anticodon *UGG*, where *U* at the first position of the anticodon of the tRNA is paired with *G* at the third position of the triple code of an additional single strand *y*_*t*_10. The above explanation of the wobble pairing rules by tRNA mutations is supported by the observations of nonsense suppressor. For instance, the wobble pairing rule *C* : *A* for a *UGA* suppressor can be established by a transition from *G* to *A* at the 24 − *th* position of *tRNA*^*Trp*^. The wobble pairing rules *G* : *U or C* and *U* : *G or A* had been established early in the evolution of the genetic code (Fig 2A), which continued to flourish so as to make full use of the tRNAs in short supply.

The 20 anticodons in the tRNAs from *t*1 to *t*20 correspond to the 20 amino acids, respectively. Hence, the corresponding 20 codons can be regarded as the minimum subset of the genetic code that encodes all the amino acids (Fig S2a2). The minimum subset consists of *GGG*, *GCG*, *GAG*, *GAC*, *GUG*, *CCG*, *UCC*, *CU A*, *ACG*, *AGG*, *UGC*, *UGG*, *CAC*, *CAG*, *AUC*, *AUG*, *UUC*, *U AC*, *AAU*, *AAG*. For each amino acid, its corresponding codon in the minimum subset generally recruited earlier than the other corresponding codons out of the minimum subset (Fig S2c2). The codons in the minimum subset whose first base are purines generally correspond to the anticodons that come from the strands *Y*_*t*_, except for *GUC* and *AUC* (Fig S2a2); those whose first base are pyrimidines generally correspond to anticodons that come from the strands *R*_*t*_, except for *CAG* and *UGG* (Fig S2a2). The codons in the minimum subset are generally situated in the branch nodes, except for *AAG*, *UCC*, *AAU*, *AUG* and *UGC* (Fig 2A, S2a2). The numbers of codons in the minimum subset in *Route* 0 − 3 are 6, 4, 6, 4 respectively (Fig 2A, S2a2). The minimum subset can be extended through wobble pairings. Additional tRNAs can be combined according to the roadmap so as to recognise all the codons.

### 1.5 Codon degeneracy

The four routes of the roadmap (Fig 1A) can be represented by four cubes (Fig 2C). The vertices of the cubes represent codon pairs, while the edges of the cubes represent substitution relationships (Fig 2C). Cubes are convenient to demonstrate pair connections and route dualities, by which it is more intuitive to study the codon degeneracy. The degeneracies 6, 4, 3, 2 or 1 for the 20 amino acids can be explained one by one according to pair connections and route dualities in the roadmap (Fig 1A, 2C). The degeneracy 2 mainly results from pair connections. The degeneracy 4 or 6 mainly result from the expansion of the genetic code from the initial subset by route dualities pertaining to *S er*, *Leu*, *Ala*, *Val*, *Pro* and *T hr* (Fig 2B, 2C).

The degeneracy 6 for *S er*, *Leu* and *Arg* can be explained by pair connections and route dualities (Fig 1A, 2C), where *S er* and *Leu* belong to the initial subset and *Arg* was recruited immediately after the initial subset. And all of them have appeared in *Route* 0. The 6 codons of *S er* satisfy both the pair connection #8-*S er* -#17 and the route duality

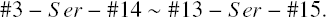

The 6 codons of *Leu* satisfy both the pair connection #28 − *Leu* − #31 and the route duality

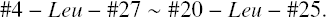

The 6 codons of *Arg* satisfy both the pair connection #7 − *Arg* − #16 and the route duality

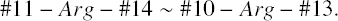

The degeneracy 4 for *Gly*, *Ala*, *Val*, *Pro* and *T hr* can be explained by route dualities (Fig 1A, 2C). All of them belong to the initial subset. The degeneracy 4 for *Gly* satisfy the route duality:

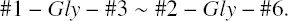

The degeneracy 4 for *Ala* satisfy the route duality:

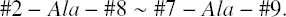

The degeneracy 4 for *Val* satisfy the route duality:

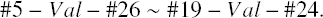

The degeneracy 4 for *Pro* satisfy the route duality:

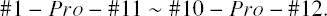

The degeneracy 4 for *T hr* satisfy the route duality:

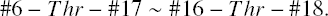

The degeneracy 2 for *Glu*, *Asp*, *Cys*, *His*, *Gln*, *Phe*, *T yr*, *Asn* and *Lys* can be explained by pair connections (Fig 1A, 2C). They satisfy the following pair connections respectively: #4 − *Glu* − #23, #5 − *Asp* − #21, #9 − *Cys* − #18, #19 − *His* − #22, #20 − *Gln* − #28, #23 − *Phe* − #32, #24 − *T yr* − #29, #26 − *Asn* − #30, #27 − *Lys* − #32. The degeneracy 3 for *Ile* and the degeneracy 1 for *Met* satisfies the route duality (Fig 1A, 2C)

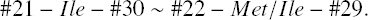

The degeneracy 1 for *T r p* satisfies the pair connection for nonstandard genetic code #12 − *T r p*/*stop*(*T r p*) -#15. This pair connection includes a stop codon; the other stop codons satisfy the pair connection: #25 -*stop* − #31 (Fig 1A, 2C).

It is convenient to illustrate the symmetry of the roadmap by the cubes (Fig 2C). Some correspondences of codon pairs among routes are as follows (Fig 2C):

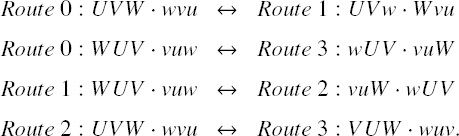

The feature of pair connections in the roadmap can be explained by the wobble pairings. The pair connections in the routes: *Route* 0 *R* strand, *Route* 2 *R* strand and *Route* 3 can be explained by the wobble pairing rule *U* : *G*, *A* (Fig 2C); The pair connections in the routes: *Route* 0 *Y* strand, *Route* 2 *Y* strand and *Route* 1 can be explained by the wobble pairing rule *G* : *U*, *C* (Fig 2C). By and large, there is a symmetry between *Route* 0, 1 and *Route* 3, 2 in the roadmap (Fig 1A, 2C). Considering the expansion of the genetic code from the initial subset, the pair connections pertaining to *Pro*, *S er* and *Leu* in *Route* 0 *Y* strand are dual to the pair connections in *Route* 3 *Y* strand, respectively (Fig 2B, 2C); the pair connections pertaining to *Ala*, *T hr* and *Val* in *Route* 1 *Y* strand are dual to the pair connections in *Route* 2 *R* strand, respectively (Fig 2B, 2C). The codon pairs can be divided into groups by the 6 facets of the cubes for the routes respectively: *Facet u* and *Facet d* by up and down; *Facet w* and *Facet e* by west (left) and east (right); *Facet n* and *Facet s* by north (back) and south (front) (Fig 2C). According to the route dualities, *Route* 0 and *Facet u* are dual to *Route* 3 and *Facet s*; *Route* 0 and *Facet d* are dual to *Route* 3 and *Facet n*; *Route* 1 and *Facet s* are dual to *Route* 2 and *Facet u*; *Route* 1 and *Facet n* are dual to *Route* 2 and *Facet d* (Fig 2C).

The roadmap not only explain the codon degeneracy for the 20 amino acids, but also discriminate the 5 biosynthetic families of the amino acids. A *GCAU* genetic code table can be obtained by changing the order *U*, *C*, *A*, *G* in the standard genetic code table to a new order *G*, *C*, *A*, *U*, where the third base are sorted by the order *G*, *A*, *C*, *U* in considering the wobble pairings (Fig S2c2). The biosynthetic families of *Glu*, *Asp*, *Val*, *S er* and *Phe* situate in the continuous regions in the *GCAU* genetic code table, respectively (Fig S2c2). The codons as well as amino acids, in the recruitment orders, expand roughly from up to down, from left to right in the *GCAU* genetic code table (Fig S2c2). So the evolution of the genetic code can be illustrated better in the *GCAU* genetic code table than in the standard genetic code table. The clusters of the biosynthetic families can be explained by the roadmap. The codons are classified by the *R* strands and *Y* strands of the four routes as follows:

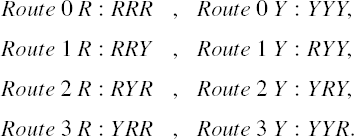

In such a classification, the *GCAU* genetic code table can be divided into 8 × 4 blocks (Fig S2c1), which consequently influences the codon degeneracy and the clustering of biosynthetic families (Fig S2c2). The region of *Route* 0 *R GRR* and *Route* 1 *GNY* (*N* denotes purine or pyrimidine) relate to the biosynthetic relationships among *Gly*, *Ala*, *Glu*, *Asp*, *Val* (Fig S2c1, Fig S2c2). The region of *Route* 3 *CNR* explains the alignment of *Arg*, *Pro*, *Gln* that belong to the biosynthetic family of *Glu* (Fig S2c1, Fig S2c2). The region of *Route* 1 *ANY* explains the alignment of *T hr*, *Asn*, *Ile* that belong to the biosynthetic family of *Asp* (Fig S2c1, Fig S2c2). The region of *Route* 1 *RNY* relates to the biosynthetic relationships among *Asp* and *T hr*, *Asn* (Fig S2c1, Fig S2c2). The region of *Route* 3 *R URR* explains the correlations of positions among the 3 stop codons (Fig S2c1, Fig S2c2). Moreover, the region of *Route* 0 *R RRR* can explain the biosynthetic relationships from *Glu* to *Lys*; and *ARG* was assigned to *Arg* of the biosynthetic family of *Glu* (Fig S2c1, Fig S2c2). The region of *Route* 2 *R GYR* explains the biosynthetic relationships from *Ala* to *Val* (Fig S2c1, Fig S2c2). The region of *Route* 3 *Y YUR* explains the degeneracy of *Leu* (Fig S2c1, Fig S2c2).

The roadmap provides a framework beyond the explanation of the evolution of the genetic code. According to the roadmap, the genetic code, tRNA, amino acid and aminoacyl-tRNA synthetase evolved together. Except the certain three positions of the codon, other positions also evolve along the roadmap (Fig S3a1). The evolution of the triplex nucleic acids along the roadmap might generate the corresponding sequence encoding the aminoacyl-tRNA synthetase. The classification of the aminoacyl-tRNA synthetases distributed regularly in the roadmap. The aminoacyl-tRNA synthetases are divided into two classes *Class II* and *Class I*, and accordingly subclasses *IIA*, *IIB*, *IIC*; *IA*, *IB*, *IC* [10]. According to the recruitment order of the amino acids, the *Class II* aminoacyl-tRNA synthetases originated earlier than the *Class I* aminoacyl-tRNA synthetases (Fig 2A, 2C). In the initial subset, #1, #2, #3, #5 *Y*, #6 are *Class II* aminoacyl-tRNA synthetases; *Gly* − *tRNA* appeared at #1 as the first *Class II* aminoacyl-tRNA synthetase (Fig 2C). #4, #5 *R* are *Class I* aminoacyl-tRNA synthetases; *Glu*-*tRNA* appeared at #4 as the first *Class I* aminoacyl-tRNA synthetase (Fig 2C). Majority of the *Route* 0, 1 are *Class II* aminoacyl-tRNA synthetases; majority of the *Route* 2, 3 are *Class I* aminoacyl-tRNA synthetases (Fig 2C). The exceptions mostly concern with the route duality between *Route* 0 and *Route* 3 pertaining to *S er*, *Leu*, *Pro*, and route duality between *Route* 1 and *Route* 2 pertaining to *Ala*, *Val*, *T hr* (Fig 2C). The early *Class II* aminoacyl-tRNA synthetases recognize the variable arm of the tRNA. According to the explanation of the origin of the tRNA, the variable arm is complementary with the anticodon arm (Fig S2a3). Paracodons may be around the variable arm. The anticodons of the degeneracies 4 or 6 are *GGN*, *GCN*, *GU N*, *CGN*, *CCN*, *CU N*, *ACN* and *UCN*. The aminoacyl-tRNA synthetases in these codon boxes are *IIA* or *IA* (Fig S2c3). The aminoacyl-tRNA synthetases with anticodons of the small degeneracies are *IIB*, *IIC*, *IB*, *IC* (Fig S2c3). The aminoacyl-tRNA synthetases can also be divided into major groove (*M*) and minor groove (*m*) [11]. They distribute regularly in the roadmap (Fig 2C):

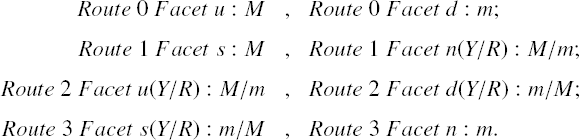

The aminoacyl-tRNA synthetasis and the tRNA determines the pair connections and route dualities in the roadmap, which resulted in the codon degeneracy for amino acids and the 5 biosynthetic families of amino acids (Fig 2C).

### 1.6 Origin of homochirality of life

Homochirality is an essential feature of life. In Part I of this paper, the origin of the homochirality of life is explained by chiral symmetry breaking between the left-handed roadmap and the right-handed roadmap (Fig 2D), which helps to explain the origin of the living system from the non-living system. This is analogous to (but not trivially related to) the spontaneous symmetry breaking in particle physics [12] [13], which helps to explain the generation of masses from massless fields. The time for the existence of life on the Earth is in the same order of magnitude of the age of the universe, which implies that life is an intrinsic phenomenon in the universe that originates spontaneously at the macromolecular level. Homochirality safeguards the stability of the living system, which has ensured the life to last comparable with the age of the universe.

The first recruited amino acid *Gly* is not chiral (Fig 1A #1). The following 19 amino acids are chiral (Fig 1A #2). In the initiation stage of the roadmap, the nonchiral *Gly* helped to create the first pair connection #1 − *Gly* − #2, recruiting chiral *Ala* at #2 (Fig 1A). And the nonchiral *Gly* also helped to create the first route duality in the roadmap (Fig 1A):

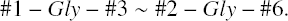

This route duality played a central role in the initiation stage; consequently the initial subset played a central role in the expansion stage (Fig 2B). The chirality was required at the beginning of the roadmap by the triplex DNA itself (Fig 1A, 1B). Even so, there was still a transition period from nonchirality to chirality, in consideration of the special role of nonchiral *Gly*.

The complex evolution from *Hierarchy* 1 to *Hierarchy* 4 was a recurrent and cyclic process, with the intermittent chemical reaction conditions in environment. There were two possible chiral roadmaps in the evolution of the genetic code as follows (Fig 2D):

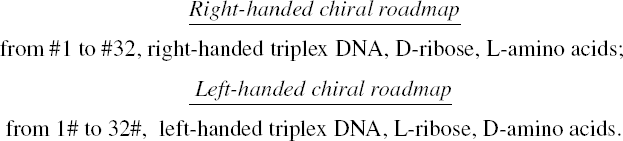

The four bases themselves are not chiral, which are shared in common by the two chiral roadmaps. The two chiralities were selected randomly in the winner-take-all competition. The translation efficiency improved greatly as soon as all the 64 codons were recruited into the roadmap. First completed chiral roadmap dominated and was able to decompose the uncompleted opposite chiral system. Hence, there is no longer the chance for the existence of the opposite chirality.

The origin of the homochirality of life can be simulated based on the roadmap. The competition between the right-handed chiral roadmap and the left-handed chiral roadmap can be simulated by a stochastic model. The selection of chirality of life was absolutely by chance with equal probability. The chiral roadmap evolves from #1 to #32 or from 1# to 32# by a constance substitution probability step by step (Fig S2d1). The struggle for shared resources, such as the bases, might existed between the two chiral roadmaps (Fig S2d1). The model can estimate how many steps the winner pulled away from the loser in the competition of the two chiral roadmaps. The results of the estimations are as follows: around 3 steps gap in the case without struggle for bases (Fig S2d1), and around 7 steps gap in the case with struggle for bases (Fig S2d1). In either case, the chiral roadmap of the loser had already finished the initiation stage and expansion stage according to the simulations. Namely, it was in the ending stage that the two chiral roadmaps were separated (Fig S2d1). The recruitment of the last two stop codons in the ending stage may promote the separation of the two chiral roadmaps (Fig 1A, S2d1).

It is a great expectation to achieve the homochirality of life based on the roadmap for the experimentalists. Artificial conditions are more operational and efficient than the environmental changes. Proper procedures and schedules are expected to considerably shorten the time to achieve the homochirality of life in laboratory. This is indeed a challenge.

### 1.7 Evidence for the roadmap

So far, a detailed description of the evolution of the genetic code has been proposed according to the roadmap theory. In this subsection, numerous theoretical and experimental evidences are listed concretely that will be explained here or later on.

The key experimental evidences in supporting of the roadmap are as follows. The roadmap itself is based on the experimental results on the relative stabilities of the base triplexes. The delicate roadmap has narrowly avoided the unstable base triplexes. The layout of the roadmap is, therefore, determined by the increasing stabilities of the base triplex along the roadmap (Fig S1a1). Genomic data provide the most successful and most straightforward experimental evidence for the roadmap, which will be explained in detail in Section 2. An evolutionary tree of codons is obtained based on the genomic codon distributions, where the four hierarchies of the roadmap are distinguished clearly (Fig 3A). This is a straightforward evidence to support the roadmap theory. Coincidentally, an evolutionary tree of species can also be obtained based on the same genomic codon distributions and the roadmap (Fig S3d8, Fig S3d9), which agrees with the tree of three domains. The successful explanation of the origin of the three domains also conversely supports the roadmap theory. Another strong and heuristic evidence for the roadmap is that the origin of the homochirality of life can be explained by the roadmap straightforwardly (Fig 2D). If the homochirality of life can be achieved by the roadmap in experiment, the roadmap will be conclusively confirmed. The origin of the genetic code, the origin of the homochirality of life and the origin of the three domains can be explained together in the roadmap picture (Fig 1, 2, 3). These three theories support one another.

Furthermore, the roadmap is supported by the following evidences. The roadmap predict the recruitment order of the 64 codons, the recruitment order of the 3 stop codons and the recruitment order of the 20 amino acids. Evaluation of the three recruitment orders may verify the roadmap theory.

The recruitment order of the 32 codon pairs can be obtained from #1 to #32 by the roadmap (Fig 2B, S2b1). The declining *GC* content indicates the evolution direction because of the substitutions from *G* to *A*, from *G* to *C* and from *C* to *T* in the roadmap (Fig S2b1). According to the roadmap, the total *GC* content and the position specific *GC* content for the 1*st*, 2*nd* and 3*rd* codon positions are calculated for each step from #1 to #32. Then the relationship between the total *GC* content and the position specific *GC* content is obtained (Fig S2b2). The 1*st* position *GC* content is higher than the 2*nd* position *GC* content. And the 3*rd* position *GC* content declined rapidly from the highest to the lowest in the evolution direction when the total *GC* content declines. The *GC* content variation in the simulation agrees with the observation (to compare Fig S2b2 with Figure 2 in [14] and Figure 5 in [15]), so the recruitment order obtained by the roadmap is reasonable. The recruitment order of the codons by the roadmap generally agree with the order in [4] [16] [17]. It should be noted that the roadmap itself was conceived firstly by studying the substitution relationships based on the recruitment order in [4], which was furthermore improved in accordance with the recruitment order of amino acids.

The recruitment order for the three stop codons are #15 *UGA*, #25 *U AG*, #31 *U AA* (Fig 1A, 2B), which results in the different variations of the stop codon usages. Along the evolution direction as the declining *GC* content, the usage of the first stop codon *UGA* decreases; the usage of the second stop codon *U AG* remains almost constantly; the usage of the third stop codon *U AA* increases (Figure 1 in [18]). The observations of the variations of stop codon usages can be simulated (to compare Fig S2b3 with Figure 1 in [18]) according to the recruitment order of the codon pairs (Fig 2B, S2b1) and the variation range of the stop codon usages. Especially, the detailed features in observation can be simulated that the usage of *UGA* jump downwards greatly; *U AA*, upwards greatly, around half *GC* content (to compare Fig S2b3 with Figure 1 in [18]).

The recruitment order of the 20 amino acids from *No*.1 to *No*.20 can be obtained by the roadmap (Fig 2B, S2b1), which meets the basic requirement that Phase *I* amino acids appeared earlier than the Phase *II* amino acids [19] [20]. The species with complete genome sequences are sorted by the order *R*_10/10_ according to their amino acid frequencies, where the order *R*_10/10_ is defined as the ratio of the average amino acid frequencies for the last 10 amino acids to that for the first 10 amino acids [21]. Along the evolutionary direction indicated by the increasing *R*_10/10_, the amino acid frequencies vary in different monotonous manners for the 20 amino acids respectively (Fig S2b4). For the early amino acids *Gly*, *Ala*, *Asp*, *Val*, *Pro*, the amino acid frequencies tend to decrease greatly, except for *Glu* to increase slightly (Fig S2b4); for the midterm amino acids *S er*, *Leu*, *T hr*, *Cys*, *T r p*, *His*, *Gln*, the amino acid frequencies tend to vary slightly, except for *Arg* to decrease greatly (Fig S2b4); for the late amino acids *Ile*, *Phe*, *T yr*, *Asn*, *Lys*, the amino acid frequencies tend to increase greatly, except for *Met* to increase slightly (Fig S2b4). In the recruitment order from *No*.1 to *No*.20, the variation trends of the amino acid frequencies increase in general (Fig S2b5); namely, the later the amino acids recruited, the more greatly the amino acid frequencies tend to increase (Fig S2b4, Fig S2b5). The recruitment order of the amino acids from *No*.1 to *No*.20 is supported not only by the previous roadmap theory but also by this pattern of amino acid frequencies based on genomic data, so it help to resolve the disputes on such an issue in the literatures [22].

## 2 Origin of the three domains

### 2.1 Genomic codon distributions reveal the relationship between the evolution of the genetic code and the tree of life

In the previous section, the roadmap theory was proposed, as a hypothesis, to explain the evolution of the genetic code and the origin of the homochirality of life (Fig 1, 2). In this section, more experimental evidences are provided to enhance and reinforce the roadmap theory (Fig 3). Especially, the four hierarchies in the roadmap are directly distinguished in a tree of codons obtained by studying the genomic codon distributions (Fig 3A). And the origin of the three domains of life can be explained based on the roadmap and the genomic data (Fig 3D). In short, the biodiversity originated in the evolution of the genetic code, and has flourished by the environmental changes.

The viewpoint that the tree of life was rooted in the recruitment of codons can be proved by analysing the complete genome sequences. The genomic codon distribution is a powerful method to study the statistical properties of the complete genome sequences, which can reveal the profound relationship between the primordial genetic code evolution and the contemporary tree of life (Fig 3A, S3a5). The genomic codon distribution for a species with complete genome sequence is defined as follows (Fig S3a3): first, mark all the positions for each of the 64 codons in its complete genome sequence respectively; second, calculate all the codon intervals between the adjacent marked positions for each of the 64 codons respectively, and obtain the distributions of codon intervals according to the 64 sets of codon intervals respectively (some codon intervals count more, and the others less), where a cutoff 1000 is used when counting the number of common intervals; third, obtain the genomic codon distribution by normalising the 64 distributions of codon intervals. Normalisation eliminates the direct impact of a wide range variation of genome sizes.

Both the tree of codons (Fig 3A) and the tree of species (Fig S3a5) are obtained according to the same set of correlation coefficients among the genomic codon distributions. 952 complete genomes in GeneBank (up to 2009) were used in the calculations. There are 64 genomic codon distributions for each of the 952 species. On one hand, the average correlation coefficient matrix for codons is obtained by averaging these 64 × 64 correlation coefficient matrices for the 952 species; hence the tree of codons is obtained based on the 64 × 64 distance matrix that equals to half of one minus the average correlation coefficient matrix for codons (Fig 3A); on the other hand, the average correlation coefficient matrix for species is obtained by averaging these 952 × 952 correlation coefficient matrices for the 64 codons, hence the tree of species is obtained based on the 952 × 952 distance matrix that equals to half of one minus the average correlation coefficient matrix for species (Fig S3a5). The codons from the four hierarchies *Hierarchy* 1−4 in the roadmap gather together in the tree of codons respectively, where the *Hierarchy* 1 and *Hierarchy* 2 situated in one branch, and *Hierarchy* 3 and *Hierarchy* 4 in another branch (Fig 3A). This is a direct evidence to support the roadmap (Fig 1A, 3A). According to the NCBI taxonomy, the tree of species roughly presents the phylogeny of species (Fig S3a5). Especially, the tree of the eukaryotes with complete genomes is reasonable to some extent (Fig S3a6). In Subsection 2.3, an improved tree of species (Fig S3a8, S3a10) will be obtained by dimension reduction for the 64 × 952 genomic codon distributions according to the roadmap (Fig S3d2, 3D).

The validity of the method of genomic codon distribution can be explained in the roadmap picture. Whereas the more concerns were given to the three-base-length segments of the triplex DNA in the previous section, base substitutions can occur in any places through the whole length of the triplex DNA (Fig S3a1). Starting from the initial *Poly C* · *poly G* * *poly G*, several codons in different regions of the triplex DNA can evolve independently along the roadmap (Fig S3a1). In this way, the triplex DNA evolves gradually from the initial identical sequences to the final non-identical sequences (Fig S3a1). The identical sequences tend to form a triplex DNA, but it becomes more and more difficult for the non-identical sequences to form a triplex DNA. So there was a transition from triplex DNA to double DNA after the genetic code evolution. The evolution of the whole sequence may result in varieties of distributions of codons in the genomes (e.g., *T AA* in the last strand in Fig S3a1). The genomic codon distributions, therefore, resulted from the evolution of the genome sequences. Thus, the biodiversity at the species level actually originated at the sequence level. The genomic codon distributions keep the essential features of species at the sequence level. The genomic codon distributions were reluctant to change, in the context of normalisation, during the evolution of genome sizes mainly promoted by large scale duplications (Fig S3a4). In the long history of evolution, even if genome sizes have been doubled for many times, the essential features in the ancestors’ sequences can be inherited generation by generation due to the relative invariant of the genomic codon distributions. As a result, the ancient information on the evolution of the genetic code and the origin of the three domains can be dug out from the complete genome sequences of contemporary species. It should be emphasised that the sequences evolved based on the roadmap were by no mean random, which may account for the linguistics in the biological sequences.

According to the roadmap, the initial subset took shape in the initiation stage of the evolution of the genetic code (Fig 1A), where the *GC* content of the initial subset was 13/18; leaf nodes appeared in the ending stage (Fig 1A), where the *GC* content of the leaf nodes decreased to 16/48 (Fig S3a1). The variation range of *GC* contents from 16/48 to 13/18 obtained by the roadmap generally agrees with the observations of *GC* contents for contemporary species (Fig S3a1, Fig S3a2). Along with the evolution of the whole sequences (Fig S3a1), the shorter triplex DNAs only accommodate a proper subset of the genetic code; the longer triplex DNAs accommodate almost all the codons. Such a difference will be considered in Subsection 2.2 to explain the different features of genomic codon distributions between Eukaryotes and Bacteria etc. Note that Eukaryotic genomes mainly consists of non-coding DNA, the tree of eukaryotes based on their genomic codon distributions is, therefore, mainly depended on the non-coding DNA (Fig S3a6, S3d10). This indicates that the non-coding DNA has played a substantial role in the evolution of eukaryotes.

### 2.2 Features of genomic codon distributions: observations and simulations

The features of genomic codon distributions are different among Virus, Bacteria, Archaea and Eukarya (Fig 3B, S3b2), which are also called the (3 + 1, including Virus) domains for convenience. The genomic codon distribution for a domain is obtained by averaging the genomic codon distributions of all the species in this domain (only considering those with complete genome sequences, in practice) (Fig 3B). The feature of genomic codon distribution for a species is similar to that of the corresponding domain (Fig 3B, S3b2). The method based on the genomic codon distribution apply not only for cellular life but also for viruses. Explanation of the features of the genomic codon distributions (Fig 3B, S3b3) is helpful to understand the origin of the three domains and to clarify the status of virus.

The features of genomic codon distributions for the domains can be described respectively from two aspects: first, the periodicity and the amplitudes of the genomic codon distributions; second, the order of the average distribution heights for 64 codons. (*i*) The features of genomic codon distribution for Virus include declining three-base-periodic fluctuations, high amplitudes (Fig 3B, S3b2). In addition, the order of the average distribution heights is: *Hierarchy* 1, highest; followed by *Hierarchy* 2 and *Hierarchy* 3; and *Hierarchy* 4, lowest (Fig S3c1). (*ii*) The features of genomic codon distribution for Bacteria include declining three-base-periodic fluctuations, medium amplitudes (Fig 3B, S3b2). In addition, the order of the average distribution heights is: *Hierarchy* 1, highest; followed by *Hierarchy* 2, *Hierarchy* 3 and *Hierarchy* 4 (Fig S3c1). (*iii*) The features of genomic codon distribution for Archaea include declining three-base-periodic fluctuations, medium amplitudes (Fig 3B, S3b2). In addition, the order of the average distribution heights is roughly opposite to that of Bacteria: *Hierarchy* 4, highest; followed by *Hierarchy* 3 and *Hierarchy* 2; and *Hierarchy* 1, lowest (Fig S3c1). Comparing Archaea with Bacteria, the dwarf 2nd peak at 6-base interval in the genomic codon distribution is even lower than the 3rd peak at 9-base interval (Fig 3B, S3b2). This detailed feature is different from Bacteria. For Bacteria, the 2nd peak is higher than the 3rd peak for codons in *Hierarchy* 1 and the heights are almost the same for the 2nd and 3rd peaks for other codons (Fig 3B, S3b2). (*iv*) The features of genomic codon distribution for Eukarya are more complicated than for other domains, which include declining distributions, indistinct multi-base-periodic fluctuations, ups and downs of peaks, or overlapping peaks for some codons (Fig 3B, S3b2). In addition, the order of the average distribution heights is similar to that of Archaea: *Hierarchy* 4, highest; followed by *Hierarchy* 3 and *Hierarchy* 2; and *Hierarchy* 1, lowest (Fig S3c1). The features of genomic codon distribution for Archaea put together the features of both Archaea and Eukarya (Fig 3B, S3b2). Crenarchaeota and Euryarchaeota are similar in genome size distributions (Fig 3B, S3b1). It is showed that the tree of the three domains is more reasonable than the eocyte tree of life, according to the features of genomic codon distributions (Fig 3B, S3b1).

The origin order for the domains can be obtained according to the linear regression plot whose abscissa and ordinate represent the recruitment order of codon pairs and the order of average distribution heights respectively (Fig 3C, S3c1). The greater the slope of the regression line for a domain is, the more the late recruited codons are. So, the order of the slopes of the regression lines represents the origin order of the domains: 1 Virus, 2 Bacteria, 3 Archaea, 4 Eukarya (Fig 3C). The slope for Virus is negative and least (Fig 3C), which means that majority of codons in the sequences recruited early. The virus originated earlier than the three domains. The slope for Bacteria is negative (Fig 3C). The proportion for *Hierarchy* 1 is high; that for *Hierarchy* 2 − 4, lower. The slope for Archaea is positive (Fig 3C), which means that majority of codons in the sequences recruited late. The slope for Eukarya is positive and greatest (Fig 3C), which also means that majority of codons in the sequences recruited late. The Eukarya originated latest. Archaea originated between Bacteria and Eukarya (Fig 3C). Limited data indicate that the slope for Crenarchaeota is less than the slope for Euryarchaeota (Fig 3C), which suggests that Crenarchaeota may originate early than Euryarchaeota. This origin order of the domains is independent from the base compositions for the domains, which are different from the order of the domains by *GC* content (Fig S3a2).

The features of the genomic codon distributions for the domains can be simulated by a stochastic model. The features of declining three-base-periodic fluctuations can be simulated by the model (Fig 3B, S3b3). In the model, the bases *G*, *C*, *A*, *T* are given in certain base compositions, from which three bases are chosen statistically to make up a codon. Many a codon concatenates together to form a genome sequence in simulation. Hence, the genomic codon distribution is obtained by calculating the simulated genome sequence. When the length of the triplex DNA increased in the evolution along the roadmap (Fig S3a1), more and more codons are accommodated into the triplex DNA. The feature of declining three-base-periodic fluctuations, therefore, is achieved in the simulations. Let *n*_0_ denote the number of codons accommodated in the triplex DNA. *n*_0_ codons are generated statistically to concatenate the simulated genome in several cycles. The genomic codon distribution thus obtained has the feature of declining three-base-periodic fluctuations. When *n*_0_ is small (e.g., around 16), the amplitude of the three-base-periodic fluctuations is great. The greater *n*_0_ is, the less the amplitude of the three-base-periodic fluctuations is. When *n*_0_ is 32 (namely all codons participated in the concatenation), the three-base-periodic fluctuations vanish. The amplitude of the three-base-periodic fluctuations decreases from Virus to Bacteria, then Archaea, finally Eukarya (Fig 3B, S3b3), which results from the increasing number of codons accommodated in the triplex DNA at different stages in the evolution of the genetic code. The complicated features of genomic codon distributions for eukaryotes can also be simulated by the model. The features of ups and downs of peaks for eukaryotes can be simulated by concatenating different set of codons that are generated from different subsets of the genetic code (Fig S3b3). The alternative “high peak - low peak” feature for some codons and the low amplitudes of the genomic codon distributions for other codons agree with the observations for true eukaryotes (Fig 3B, S3b2, S3b3).

The domains differ in the orders of the average distribution heights, which can also be simulated by the statistical model (Fig S3c2). The *GC* content tends to decrease along the evolutionary direction according to the roadmap (Fig 1A, S2b2), which accounts for the above different orders among the domains. Different base compositions for *G*, *C*, *A*, *T* are assigned for Virus, Bacteria, Archaea and Eukarya in the simulations, in consideration of the decrement of *GC* content effected by the origin order of the domains. The simulations of genomic codon distributions thus obtained for the domains (Fig S3b3) differ in the corresponding orders of the average distribution heights (Fig S3c2). If high *GC* contents are assigned to Virus or Bacteria in the simulation, the average distribution heights from high to low are as follows: *Hierarchy* 1, *Hierarchy* 2, *Hierarchy* 3, and *Hierarchy* 4 (Fig S3c2). If low *GC* contents are assigned to Archaea or Eukarya in the simulation, the average distribution heights from high to low are as follows: *Hierarchy* 4, *Hierarchy* 3, *Hierarchy* 2, and *Hierarchy* 1 (Fig S3c2). The simulations generally agree with the observations as to the order of the average distribution heights (Fig S3c1, Fig S3c2).

Incidentally, the validity of the composition vector tree (a method to infer the phylogeny from genome sequences) in the literatures [23] [24] can be explained based on the genomic codon distribution in this paper (Fig S3a3). The average distribution heights for the 64 codons are proportional to the numbers of codons in the genomes. The summations of the elements of the unnormalized genomic codon distributions for a species represent the numbers of the 64 codons in the genome. So a set of average distribution heights is equivalent to the *K* = 3 composition vector for this species. For example, the summations of the elements of the unnormalized genomic codon distributions of *GCA* and *GCA* are 32 and 38 respectively, which means that there are 33 *GCA* and 39 *T AT* in the genome (Fig S3a3). Hence, the *K* = 3 composition vector for this species is obtained, where the elements corresponding to *GCA* and *T AT* are 33 and 39 respectively (Fig S3a3). The *K* = 3 composition vector (Fig S3a3) in fact corresponds to the order of the average distribution heights (Fig S3c1). It has been emphasised that the genomic codon distributions are essential features of species at the sequence level. The composition vector, as a deformed method of the genomic codon distribution, is certainly valid to infer the phylogeny.

### 2.3 The evolution space and the tree of life

According to the roadmap theory and the features of genomic codon distributions, an evolution space (Fig 3D) has been constructed to explain the major and minor branches of the tree of life (Fig S3d8, Fig S3d9). The evolution space is 3-dimensional, the coordinates of which are fluctuation, route bias and hierarchy bias (Fig 3D). (*i*) The fluctuation is defined by the total summation of the absolute value of the neighbouring differences of the fluctuations in the 64 genomic codon distributions for a species. (*ii*) The route bias is defined by the difference of the average genomic codon distributions between *Route* 0 and *Route* 1 − 3 (Fig S3d2). (*iii*) The hierarchy bias is defined by the difference of the average genomic codon distributions between *Hierarchy* 1−2 and *Hierarchy* 3 − 4 (Fig S3d2).

It should be pointed out that the domains cannot easily be distinguished into clusters by the bias similarly defined on difference of average genomic codon distributions between arbitrarily divided two groups of codons. So, the definitions of route bias and hierarchy bias themselves reveal the essential features of the genetic code. After a lot of groping around, it was realised that only proper groupings of codons by the roadmap are able to distinguish the domains (Fig 3D). When defining route bias and hierarchy bias under the major premise of clusterings of the three domains, the recruitment order of codons and the roadmap symmetry are also in consideration (Fig S3d1, Fig S3d2). The genome size distributions can be divided into 20 groups according to the 20 combinations of the 4 bases (Fig S3d1). The number 20 of combinations is equal to the number of canonical amino acids. And the 20 combinations generally correspond to the 20 tRNA in Subsection 1.4, among which 14 combinations correspond to certain tRNAs respectively (Fig S3d1, Fig S2a3). So, the classifications of genome size distributions are of biological significance.

The three coordinates of the evolution space are obtained by calculations for each species, thus a species corresponds to a point in the evolution space (Fig 3D). The species gather together in different clusters for Virus, Bacteria, Archaea and Eukarya (Fig 3D, S3d3, S3d4, S3d5). The cluster distributions of taxa of lower rank are also observed in detail in the evolution space (Fig 3D, S3d3, S3d4, S3d5). The evolutionary relationships among species are thereby illustrated in the evolution space, which reveals the close relation between the origin of biodiversity and the origin of the genetic code. According to the fluctuations of genomic codon distributions, the order of domains is as follows: first Virus, then Bacteria and Archaea, and last Eukarya (Fig S3d4, Fig S3d5). Bacteria and Archaea are distinguished roughly by route bias (Fig S3d3, Fig S3d5), and they can be distinguished better by both hierarchy bias and route bias (Fig 3D, S3d3). Diversification of the domains is, therefore, due to route bias and hierarchy bias at the sequence level. In addition, the leaf node bias can be defined by the difference between the average genomic codon distributions of the leaf nodes in *Route* 0 − 1 and that of the leaf nodes in *Route* 2 − 3 (Fig 1A, S3d2). Hence, Crenarchaeota and Euryarchaeota are distinguished by leaf node bias (Fig S3d6, Fig S3d7). Diversification of Crenarchaeota and Euryarchaeota is, therefore, due to leaf node bias at the sequence level. The cluster distributions of the domains in the evolution space shows that the biodiversity originated in the evolution of the genetic code at the sequence level. And the effects from the earlier stage of the evolution of the genetic code contribute more to the tree of life.

A tree of life (Fig S3d8) can be obtained based on the distances among species in the evolution space (Fig 3D). The three coordinates of the evolution space need to be equally weighted by non-dimensionalization of the three coordinates through dividing by their standard deviations respectively first. Then a distance matrix is obtained by calculating the Euclidean distances among species in the non-dimensionalized evolution space. Thus, the tree of life is constructed by PHYLIP software based on the distance matrix (Fig S3d8). Note that all the phylogenetic trees in this paper are constructed by PHYLIP software using UPGMA method [25]. The tree of life based on the evolution space is reasonable to describe the evolutionary relationships among species. Take the eukaryotes for example. Such a tree provides reasonable inclusion relationships among vertebrates, mammals, primates, where chimpanzee is nearest to human beings. And yeast indicates its root (Fig S3d10).

A tree of taxa can also be obtained according to the average distances of species among Virus, Bacteria, Crenarchaeota, Euryarchaeota and Eukarya in the non-dimensionalized evolution space (Fig S3d9). The tree of taxa shows that the relationship between Crenarchaeota and Euryarchaeota is nearer than the relationship between Euryarchaeota and Eukarya (Fig S3d9). The tree of taxa based on the evolution space agrees with the three domain tree rather than the eocyte tree [26] [27] [28] [29] [30]. This result is accordant with the observations that the features of genomic codon distributions between Crenarchaeota and Euryarchaeota are also more similar than that between Euryarchaeota and Eukarya in Subsection 2.2.

Life originated in the chiral chemistry in the roadmap scenario. The basic pattern of the genetic code determines the main branches of the tree of life, so biodiversity is an inborn feature of life. Thus the primordial ecosystem originated from the evolution of the genetic code, and the living system is a natural expansion of the primordial ecosystem. The living system of many species can adapt to the non-chiral surroundings through the birth and death of diverse individuals in the ecological cycle; however a system with only one species never has had chance to survive in the non-chiral surroundings. Hence the last universal common ancestor becomes unnecessary in the theories of evolution. And the phylogenetic relationships among species ought to form a circular network rather than a tree.

## 3 Reconstruction of the Phanerozoic biodiversity curve

### 3.1 Explanation of the trend of the Phanerozoic biodiversity curve based on genome size evolution

Evolution has taken place and is continuing to occur at both the species level and the sequence level. History archives of biodiversity on the earth have been stored not only in the fossil records at the species level [31] [32] [33] but also in the genomes of contemporary species at the sequence level. A Phanerozoic biodiversity curve can be plotted based on the fossil records [32] (Fig 4C, S4a8). The growth trend of this curve is exponential [32] (Fig 4C, S4a8), and the deviations from the exponential trend represent extinctions and originations [32] (Fig 4B, S4a8). Environmental changes substantially influence the evolution of life at the species level, but do not directly impact on the molecular evolution. Genome sizes are the most coarse-grained representations of species at the sequence level. So, the evolution of life can be outlined by the evolution of genome size. And the genome size increased mainly via large scale duplications. Additionally considering the relative invariance of the genomic codon distributions in the duplications (Fig S3a4), the features of species at the sequence level do not change in general through genome size evolution. So, it is relatively independent between the evolution at the species level and the evolution at the sequence level. In this paper, a split and reconstruction scheme is proposed to explain the Phanerozoic biodiversity curve (Fig 4C, S4a8, S4c1): the exponential growth trend of the curve is postulated to be driven by the genome size evolution (Fig 4A), while the extinctions and originations in the fluctuations of the curve is postulated due to the Phanerozoic eustatic and climatic changes (Fig 4B).

It is necessary to shed light on the genome size evolution. It will be shown that the growth trend of genome size evolution is also exponential in general [34] [35]. The genome size distributions of species in certain taxa generally satisfy logarithmic normal distributions, according to the statistical analysis of genome sizes in the databases Animal Genome Size Database [36] and Plant DNA C-values Database [37] (Fig S4a1). The genome size distributions for the 7 higher taxa of animals and plants, namely, diploblastica, protostomia, deuterostomia, bryophyte, pteridophyte, gymnosperm and angiosperm, generally satisfy logarithmic normal distributions, respectively. The genome size distribution for all the species in the two databases satisfies logarithmic normal distribution very well, due to the additivity of normal distributions (Fig S4a1).

The logarithmic normal distributions of genome sizes for taxa can be simulated by a stochastic model (Fig S4a2). Based on the duplication mechanism, the genome size evolution can be simulated by the stochastic model. The genome sizes obtained at a certain time in the simulation represent the genome sizes of species in a taxon at that time, which satisfy logarithmic normal distribution. The simulation indicates that, along the direction of time, the logarithmic mean of genome sizes for a taxon *G*_*logMean*_ increases, and the logarithmic standard deviation of genome sizes for a taxon *G*_*logS D*_ also increases (Fig S4a2). According to the simulation, the logarithmic mean of genome sizes for the ancestor of a taxon at the origin time *G*_*orilogMean*_ can be estimated by the genome sizes at present. Details are as follows. When genome size evolves in an exponential growth trend, *G*_*orilogMean*_ at the origin time is always less than *G*_*logMean*_ at present for a certain taxon. The greater *G*_*logS D*_ is (e.g. 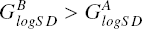 in Fig S4a2), the earlier the origin time is (e.g. 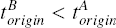 in Fig S4a2), and *origin orilogMean* thereby the less *G*_*orilogMean*_ is (e.g. 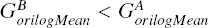 in Fig S4a2). Thus, the genome size for ancestor at the origin time *G*_*orilogMean*_ is obtained by calculating the genome sizes at present: *G*_*logMean*_ minus *G*_*logS D*_ times an undetermined factor χ (Fig S4a2, Fig S4a7).

Based on the genome sizes of contemporary species in the databases, the logarithmic means of genome sizes *G*_*logMean*_ and the logarithmic standard deviations of genome sizes *G*_*logS D*_ are obtained for the 7 higher taxa of animals and plants, respectively (Fig S4a7). The genome sizes for the ancestors of the 7 higher taxa at their origin times *G*_*orilogMean*_ are less than the corresponding *G*_*logMean*_ at present, whose differences can be estimated by the corresponding *G*_*logS D*_ times an undetermined factor χ, respectively (Fig S4a7). The trend of the genome size evolution cannot be worked out just only based on the genome sizes of contemporary species. Furthermore, a chronological reference has to be introduced here. The origin times for taxa can be estimated according to the fossil records. In this paper, the origin times for the 7 higher taxa are assumed as follows respectively: diploblastica, 560 *Ma*; protostomia, 542 *Ma* (Ediacaran-Cambrian); deuterostomia, 525 *Ma*; bryophyte, 488.3 *Ma* (Cambrian-Ordovician); pteridophyte, 416.0 *Ma* (Silurian-Devonian); gymnosperm, 359.2 *Ma* (Devonian-Carboniferous); angiosperm, 145.5 *Ma* (Jurassic-Cretaceous) [38] [39] [40]. The main results in this paper cannot be affected substantially by the disagreements among the origin times in the literatures or the expansion of the databases. A rough exponential growth trend of genome size evolution is observed in the plot whose abscissa and ordinate represent the origin time and *G*_*logMean*_ respectively (Fig S4a7), where the differences between the genome sizes at the origin time and that at present are not considered yet. A more reasonable plot is introduced by taking *G*_*orilogMean*_ rather than *G*_*logMean*_ as the ordinate (Fig S4a7). In the calculation, *G*_*orilogMean*_ are obtained respectively for the 7 higher taxa by (Fig S4a7)

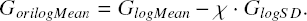

According to the regression analysis, a regression line approximates the relationship between the origin time and *G*_*orilogMean*_ (Fig S4a7). Let the intercept of the regression line be the logarithmic mean of genome sizes for all the species in the databases, then the undetermined factor is χ = 1.57 (Fig S4a7). The slope of the regression line represents the exponential growth speed of genome size evolution, which was doubled for each 177.8 *Ma* (Fig S4a7, 4A). It does not violate common sense that the genome size at the time of the origin of life about 3800 *Ma* is about several hundreds base pairs, according to the regression line.

There are more evidences to support the method for calculating the ancestors’ genome size based on the data of contemporary species. The values of *G*_*logMean*_, *G*_*logS D*_ and *G*_*orilogMean*_ are obtained for the 19 animal taxa or for the 53 angiosperm taxa, respectively (Fig S4a3). The numbers of species in these taxa are great enough for statistical analysis. The chronological order for the 19 animal taxa is obtained by comparing *G*_*orilogMean*_ (Fig S4a5). The order of the three superphyla is suggested to be: 1 diploblastica, 2 protostomia, 3 deuterostomia, which can be inferred from the chronological order of the 19 animal taxa and the superphylum affiliation of the 19 taxa (Fig S4a5). This result supports the three-stage pattern in Metazoan origination based on fossil records [41] [42] [43] [44] [45] [46] [47] [48]. The chronological order for the 53 angiosperm taxa is also obtained by comparing *G*_*orilogMean*_ (Fig S4a6), which confirms the common sense that dicotyledoneae originated earlier than monocotyledoneae. Let the abscissa and ordinate represent respectively *G*_*orilogMean*_ and *G*_*logMean*_ for the 7 higher taxa and 19 + 53 taxa, where the variation ranges of logarithm of genome sizes *G*_*logMean*_ ± *G*_*logS D*_ were also indicated on the ordinate. The upper triangular distribution of the genome size variations thus obtained (Fig S4a4) reveals intrinsic features of the genome size evolution. The greater *G*_*orilogMean*_ is, the greater *G*_*logMean*_ is, and the less *G*_*logS D*_ is (Fig S4a4). The upper vertex of the triangle indicates approximately the upper limit of genome sizes of contemporary species. The upper triangular distribution (Fig S4a4) can be explained by the simulation of genome size evolution (Fig S4a2). The simulation shows that *G*_*orilogMean*_ and *G*_*logMean*_ increase along the evolutionary direction; the earlier the time of origination is, the greater *G*_*logS D*_ is (Fig S4a2, Fig S4a4).

The exponential growth trend of Phanerozoic biodiversity curve can be explained by the exponential growth trend of genome size evolution. Bambach et al. obtained the Phanerozoic biodiversity curve based on fossil records, which satisfies the exponential growth trend (Fig 4C). The number of genera was doubled for each 172.7 *Ma* in the Phanerozoic eon (Fig 4A). The genome size was doubled for each 177.8 *Ma* in the Phanerozoic eon (Fig 4A). The exponential growth speed for the number of genera agrees with the exponential growth speed for the genome size. It is conjectured that the genome size evolution at the sequence level drives the evolution of life at the species level.

Note that Bambach et al.’s curve is chosen as the Phanerozoic biodiversity curve in this paper. It is based on the following considerations. Based on fossil records, Sepkoski obtained the Phanerozoic biodiversity curve at family level for the marine life first [49], then obtained the curve at genus level [50]. Sepkoski did not include single fossil records in order to reduce the impact on the curve at genus level from the samplings and time intervals. In order to reduce such an impact furthermore, Bambach et al. chose only boundary-crossing taxa in construction of the Phanerozoic biodiversity curve at genus level [32]. Rohde and Muller obtained a curve for all genera and another curve for well-resolved genera (removing genera whose time intervals are epoch or period, or single observed genera) respectively based on Sepkoskis Compendium, which can roughly be taken as the upper limit and lower limit of the Phanerozoic curve (Fig 4C). These Phanerozoic biodiversity curves are roughly in common in their variations, from which the five mass extinctions O-S, F-F, P-Tr, Tr-J, K-Pg can be discerned. There are several models to explain the growth trend of the Phanerozoic biodiversity curve. Sepkoski and other co-workers tried to explain the growth trend by 2-dimensional Logistic model [51] [52] [53]. Benton et al. explained it by exponential growth model [54]. The Bambach et al.’s curve is lower in the Paleozoic era, and obviously higher in the Cenozoic era than Sepkoski’s curve. It is more obvious that the Bambach et al.’s curve is in exponential growth trend than Sepkoski’s curve.

The exponential growth trend and the net fluctuations of the Bambach et al.’s biodiversity curve can be explained at both the molecular level and the species level respectively by a split and reconstruction scheme (Fig S4a8). In this scheme, the exponential growth trend of biodiversity is attributed to the driving force from the genome size evolution, in view of the approximate equality between the growth speed of number of genera and the growth speed of genome size (Fig 4A). The homochirality of life, the genetic code and the tree of life are generally invariant with the duplications of genome sizes. It maintains the features of life at the sequence level. Expansion of genome size provides conditions at the sequence level for the innovation of biodiversity at the species level. According to the split and reconstruction scheme, the extinctions and originations by the deviations from the exponential growth trend of the biodiversity curve is interpreted as a biodiversity net fluctuation curve. An unnormalised biodiversity net fluctuation is obtained by subtracting the growth trend from the logarithmic Phanerozoic biodiversity curve (Fig S4a8), and the biodiversity net fluctuation curve (Fig 4B, S4b5) is defined by dividing its standard deviation from the biodiversity net fluctuation curve. The biodiversity net fluctuation curve can be explained by the Phanerozoic eustatic curve and Phanerozoic climatic curve at the species level in the next subsection (Fig 4B).

### 3.2 Explanation of the variation of the Phanerozoic biodiversity curve based on climatic and eustatic data

According to the split and reconstruction scheme for the Phanerozoic biodiversity curve, the growth trend of biodiversity is attributed to the genome size evolution at the sequence level, and the net fluctuations deviated from the growth trend is attributed to the couplings among earth’s spheres. The history of biodiversity in the Phanerozoic eon can be outlined by the biodiversity net fluctuation curve (Fig 4B): the curve reached the high point after the Cambrian radiation and the Ordovician radiation; then the curve reached the low point after a triple plunge of the O-S, F-F and P-Tr mass extinctions; and afterwards the curve climbed higher and higher except for mass extinctions and recoveries around Tr-J and K-Pg. The biodiversity net fluctuation curve can be reconstructed through a climato-eustatic curve based on climatic and eustatic data (Fig 4B), which is free from the fossil records. Accordingly, it is advised that the tectonic movement can account for the five mass extinctions.

Numerous explanations on the cause of mass extinctions have been proposed, such as climate change, marine transgression and regression, celestial collision, large igneous province, water anoxia, ocean acidification, release of methane hydrates and so on. According to a thorough study of control factors on mass extinctions, it is almost impossible to explain all the five mass extinctions by a single control factor. Although the O-S mass extinction may be attributed to the glacier age, the other mass extinctions almost happened in high temperature periods. The other glacier ages in the Phanerozoic eon had always avoided the mass extinctions. Although the mass extinctions tend to happen with marine regressions, it is commonly believed that no marine regression happened in the end-Changhsingian, the greatest mass extinction [55] [56] [57] [58] [59]. Large igneous province may play roles only in the last three mass extinctions [60] [61]. The K-Pg mass extinction was most likely to be caused by celestial collision, but there are still significant doubts on this explanation [62] [63] [64]. According to the high resolution study of the mass extinctions based on fossil records, more understandings on the complexity of mass extinctions arise. For example, the P-Tr mass extinction can be divided into two phases: End-Guadalupian and end-Changhsingian [65] [66] [67] [68] [69]; furthermore the end-Changhsingian stage can be subdivided into main stage (B line) and final stage (C line) [70] [71], and the course of extinction may happened very rapidly [72] [73] [74] [75] [76]. Such a complex pattern of the mass extinction challenges any explanations on this problem. it is also almost impossible to attribute a mass extinction to a single control factor. Based on the above discussions, it is wise to consider a multifactorial explanation for the mass extinctions. In consideration of the interactions among the lithosphere, hydrosphere, atmosphere and biosphere, the tectonic movement of the aggregation and dispersal of Pangaea influenced the biodiversity net fluctuation curve at several time scales through the impacts of hydrosphere and atmosphere (Fig 4B).

Before explaining the mass extinctions based on the tectonic movement, two preparations are required: first, to construct a Phanerozoic eustatic curve; second, to construct a Phanerozoic climatic curve.

The Phanerozoic eustatic curve is based on the following two results (Fig S4b3): (i) Haq sea level curve [77] [78], (ii) Hallam sea level curve [79]. In 1970s, Seismic stratigraphy developed a method called sequence stratigraphy to plot the sea level fluctuations. Vail in the Exxon Production Research Company obtained a sea level curve based on this method, which concerns controversial due to business confidential data. Haq et al. replotted a sea level curve for the mesozoic era and cenozoic era also based on this method [77], and a sea level curve for the paleozoic era [78]. The Haq sea level curve here is obtained by combining these two curves (Fig S4b3). Hallam obtained a sea level curve by investigating large amounts of data on sea level variations [79] (Fig S4b3). Generally speaking, it is not quite different between the Haq sea level curve and the Hallam sea level curve. Therefore, a Phanerozoic eustatic curve can be obtained by averaging the Haq sea level curve and the Hallam sea level curve after nondimensionalization at equal weights (Fig S4b3).

The Phanerozoic climatic curve is based on the following four results (Fig S4b2): (*i*) the climatic gradient curve based on climate indicators in the Phanerozoic eon [80] [81] (Fig S4b1); (*ii*) the atmospheric *CO*_2_ fluctuations in the Berner’s models that emphasise the weathering role of tracheophytes [82] [83] [84]; (*iii*) the marine carbonate ^87^*S r*/^86^*S r* record in the Phanerozoic eon in Raymo’s model [85] [86]. The more positive ^87^*S r*/^86^*S r* values, assumed to indicate enhanced *CO*_2_ levels and globally colder climates; more negative ^87^*S r*/^86^*S r* values, indicating enhanced sea-floor speeding and hydrothermal activity, are associated with increased *CO*_2_ levels; (*iv*) the marine *δ*^18^*O* record in the Phanerozoic eon based on thermodynamic isotope effect [86] [87]. The above four independent results are quite different to one another (Fig S4b2). Considering the complexity of the problem on the Phanerozoic climate change yet, all the four methods are reasonable in principle on their grounds respectively. A consensus Phanerozoic climatic curve is obtained based on the above four independent results after nondimensionalization at equal weights. Namely, the Phanerozoic climatic curve is defined as the average of nondimensionalized climate indicator curve [80] [81], Berner’s *CO*_2_ curve [82], 87*S r*/^86^*S r* curve [85] [86] and *δ*^18^*O* curve [86] [87] (Fig S4b2).

Nondimensionalization is a practical and reasonable method to obtain the consensus Phanerozoic climatic curve. As the temperature in a certain time is concerned, when the four results agree to one other, the common temperatures are adopted; when the four results disagree, the majority consensus temperatures are adopted (Fig S4b2). According to the Phanerozoic climatic curve, the temperature was high from Cambrian to Middle Ordovician (Fig S4b2); the temperature was low during O-S (Fig S4b2); the temperature increases from Silurian to Devonian, and it reach high during F-F (Fig S4b2); the temperature is extreme low during D-C (Fig S4b2); the temperature rebounded slightly during Missisippian, and the temperature became lower again during Pennsylvanian, and the temperature decreased to extreme low during early Permian (Fig S4b2); the temperature increase in Permian, and especially the temperature increase rapidly during the Lopingian, and reach an extreme high peak during P-Tr (Fig S4b2); the temperature is high from Lower Triassic to Lower Jurassic, and it reached to extreme high during Lower Jurassic (Fig S4b2); the temperature became low from Middle Jurassic to Lower Cretaceous, where it is coldest in the Mesozoic era during J-K, which was not low enough to form glaciation (Fig S4b2); the temperature is high again from Upper Cretaceous to Paleocene, and the detailed high temperature during the Paleocene-Eocene event is observed (Fig S4b2); the temperature decreased from Eocene to present, especially reached to the minimum in the Phanerozoic eon during Quaternary (Fig S4b2). Four glacier ages are observed in the Phanerozoic eon (Fig S4b2): Hirnantian glacier age, Tournaisian glacier age, early Permian glacier age and Quaternary glacier age, especially bipolar ice sheet at present. The Phanerozoic climatic curve also indicates the extreme low temperature during J-K (Fig S4b2), which corresponds to possible continental glaciers at high latitudes in the southern hemisphere [88]. In short, the fine-resolution Phanerozoic climatic curve agrees with the common sense on climate change at the tectonic time scale.

The Phanerozoic climatic curve is taken as a standard curve to evaluate the four independent methods. The climate indicator curve is the best result among the four. It accurately reflects the four high temperature periods and four low temperature periods, and the varying amplitudes of the curve are relatively modest and reasonable. But it overestimated the temperature during Pennsylvanian (Fig S4b2). The *δ*^18^*O* curve is also reasonable. It also reflects the four high temperature periods and four low temperature periods. But it overestimated the temperature during Guadalupian, and underestimated the temperature during Jurassic (Fig S4b2). The 87*S r*/^86^*S r* curve is reasonable to some extent. It reflects the high temperature during Mesozoic and low temperature during Carboniferous and Quaternary. But it underestimated the temperature during Cambrian, and overestimated the temperature during O-S and J-K (Fig S4b2). The Berner’s *CO*_2_ curve is relatively rough. It reflects the high temperature during Mesozoic, But it overestimated the temperature from Cambrian to Devonian (Fig S4b2).

As the climate changes at the tectonic time scale is concerned, the Berner’s *CO*_2_ curve indicates relatively long term climate changes; the climate indicator curve indicates relatively mid term climate changes; and the ^87^*S r*/^86^*S r* and *δ*^18^*O* curves indicate relatively short term climate changes (Fig S4b2). The scale of the variation of the Phanerozoic climatic curve is relatively accurate because this consensus climatic curve is obtained by combining all the different time scales of the above four climatic curves. Four high temperature periods Cambrian, Devonian, Triassic, K-Pg and four low temperature periods O-S, Carboniferous, J-K, Quaternary appeared alternatively in the Phanerozoic climatic curve, which generally reflect four periods in the climate changes in the Phanerozoic eon (Fig S4b2). Such a time scale of the four periods of climate changes is about equal to the average interval of mass extinctions, and is also equal to the time scale of the four periods of the variation of the relative abundance of microbial carbonates in reef during Phanerozoic [89] [90] [91] [92]. The agreement of the time scales indicates the relationship between the climate change and the fluctuations of biodiversity in the Phanerozoic eon.

The sea level fluctuations and the climate change have an important impact on the fluctuations of biodiversity. With respect to the impact of the sea level fluctuations, this paper follows the traditional view that sea level fluctuations play positive contributions to the fluctuations of biodiversity. With respect to the impact of the climate change, this paper believes that climate change plays negative contributions to the fluctuations of biodiversity. The climate gradient curve is opposite to the climate curve (Fig S4b2). The higher the temperature is, the lower the climate gradient is, and consequently the less the global climate zones are; while the lower the temperature is, the higher the climate gradient is, and consequently the more the global climate zones are. When the climate gradient is high, there were tropical, arid, temperate and frigid zones among the global climate zones, which boosts the biodiversity. So, this paper believes that the biodiversity increases with the climate gradient. This point of view agrees with the following observations. The biodiversity rebounded during the low temperature period Carboniferous (Fig S4b5), and increased steadily during the ice age during Quaternary (Fig S4b5); and the four mass extinctions F-F, P-Tr, Tr-J, K-Pg appeared in high temperature periods (Fig S4b5). Although the O-S mass extinction occurred in the ice age (Fig S4b5), this case is quite different from other mass extinctions. In the first mass extinction during O-S, mainly the Cambrian fauna, which originated in the high temperature period, cannot survive from the O-S ice age. Mass extinctions no longer occurred thereafter in the ice ages for those descendants who have successfully survived from the O-S ice age (Fig S4b5).

Based on the above analysis, the eustatic curve and the climatic gradient curve is positively related to the fluctuations of biodiversity in the Phanerozoic eon. Accordingly, a climato-euctatic curve is defined by the average of the nondimensionaliezed eustatic curve and the nondimensionaliezed climatic grandaunt curve, which indicates the impact of environment on biodiversity (Fig S4b4). It should be emphasised that the climato-euctatic curve is by no means based on fossil records, which is only based on climatic and eustatic data. It is obvious that the climato-euctatic curve agrees with the biodiversity net fluctuation curve (Fig 4B). Judging from the overall features, the outline of the history of the Phanerozoic biodiversity can be reconstructed in the climato-eustatic curve, as far as the Cambrian radiation and the Ordovician radiation, the Paleozoic diversity plateau, the P-Tr mass extinction, and the Mesozoic and Cenozoic radiation are concerned (Fig 4B). Judging from the mass extinctions, the five mass extinctions O-S, F-F, P-Tr, Tr-J, K-Pg can be discerned from the five steep descents in the climato-eustatic curve respectively (Fig 4B, S4b7). As to K-Pg especially, there is obviously a pair of descent and ascent around K-Pg in the climato-eustatic curve (Fig 4B), so celestial explanations are actually not necessary here. Judging from the minor extinctions, most of the minor extinctions can be discerned in the climato-eustatic curve in details, as far as the following are concerned: 3 extinctions during Cambrian, Cm-O extinction, S-D extinction, 2 extinctions during Carboniferous, 1 extinction during Permian, 1 extinction during Triassic, 3 extinctions during Jurassic, 2 extinctions during Cretaceous, 2 extinctions during Paleogene, 1 extinction during Neogene and the extinction at present (Fig 4B, S4b7). Judging from the increment of the biodiversity, the originations and radiations can be discerned from the ascents in the climato-eustatic curve, as far as the following are concerned: the Cambrian radiation, the Ordovician radiation, the Silurian radiation, the Lower Devonian diversification, the Carboniferous rebound, the Triassic rebound, the Lower Jurassic rebound, the Middle Cretaceous diversification, the Paleogene rebound (Fig 4B, S4b7). The derivative of the climato-eustatic curve indicates the variation rates (Fig 4B, S4b7). The extinctions are discerned by negative derivative, and the rebounds and radiations by positive derivative. It is obvious to discern the five mass extinctions, most of minor extinctions, and rebounds and radiations in the derivative curve of the climato-eustatic curve, just like in the climato-eustatic curve (Fig 4B, S4b7).

The interactions among the lithosphere, hydrosphere, atmosphere, biosphere are illustrated by the eustatic curve, climatic curve, biodiversity net fluctuation curve and the aggregation and dispersal of Pangaea (Fig S4b5, Fig S4b6). There were about two long periods in the eustatic curve, which were divided by the event of the most aggregation of Pangaea (Fig S4b5). The first period comprises the grand marine transgression from the Cambrian to the Ordovician and the grand marine regression from the Silurian to the Permian, and the second period comprises the grand marine transgression during Mesozoic and the grand regression during Cenozoic (Fig S4b5).These observations reveal the tectonic cause of the grand marine transgressions and regressions in the Phanerozoic eon, considering the aggregation and dispersal of Pangaea. The coupling between the hydrosphere and the atmosphere can be comprehended by comparing the eustatic curve and the climatic curve. The global temperature was high during both the grand marine transgressions (Fig S4b5). And all the four ice ages in the Phanerozoic occurred during the grand marine regressions (Fig S4b5). Generally speaking, the climate gradient curve varies roughly synchronously with the eustatic curve from the Cambrian to the Lower Cretaceous along with the aggregation of Pangaea, except for the regression around the Tournaisian ice ages (Fig S4b5). And the climate gradient curve varies roughly oppositely with the eustatic curve thereafter from the Upper Cretaceous to the present along with the dispersal of Pangaea (Fig S4b5); namely the climate gradient curve increases as the eustatic curve decreases. These observations reveal the tectonic cause of the climate change at tectonic time scale in the Phanerozoic eon, considering the impact on the climate from the grand transgressions and regressions. The biodiversity responses almost instantly to both the eustatic curve and the climatic curve, because the life spans of individuals of any species are much less than the tectonic time scale. Hence, the biodiversity net fluctuation curve agrees with the climato-eustatic curve very well (Fig 4B). In conclusion, the tectonic movement has substantially influenced the biodiversity through the interactions among the earth’s spheres (Fig S4b5, Fig S4b6).

The interactions among the earth’s spheres can explain the outline of the biodiversity fluctuations in the Phanerozoic eon. During Cambrian and Ordovician, both the sea level and the climate gradient increased, so the biodiversity increased (Fig S4b5). From Silurian to Permian, both the sea level and the climate gradient decreased, so the biodiversity decreased (Fig S4b5), except for the biodiversity plateau during Carboniferous due to the opposite increasing climate gradient and decreasing sea level (Fig S4b5). From Triassic to Lower Cretaceous, both the sea level and the climate gradient increased, so the biodiversity increased (Fig S4b5). From Upper Cretaceous to present, the sea level decreased and the climate gradient increased rapidly, so the biodiversity increased (Fig S4b5).

The interactions among the earth’s spheres can explain the five mass extinctions within the same theoretical framework, though their causes are different in detail. The mass extinctions O-S, K-Pg occurred around the peaks in the eustatic curve respectively, and the P-Tr mass extinction around the lowest point in the eustatic curve. There were about two periods in the eustatic curve and about four periods in the climatic curve in the Phanerozoic eon. So the period of climate change was about a half of the period of sea level fluctuations, as far as their changes at the tectonic time scale are concerned (Fig S4b5). By comparison, the F-F mass extinction occurred in the low level of climate gradient. There was a triple plunge from the high level of biodiversity during the end-Ordovician to the extreme low level of biodiversity during Triassic in the biodiversity net fluctuation curve, which comprised a series of mass extinctions O-S, F-F, P-Tr (Fig 4B). It is interesting that a relatively less great extinction may occur after each of the triple plunge mass extinctions respectively: namely the S-D extinction after the O-S mass extinction, the Carboniferous extinction after the F-F mass extinction, and the Tr-J mass extinction after the P-Tr mass extinction (Fig 4B). Especially during the end of Permian, the aggregation of the Pangaea pull down both the eustatic curve and the climate gradient curve, and hence pull down the biodiversity curve (Fig S4b5). After a long continual descent in the biodiversity net fluctuation curve and after the two mass extinctions O-S, F-F, the ecosystem was more fragile at the end of Permian than in the previous period when biodiversity was at the high level, therefore the P-Tr mass extinction occurred as the most severe extinction event in the Phanerozoic eon (Fig 4B). The P-Tr mass extinction can be divided into the first phase at the end of Guadalupian and the second phase at the end of Changhsingian (Fig 4B). The extinction of Fusulinina mainly occurred at the end of Guadalupian; while the extinction of Endothyrina mainly occurred at the end of Changhsingian. Note that the number of Endothyrina did not decrease obviously at the first phase. Different extinction patterns in the above two phases indicate that the triggers for the two phases are different to each other. Marine regression occurred at the end of the Guadalupian, whereas it is indicated that no marine regression occurred at the end of the Changhsingian. The second phase of the end-Changhsingian extinction may be caused by the declining climate gradient (or increasing temperature). All the four independent methods in the above (climatic-sensitive sediments, Berner’s *CO*_2_, ^87^*S r*/^86^*S r* and *δ*^18^*O*) predict the rapidly increasing temperature at the end of the Changhsingian (Fig S4b2). Either Siberia trap or the methane hydrates can raise the temperature at the end of the Changhsingian, which may trigger the P-Tr mass extinction. Four mass extinctions F-F, P-Tr, Tr-J, K-Pg and the Cm-O extinction occurred during the low level of climate gradient (Fig S4b5); as a special case, the O-S mass extinction occurred during the high level of climate gradient (Fig S4b5). Taken together, these observations show that low climate gradient can contribute to the mass extinctions substantially. Generally speaking, the mass extinctions might occur coincidentally when brought together two or more factors among the low climate gradient, marine regression, and the like (Fig S4b5).

The interactions among the earth’s spheres can explain the minor extinctions, radiations and rebounds. The minor descents in the biodiversity net fluctuation curve relate to mid-term or short-term changes in sea level and climate at the tectonic time scale, which correspond approximately to the minor extinctions respectively [93] [94] (Fig 4B, S4b7). It is shown that most minor extinctions in the Paleozoic era corresponded to low climate gradient, and most minor extinctions during the Mesozoic and Cenozoic corresponded to marine regressions (Fig 4B, S4b7). After mass extinctions, both the climate gradient and sea level tended to increase; henceforth the radiations and rebounds occurred. Especially for the case of the P-Tr mass extinction, the climate gradient remained low throughout the Lower Triassic (Fig 4B, S4b5), so it took a long time for the recovery from the P-Tr mass extinction.

### 3.3 Reconstruction of the Phanerozoic biodiversity curve and the strategy of the evolution of life

Now, let’s reconstruct the Phanerozoic biodiversity curve. In Subsection 3.1, the exponential growth trend of the Phanerozoic biodiversity curve has been explained by the exponential growth trend of the genome size evolution (Fig 4A); in Subsection 3.2, the deviation from the exponential growth trend of the Phanerozoic biodiversity curve has been explained by the climato-eustatic curve (Fig 4B). By combining the exponential growth trend of the genome size evolution and the climato-eustatic curve, the biodiversity curve based on fossil records can be reconstructed based on the genomic, climatic and eustatic data (Fig 4C). The technique details are as follows (Fig S4c1). The reconstructed biodiversity net fluctuation curve is obtained based on the climato-eustatic curve times a certain weight, where the weight is the standard deviation of the biodiversity net fluctuation curve (Fig S4c1). And the slope and the intercept of the line that corresponds to the exponential growth trend of the reconstructed biodiversity is obtained as follows. Its slope is defined by the slope of the line that corresponds to the exponential growth trend of genome size evolution (Fig S4c1), and its intercept is defined by the intercept of the line that corresponds to the exponential growth trend of the biodiversity curve (Fig S4c1). The reconstructed biodiversity curve thus obtained agrees nicely with the Bambach et al.’s biodiversity curve within the error range of fossil records (Fig 4C).

The above split and reconstruction scheme of the Phanerozoic biodiversity curve can explain not only the exponential growth trend and the detailed fluctuations, but also the roughly synchronously declining origination rate and extinction rate throughout the Phanerozoic eon [95] [96] [97] [98] [99] (Fig 4D, S4d1, S4d2, S4d3). Ramp and Sepkoski first found the declining extinction rate throughout the Phanerozoic eon [101]. And Sepkoski also found the declining origination rate throughout the Phanerozoic eon [94]. The numbers of origination genera and extinction genera are marked in the Bambach et al.’s biodiversity curve. The origination rate and extinction rate are obtained from the numbers of origination or extinction genera divided by the corresponding time intervals (Fig S4d3). The difference between the origination rate and extinction rate is the variation rate of number of genera (Fig S4d3). It is shown that both the origination rate and extinction rate decline roughly synchronously throughout the Phanerozoic eon [102] (Fig S4d3).

The rough synchronicity between the origination rate and the extinction rate can be explained according to the split and reconstruction scheme. The Phanerozoic biodiversity curve can be divided into two independent parts: an exponential growth trend and a biodiversity net fluctuation curve (Fig S4a8, Fig S4c1). Notice that there is no declining trend in the climato-eustatic curve, the expectation value of which is constant (Fig 4B, S4d1); whereas the biodiversity curve was continuously boosted upwards by the exponential growth trend of genome size evolution (Fig 4A). The expectation value of the difference between the origination rate and extinction rate corresponds to the exponential growth trend of biodiversity boosted by the genome size evolution (Fig S4d1, Fig S4d2). If too large or too small origination rate brings the biodiversity to deviate from the exponential growth trend, the extinction rate has to response by increasing or decreasing so as to maintain the constant exponential growth trend of the biodiversity (Fig S4d2, Fig S4d3). The rough synchronicity between the origination rate and the extinction rate shows that the growth trend of biodiversity at the species level was constrained strictly by the exponential growth trend of genome size evolution at the sequence level.

The declining trends of the origination rate and the extinction rate can be explained according to the split and reconstruction scheme. It is necessary to assume that there is an upper limit of ability to create new species in the living system, which is due to the constraint of the earth’s environment at the species level and the limited mutation rate at the sequence level (Fig S4d1). Although there is no declining trend for the variation rate of the biodiversity net fluctuation curve (Fig S4b7, Fig S4d1), the trend of the variation rate of the Phanerozoic biodiversity curve tends to decline in general, because the exponential growth trend in genome size evolution is added in the denominator in the calculations (Fig 4D, S4d2, S4d3). The declining variation rate throughout the Phanerozoic eon shows that the genome size evolution played a primary role in the growth of biodiversity. The exponential growth trend of biodiversity driven by the genome size evolution can gradually weaken the impact on the biodiversity fluctuations from the changes of sea level and climate (Fig 4D, S4d2). The rough synchronicity between the origination rate and the extinction rate can explain both the declining origination rate and the declining extinction rate throughout the Phanerozoic eon (Fig S4d2, Fig S4d3).

The biodiversity evolves independently at the sequence level and at the species level. This ensures the adaptation of life to the earth’s changing environment. The essential feature of the living system is its homochirality. At the beginning of life, the homochirality, the genetic code, and the genomic codon distributions had established. The homochirality ensures the safety for the living system: the earth’s changing environment influenced the living system only at the species level, which cannot directly threaten the existence of life at the sequence level. The strategy of adaptation is that: life generates numerous redundant species at the sequence level; thus life sacrificed opportunity of survival for the vanished majority of the species in exchange for the present outstanding adaptability to the changing environment. By virtue of this strategy of adaptation, the chiral living system may adapt to a wide range of environmental changes on planets.

## Acknowledgements

My warm thanks to Jinyi Li for valuable discussions. Supported by the Fundamental Research Funds for the Central Universities.

## Index

**Section 1**

roadmap for the evolution of the genetic code (*the roadmap for short*)
initiation stage *of the roadmap*
expansion stage *of the roadmap*
ending stage *of the roadmap*
*origin of* the homochirality of life
*origin of* the genetic code
*origin of* the three domains of life
base pair of double helix DNA
base triplex of triplex DNA
relative stability of base triplex
base substitutions *in the roadmap*
codon pair *in the roadmap*
pair connection *in the roadmap*
route duality *in the roadmap Route* 0 − 3 *of the roadmap*
*Hierarchy* 1 − 4 *of the roadmap*
initial subset *in the roadmap*
recruitment order of codon pairs (#1 to #32)
recruitment order of amino acids (*No*. 1 to *No*. 20)
cooperative recruitment of codons and amino acids
non-standard codons *in the roadmap*
*origin of* stop codons
*origin of* tRNAs
branch node *in the roadmap*
leaf node *in the roadmap*
palindromic codon pair
non-palindromic codon pair
complementary single RNA strands
assignment scheme of anticodons
permutations of bases
combinations of bases
*explanation of* number of canonical amino acids
*explanation of* the wobble paring rule
*explanation of* codon degeneracy
minimum subset *in the roadmap*
facet of cube for route
biosynthetic families of amino acids
standard genetic code table
*GCAU* genetic code table
classification of aminoacyl-tRNA synthetases
chiral roadmap
winner-take-all competition between chiral roadmaps
*simulation of* the origin of homochirality
*explanation of* position specific GC content
*explanation of* stop codon usage
*explanation of* variation of amino acid frequencies
order *R*_10/10_ for amino acids
Phase *I* and *II* of amino acids

**Section 2**

genomic codon distribution
codon intervals in genome
correlation coefficient matrix
distance matrix *by genomic codon distributions*
tree of codons *by genomic codon distributions*
tree of species *by genomic codon distributions*
tree of eukaryotes *by genomic codon distributions*
rough invariance of genomic codon distribution
rough periodicity of genomic codon distribution
declining three-base-periodic fluctuations
amplitudes of genomic codon distribution
order of the average distribution heights
origin order of domains by slope of regression line
evolutionary status of Virus
*simulation of* genomic codon distributions
*explanation of* validity of composition vector tree
evolution space
fluctuation (*coordination of the evolution space*)
route bias (*coordination of the evolution space*)
hierarchy bias (*coordination of the evolution space*)
leaf node bias
classification of genomic codon distributions
non-dimensionalized evolution space
distance matrix *by the evolution space*
tree of species *by the evolution space*
tree of taxa *by the evolution space*
tree of eukaryotes *by the evolution space*
three-domain tree of life
eocyte tree of life

**Section 3**

split and reconstruction scheme
evolution of life at sequence level
evolution of life at species level
genome size evolution
Phanerozoic biodiversity curve
Phanerozoic climatic curve
Phanerozoic eustatic curve
exponential growth trend of the biodiversity curve
exponential growth trend in genome size evolution biodiversity net fluctuation curve
climate gradient curve
climto-eustatic curve
logarithmic normal distribution of genome sizes
logarithmic mean of genome sizes
logarithmic standard deviation of genome sizes
original logarithmic mean of genome sizes factor χ
*simulation of* genome size evolution
interactions among earth’s spheres five mass extinctions
control factors of mass extinctions
multifactorial explanation of mass extinctions
Haq sea level curve
Hallam sea level curve
climate indicator curve
Berner’s *CO*_2_ curve
marine ^87^*S r*/^86^*S r* curve
marine *δ*^18^*O* curve
average after nondimensionalization derivative curves
aggregation and dispersal of Pangaea
eustatic change at tectonic time scale
climatic change at tectonic time scale
instant response of biodiversity at tectonic time scale
tectonic cause of mass extinctions
outline of the biodiversity fluctuations
two phases of P-Tr mass extinction minor extinctions
radiations and rebounds of biodiversity
reconstructed Phanerozoic biodiversity curve
*explanation of* the declining trend of origination rate
*explanation of* the declining trend of extinction rate
synchronicity between origination and extinction
constraint on origination ability
redundant species created at the sequence level
strategy of adaptation

## Supplementary Figures

**Fig 1A.**
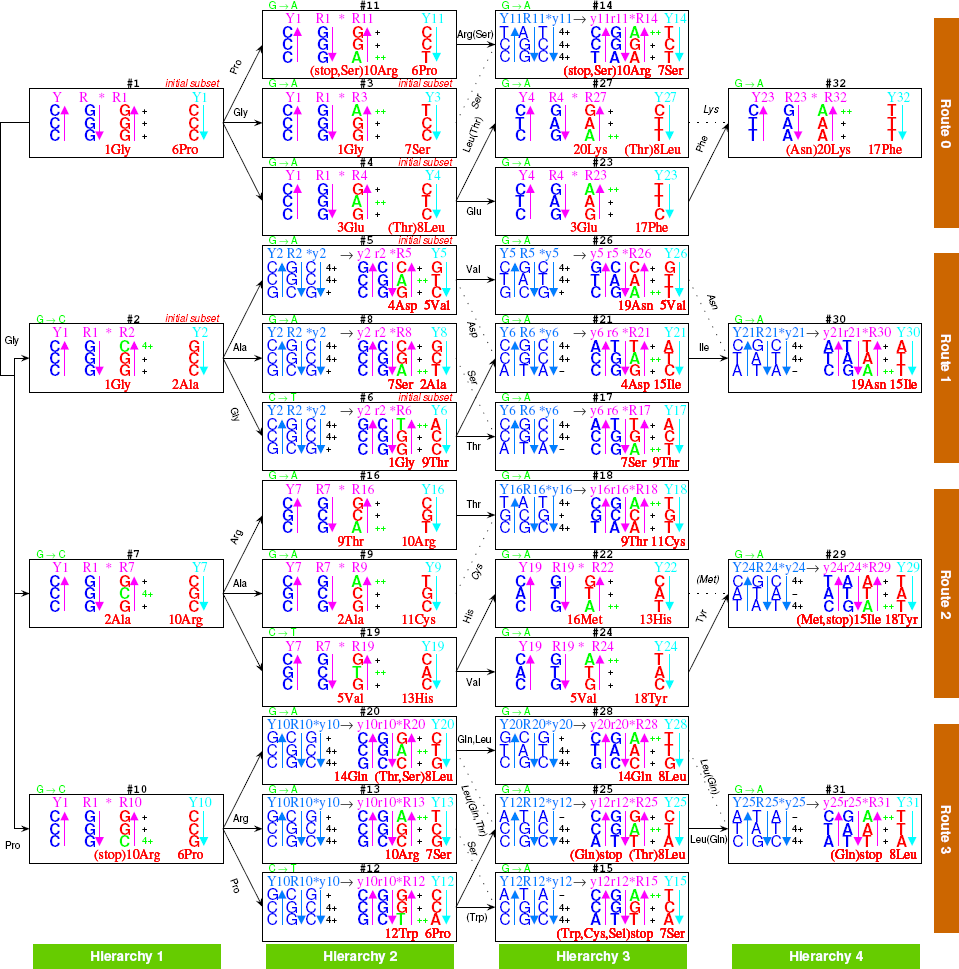
The roadmap for the evolution of the genetic code. The 64 codons formed from base substitutions in triplex DNAs are in red. Only three-base-length segments of the triplex DNAs are shown explicitly; the whole length right-handed triplex DNAs are indicated in Fig 1B. In each position #*n*, the #*n* codon pair on *Rn* and *Yn* is in red. The relative stabilities of the base triplexes (-, +, ++, 4+) are written to the right of the base triplexes, where the increasing relative stabilities of base triplexes in base substitutions are indicated in green. Each triplex DNA is denoted by three arrows, whose directions are from 5’ to 3’. The *YR R* triplex DNAs are in pink, and the *YR Y* triplex DNAs in azure. The recruitment order of codon pairs are from #1 to #32, and the recruitment order of the 20 amino acids are to the left of them respectively. Non-standard genetic codes are indicated by brackets beside the corresponding amino acids. The *Route* 0−3 and *Hierarchy* 1−4 are indicated to the right of and below the roadmap respectively. The evolution of the genetic code are denoted by black arrows, beside which pair connections are indicated by certain amino acids. Refer to an example in Fig 1B to understand details of the roadmap; refer to Fig S1a1 to understand the critical role of relative stabilities of base triplexes in achieving the real genetic code; refer to Fig 2A to see the origin of tRNAs; refer to Fig 2B to see the coherent relationship between the recruitment orders of codons and amino acids; refer to Fig 2C to see the codon degeneracy in the symmetric roadmap; and refer to Fig 2D to see the origin of homochirality of life.

**Fig S1a1.**
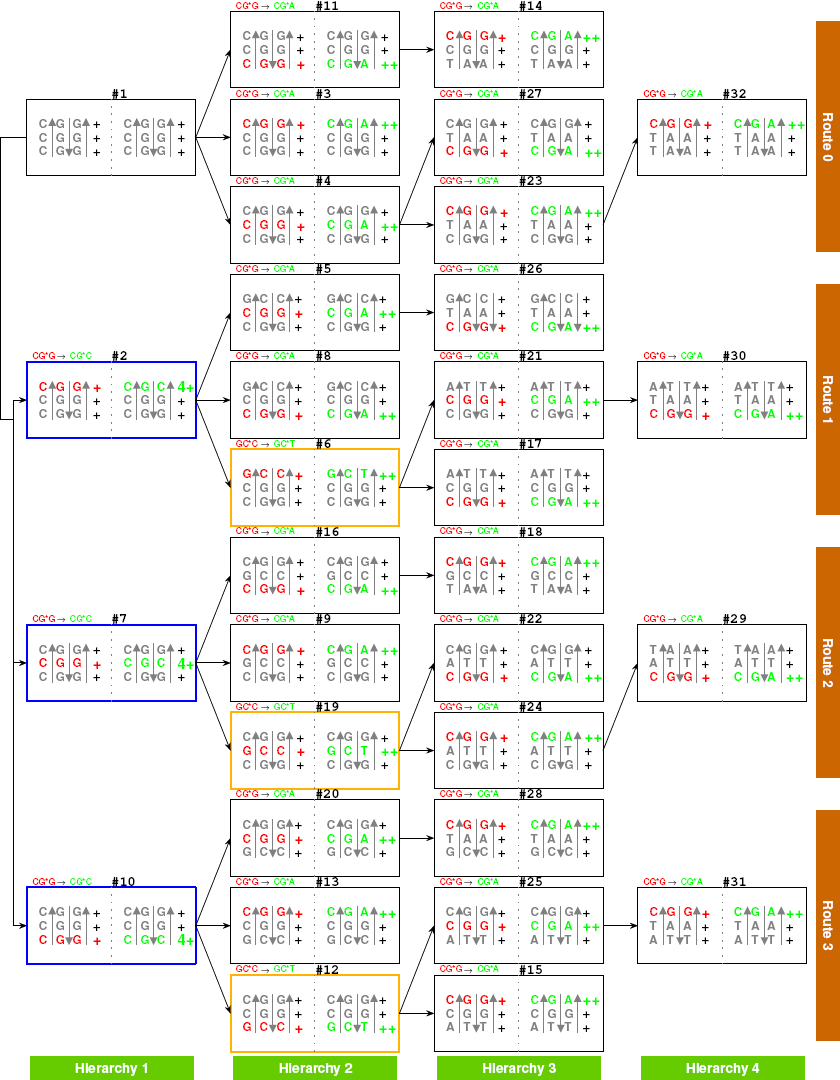
Explanation of the increasing relative stabilities of base triplexes in base substitutions along the roadmap Fig 1A. The base substitutions in the roadmap occur when the relative stabilities of base triplexes increase. The roadmap is the best result to avoid the unstable base triplexes. So, the universal genetic code is a narrow choice by the relative stabilities of base triplexes. The relative stability increases from (+) of the base triplex *CG * G* to (4+) of the base triplex *CG * C* at #2, #7 and #10 that initiates *Route* 1−3 respectively. *GC * C* (+) changes to *GC * T* (++) at #6, #19 and #12, and *CG * G* (+) changes to *CG * A* (++) at other positions in the roadmap.

**Fig 1B.**
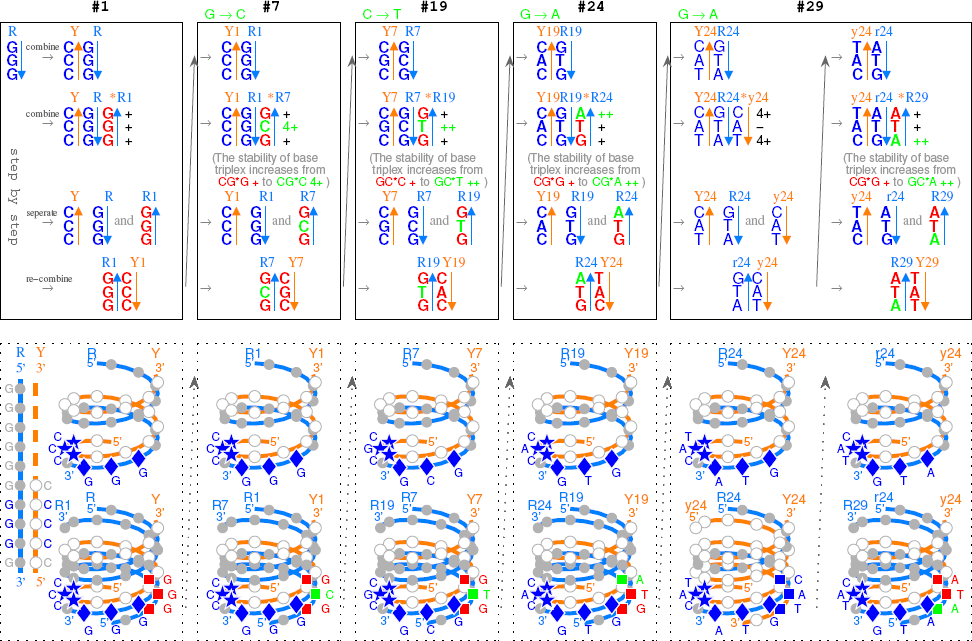
A detailed description of the roadmap in the right-handed triplex DNA picture, taking for example from #1 to #29. The evolution of the genetic code from #1, to #7, to #19, to #24, and at last to #29 are explained in detail in the upper boxes, and the corresponding right-handed single-stranded, double-stranded and triple-stranded DNAs are shown in the lower boxes, respectively.

**Fig 2A.**
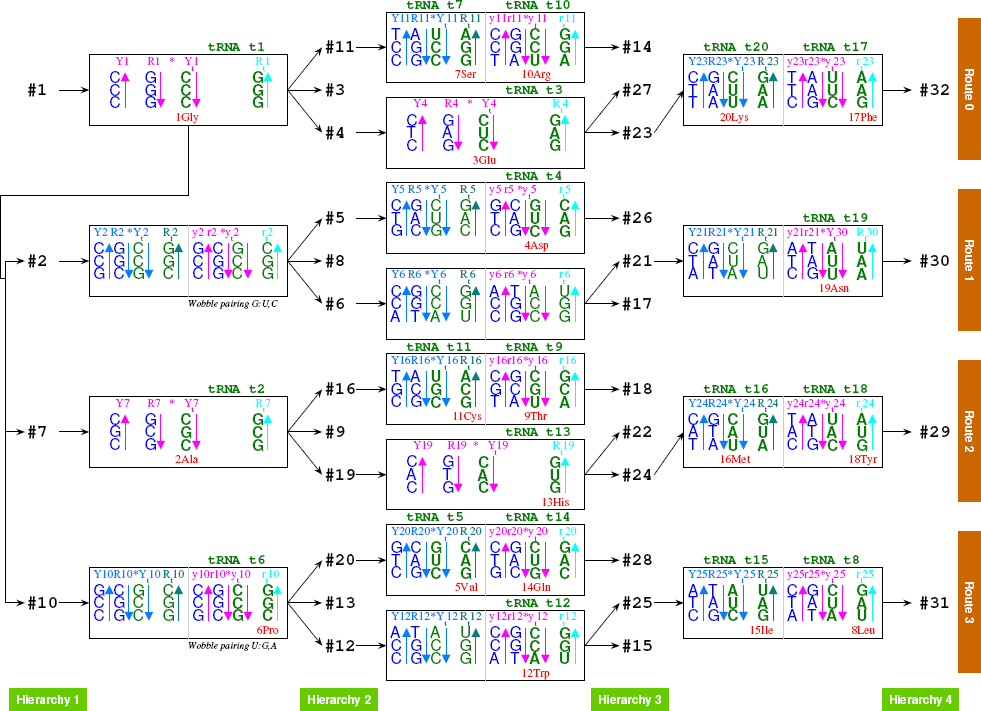
The tRNAs originated along the roadmap are able to transport all the canonical amino acids. Each triplex nucleic acid *YR * Y*_*t*_ or *YR * R*_*t*_ consists of two DNA strands *Y* and *R* and one RNA strand *Y*_*t*_ or *R*_*t*_. There are two possible ways for each codon pair to generate two complementary RNA strands (*Y*_*t*_ and *R*_*t*_), which consequently assemble into a tRNA (Fig S2a3).

**Fig S2a1.**
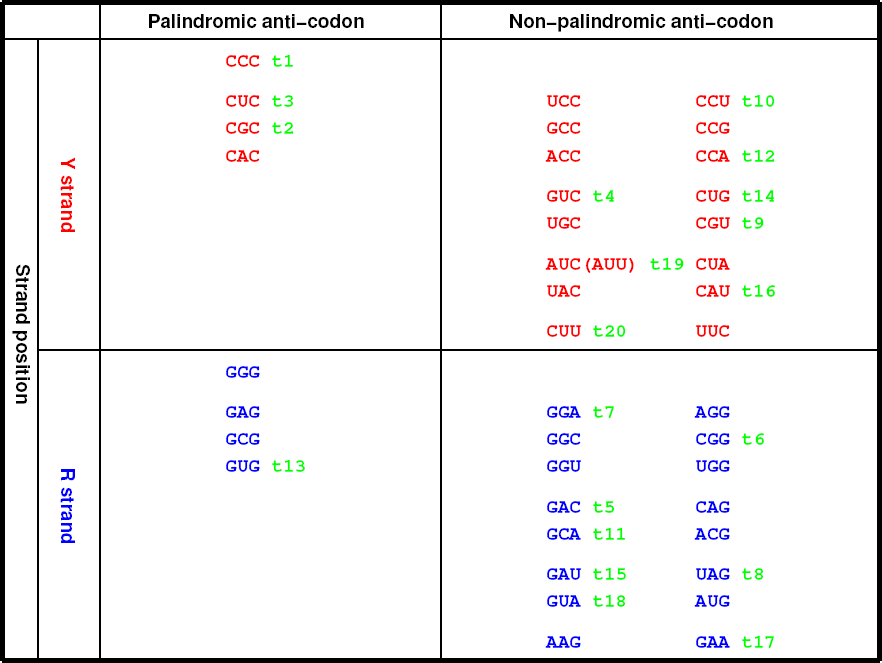
Single-strand RNA pairs at the branch nodes in the roadmap Fig 1A can form tRNAs: the tRNAs *t*1, *t*10, *t*3, *t*17, *t*2, *t*9, *t*13, *t*18 correspond to the codon pairs in *Route* 0 and *Route* 2, and the tRNAs *t*7, *t*20, *t*11, *t*16 correspond to the reverse codon pairs (considering the reverse operation) in *Route* 0 and *Route* 2 (Fig 2A). The codon pairs are reverse to each other between *Route* 1 and *Route* 3. The tRNAs *t*4, *t*19, *t*6, *t*5, *t*14, *t*12, *t*15, *t*8 correspond to codon pairs or reverse codon pairs in *Route* 1 and *Route* 3 (Fig 2A). In total, there are 20 tRNAs such obtained that correspond to the codon pairs or their reverse codon pairs at the branch nodes. The anti-codons concerned are arranged in the present figure by comparing between Y strand anti-codons and R strand anti-codons, as well as by comparing between palindromic anti-codons and non-palindromic anti-codons, and the rows in the present figure indicate the types of genomic codon distributions in Fig S3d1.

**Fig S2a2.**
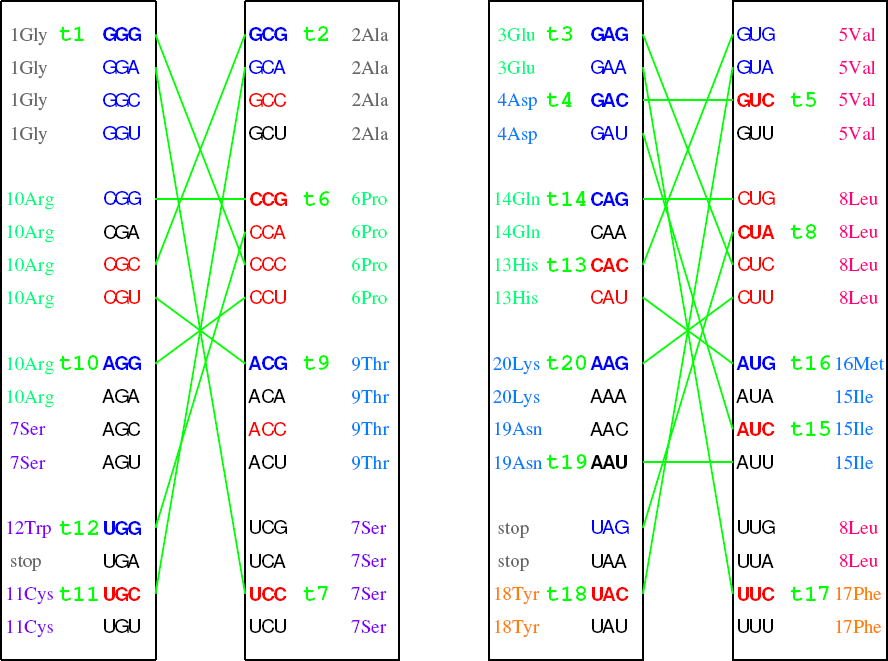
The 20 tRNAs *t*1-*t*20 (Fig 2A, S2a1) are able to carry all the 20 canonical amino acids according to the proper codon assignment in the present figure, where the blue and red codons denote the R- and Y-strand codons in Fig S2a1 respectively, and the green lines indicate the codon pairs.

**Fig S2a3.**
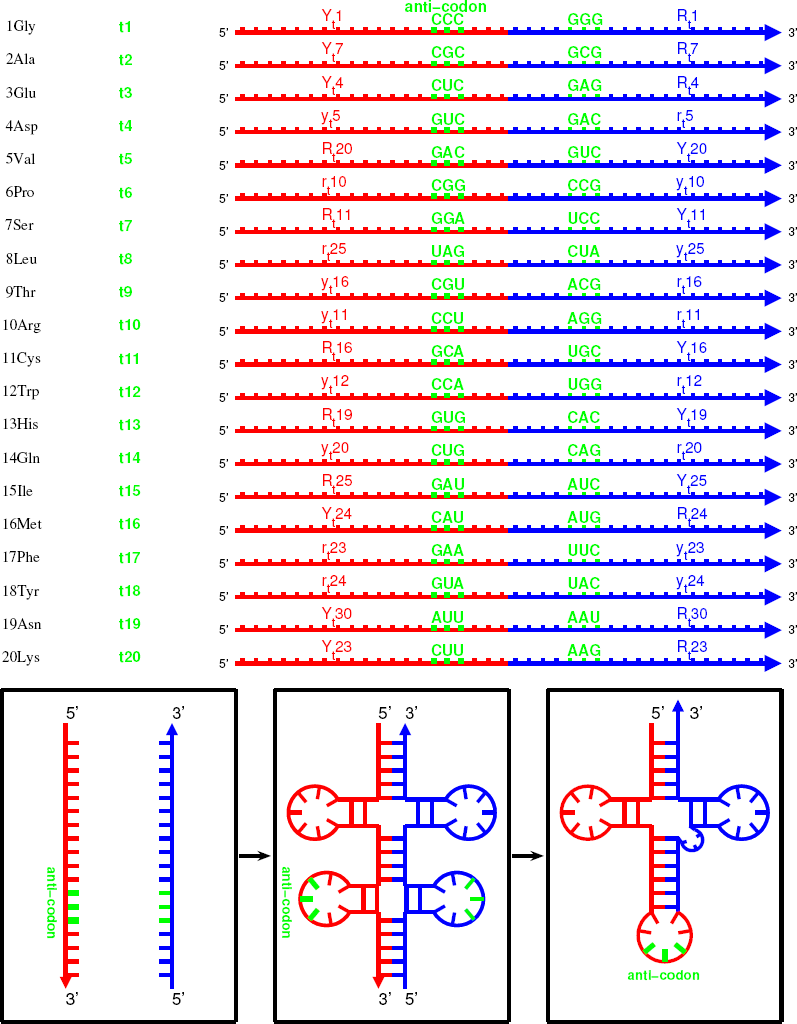
The tRNAs *t*1-*t*20 with anti-codons (Fig 2A, S2a1, S2a2) are listed here to carry the amino acids from *No*.1 to *No*.20 respectively. The two complimentary single-stranded RNAs for each tRNA join together, and fold into a cloverleaf shape by taking advantage of the complementarity between the two strands. The joining position of the two strands is near to the 3’ side of the anti-codon loop, which agrees with the position of introns in tRNA genes in observations. The additional codon in the complementary strand might be located around the variable loop, despite possibly great changes in this loop. This may relate to the paracodon in the tRNAs.

**Fig 2B.**
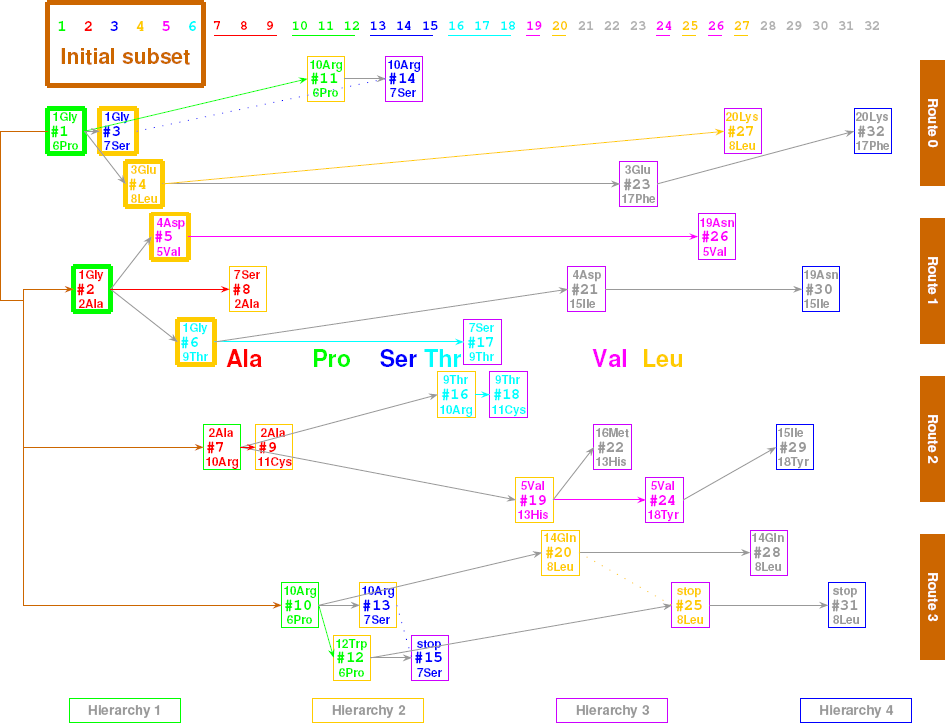
Cooperative recruitment of codons and amino acids along the roadmap. The codon pairs are plotted from left to right according to their recruitment order. The initial subset plays a crucial role in the expansion of the genetic code. The 6 biosynthetic families of the amino acid are distinguished by different colours.

**Fig S2b1.**
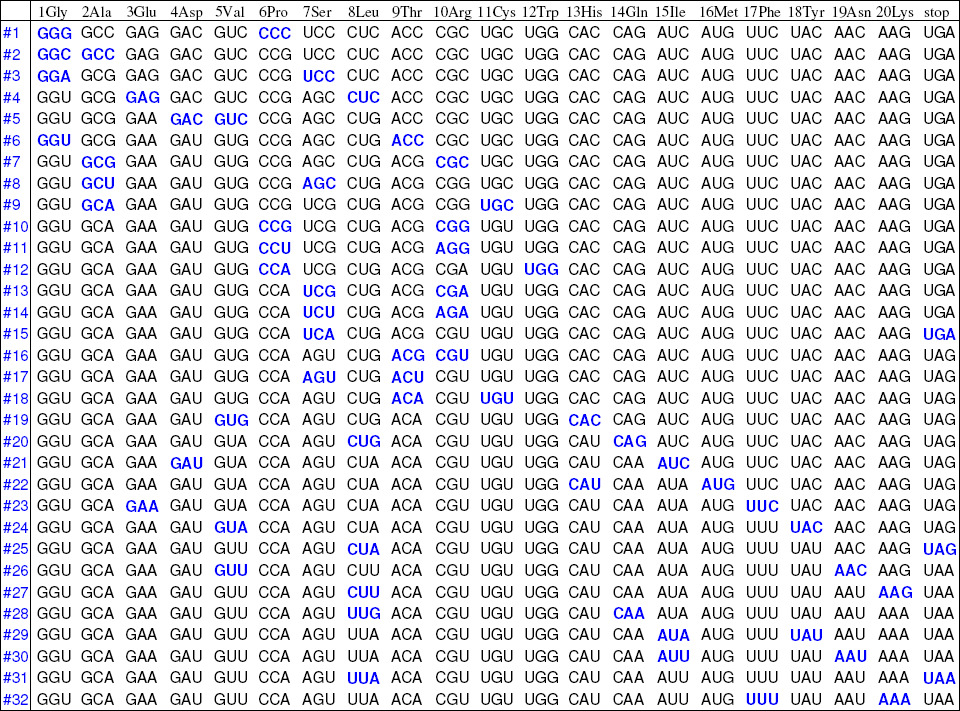
The recruitment orders of the codon pairs and the amino acids in the roadmap Fig 1A are listed in the left column and top row respectively. The codons that encode each amino acid are listed in columns respectively, where the row positions for the degenerate codons are determined by the corresponding codons in the first column of codon pairs from #1 to #32 (blue codons), and the other vacant positions in each column are filled by the nearby degenerate codons respectively. These blue codons are aligned from up-left to down-right, which shows that the later the codon pairs recruited, the later the corresponding amino acids recruited.

**Fig S2b2.**
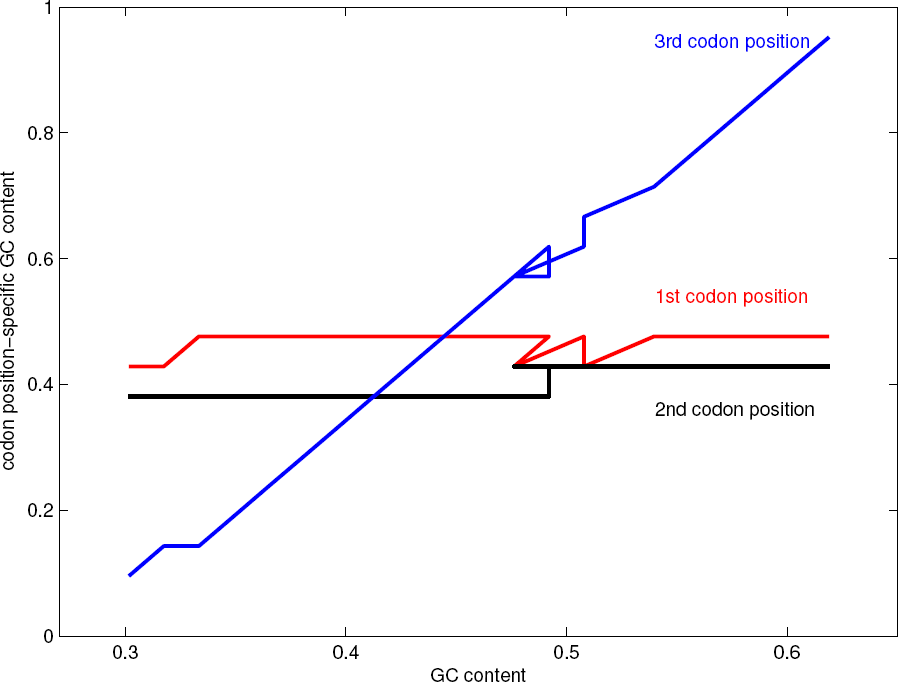
It is observed that the GC content in the third codon position decreases more rapidly than that in the first and second base positions when the total GC content decreases, and the GC content in the first codon position is generally greater than that in the second codon position. Such a pattern of variation of the base position specific GC contents in observations can be explained by the roadmap. According to the orders of degenerate codons for each amino acid in Fig S2b1, the total GC content and the GC contents at 1st-, 2nd-, 3rd-codon-position at stages from #1 to #32 can be obtained by counting and calculating in each row of Fig S2b1, respective. Consequently, the relationships between the total GC content and the base position specific GC contents are obtained respectively. Even some detailed features in observation are reproduced in the present figure based on the roadmap Fig 1A, such as the closer distance between 1st and 2nd codon position GC content at low total GC content side. Such an explanation indicates that the recruitment order of the codons from #1 to #32 obtained based on the roadmap Fig 1A is considerably reasonable, and hence confirms the validity of the roadmap theory.

**Fig S2b3.**
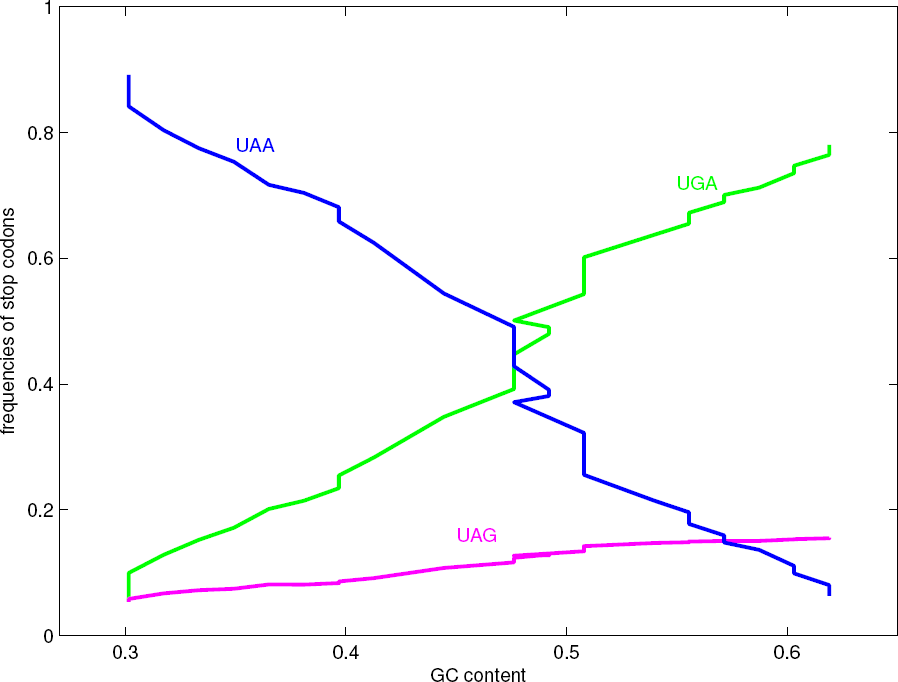
It is observed that the stop codons vary in certain pattern when the total GC content decreases based on genomic data. Namely, along the evolutionary direction of the decreasing total GC content, the first stop codon *UGA* in the roadmap decrease greatly, the second codon *U AG* in the roadmap decrease slightly, and the third stop codon *U AA* increase greatly. Such a pattern in observations can be explained by the roadmap. The relationships between the total GC content and the frequencies of the three stop codons are obtained based on the roadmap Fig 1A, respectively, by counting and calculating similarly in Fig S2b2, where the variation ranges for the three stop codons have been adjusted according to the observations. A detailed feature in observations, namely the respective downward and upward leaps of *UGA* and *U AA* at the middle total GC content, is reproduced in the present figure based on the counts in Fig S2b1. Such an explanation indicates that the recruitment order of the three stop codons obtained based on the roadmap Fig 1A is considerably reasonable, and hence confirms the validity of the roadmap theory.

**Fig S2b4.**
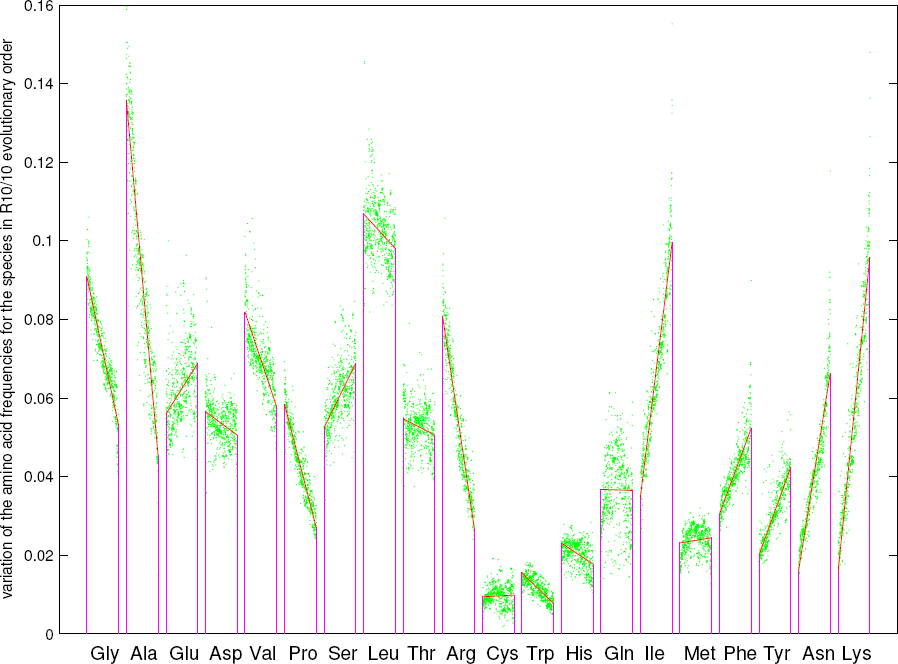
Explanation of the variation of the amino acid frequencies. The 20 amino acids are arranged in the recruitment order in the roadmap Fig 1A. The 20 amino acid frequencies for each of the 803 species are obtained respectively based on the genomic data in NCBI. And the 803 amino acid frequencies (green dots) for each of the 20 amino acids are all arranged properly in the *R*_10/10_ order, respectively. The variation trend of the amino acid frequencies for each of the 20 amino acids is obtained by the regression line (denoted in red). Generally speaking, the variation trends for the earlier amino acids tend to decrease, and the variation trends for the latecomers to increase (Fig S2b4).

**Fig S2b5.**
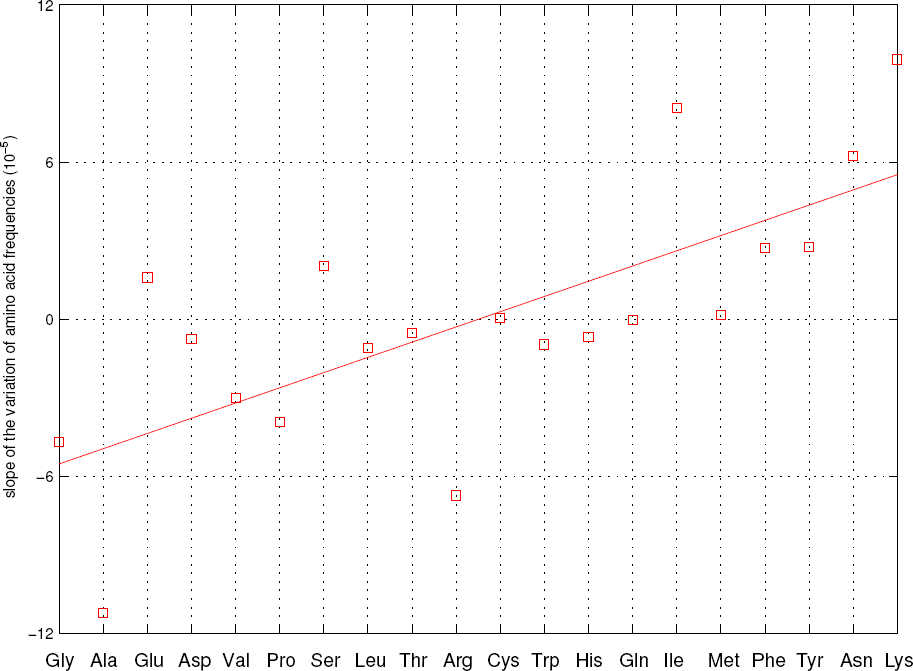
Increasing variation rates of the amino acid frequencies. The slope of the regression line of variation of amino acid frequencies for each amino acid is obtained based on the data in Fig S2b4. It is observed in the present figure that the slopes vary from negative to positive in general according to the recruitment order of amino acids in the roadmap. Such a natural result indicates that the recruitment order of the amino acids from *No*.1 to *No*.20 obtained based on the roadmap Fig 1A is considerably reasonable, and hence confirms the validity of the roadmap theory.

**Fig 2C.**
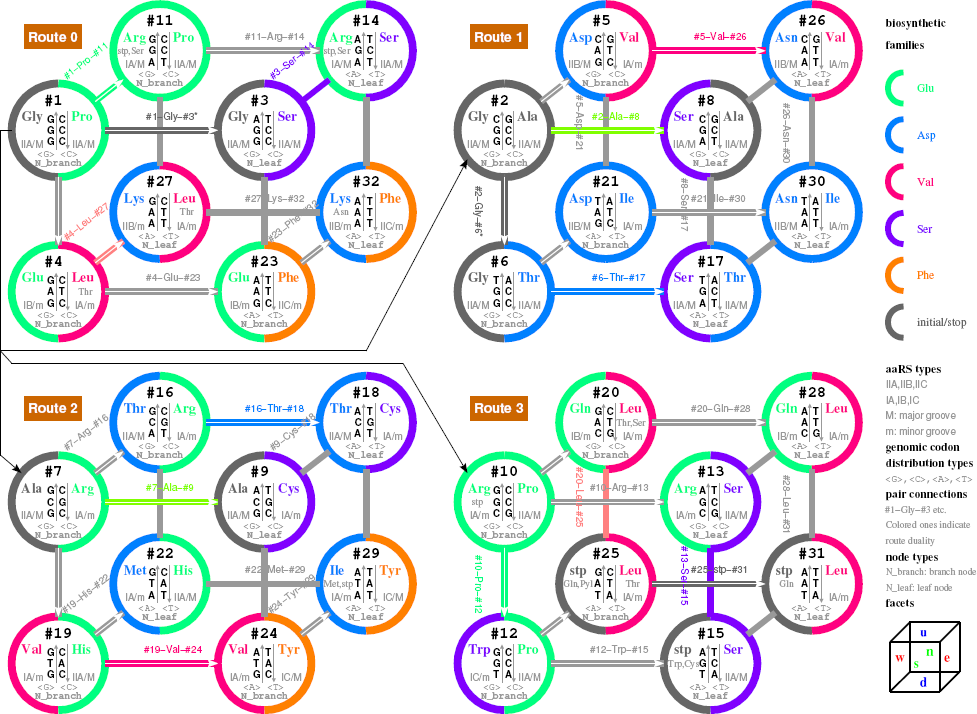
Explanation of the codon degeneracy based on the relationships among codons (pair connections and route dualities) in the roadmap. This is a revised plot of the roadmap Fig 1A to indicate the symmetry in the evolution of the genetic code, where the four routes are represented by four cubes respectively. Pair connections are marked besides the evolutionary arrows in the roadmap. Route dualities are indicated by same colours for the corresponding pair connections. The biosynthetic families of amino acids are denoted by coloured semicircles. The types of the aminoacyl-tRNA synthetases are besides the codons. Branch nodes and leaf nodes are distinguished. The 6 facets for each route are indicated on a cube at bottom right.

**Fig S2c1.**
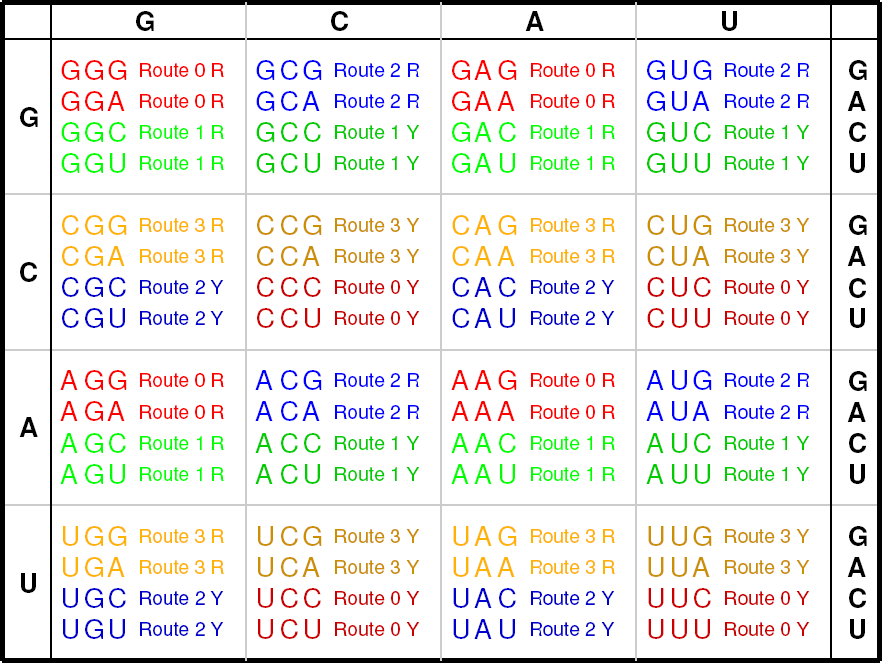
The distribution of codons from R- and Y-strands of *Route* 0−3 in the *GCAU* genetic code table. The pattern of the 4×4 codon boxes for the degenerate codons relates to such a distribution of the four routes, due to the evolution of the genetic code along the roadmap.

**Fig S2c2.**
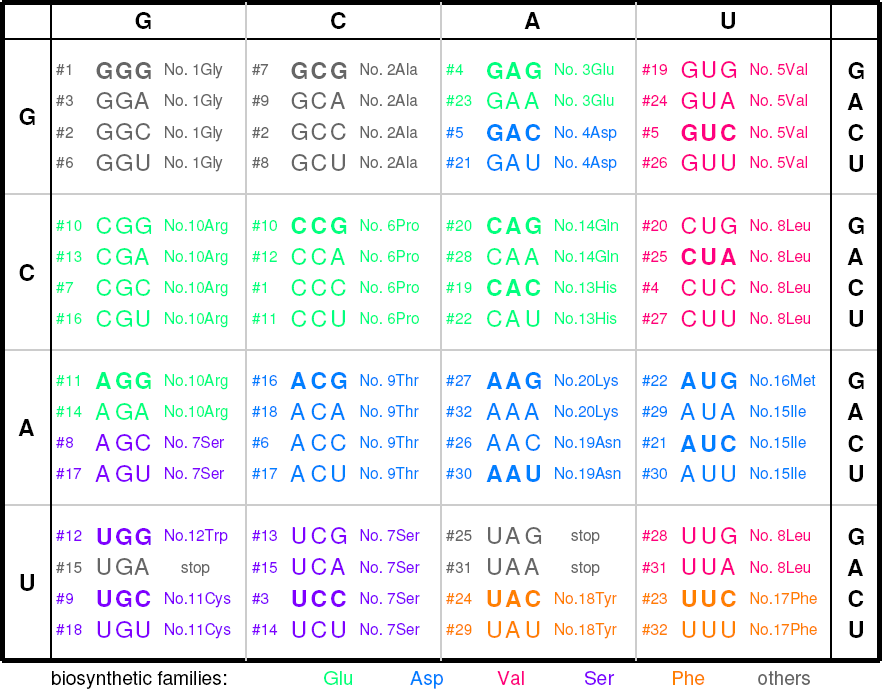
The clusterings of biosynthetic families (Glu Asp Val Ser Phe) in the *GCAU* genetic code table. Such nice clusterings are correspondingly observed in the R- and Y-strands of *Route* 0−3 in Fig 2C (denoted in the same group of colour as in the present figure). The clusterings of biosynthetic families in the present figure are closely related to the distribution of codons from R- and Y-strands of *Route* 0−3 in Fig S2c1, due to the recruitment of amino acids along the roadmap. Generally speaking, the amino acids are arranged properly in the recruitment order from *No*.1 to *No*.20 along the direction from *G*, *C* to *A*, *U* in the *GCAU* genetic code table.

**Fig S2c3.**
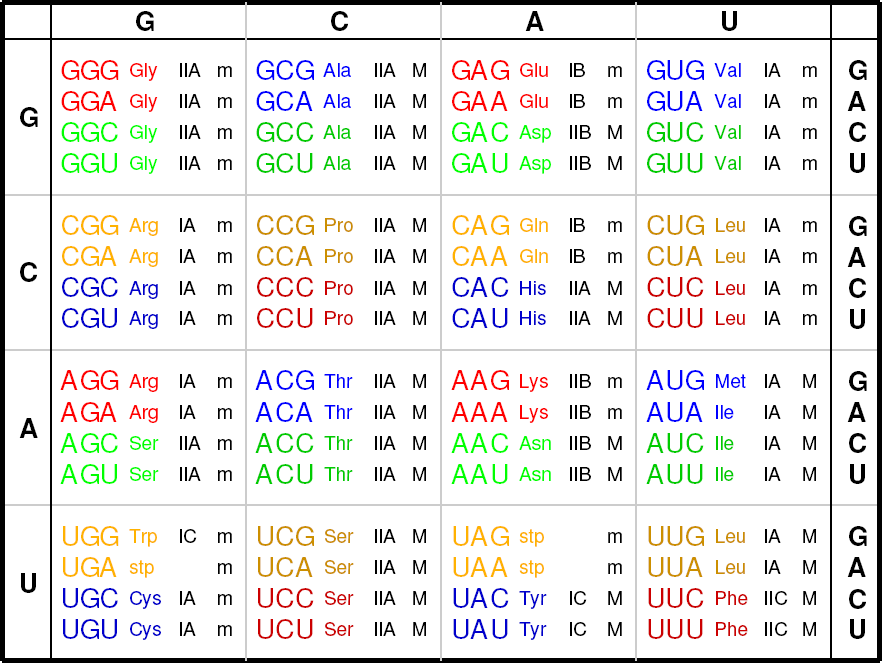
The distribution of types of aminoacyl-tRNA synthetases in the *GCAU* genetic code table. The aminoacyl-tRNA synthetases can be divided into *Class II* and *Class I*, which can be divided into subclasses *IIA*, *IIB*, *IIC*, and *IA*, *IB*, *IC*, respectively. And the aminoacyl-tRNA synthetases can also be divided into minor groove ones (*m*) and major groove ones (*M*).

**Fig 2D.**
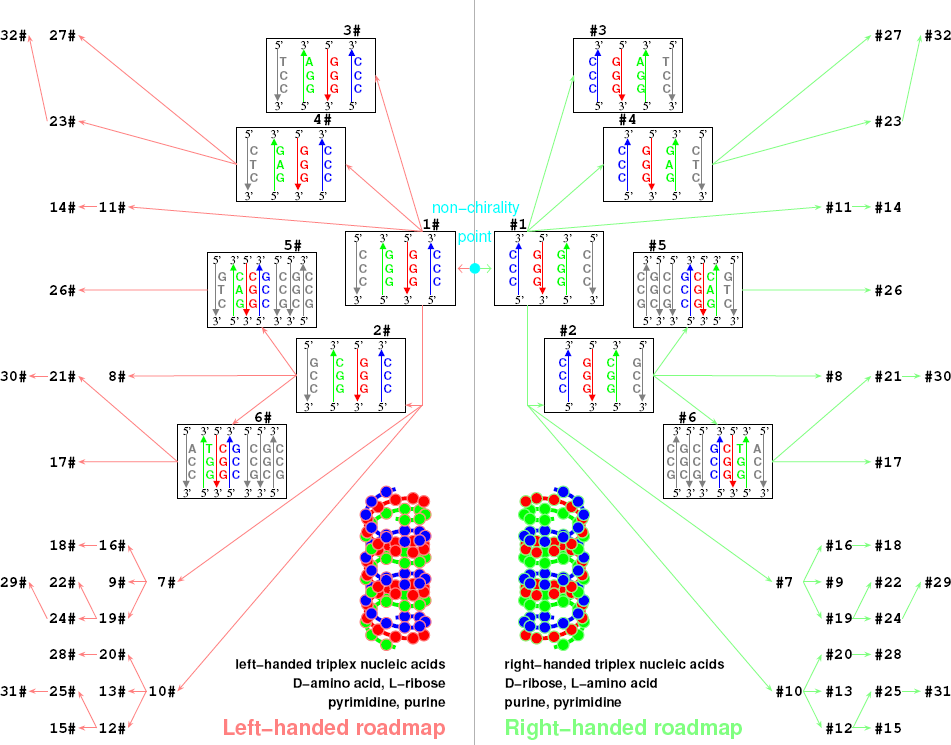
The homochirality of life originated in a winner-take-all game between the right-handed roadmap and the left-handed roadmap. The right handed roadmap is just the roadmap in Fig 1A (only the initial subset are shown explicitly here), whose chirally symmetric partner roadmap, i. e., the left-handed roadmap, is plotted on the left. Ref to Fig S2d1 to see a heuristic model on the origin of the homochirality of life.

**Fig S2d1.**
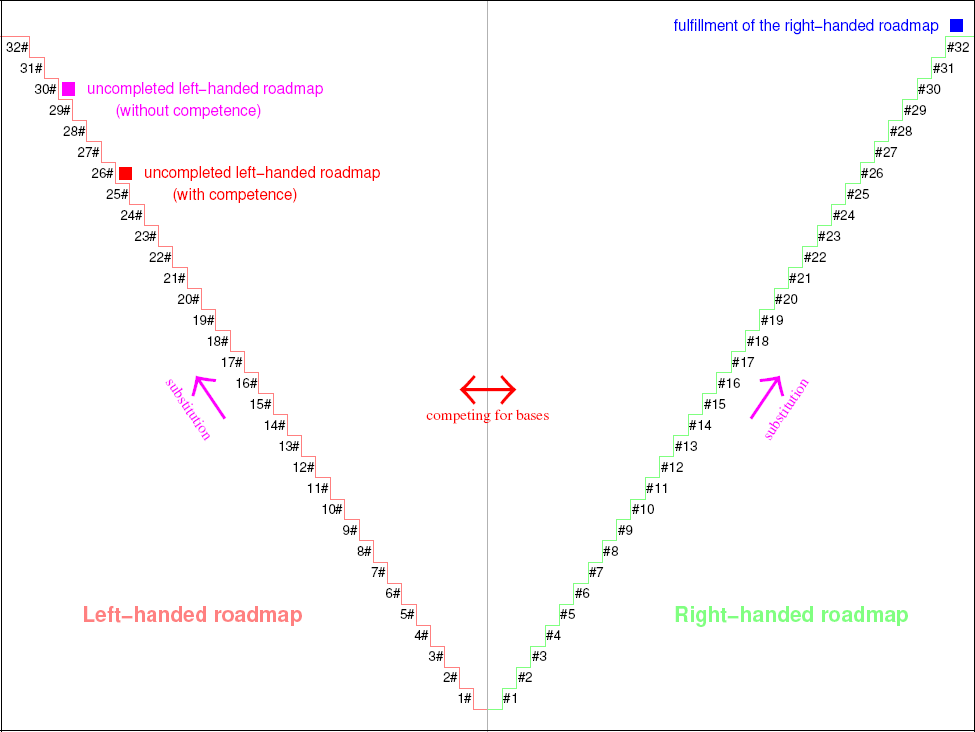
Simulation of the origin of homochirality of life. The roadmap in Fig 1A is a right-handed roadmap, whose chirally symmetric counterpart, i. e. the left-handed roadmap, can also be achieved in theory (Fig 2D). In this simulation, a constant substitution rate is assumed for the substitutions from #1 to #32 for the right-handed roadmap and from 1# to 32# for the left-handed roadmap. Which chirality to be selected is absolutely by chance. The programme can be run for many times. The two chiral roadmaps are obtained equally. When any chiral roadmap is fulfilled (such as the right-handed roadmap in the present figure), the chirally opposite roadmap may reach a certain uncompleted stage. The simulation indicates that the competition for the non-chiral bases between the two chiral roadmaps may enlarge the gap between the fulfilled chiral roadmap and the uncompleted chiral roadmap.

**Fig 3A.**
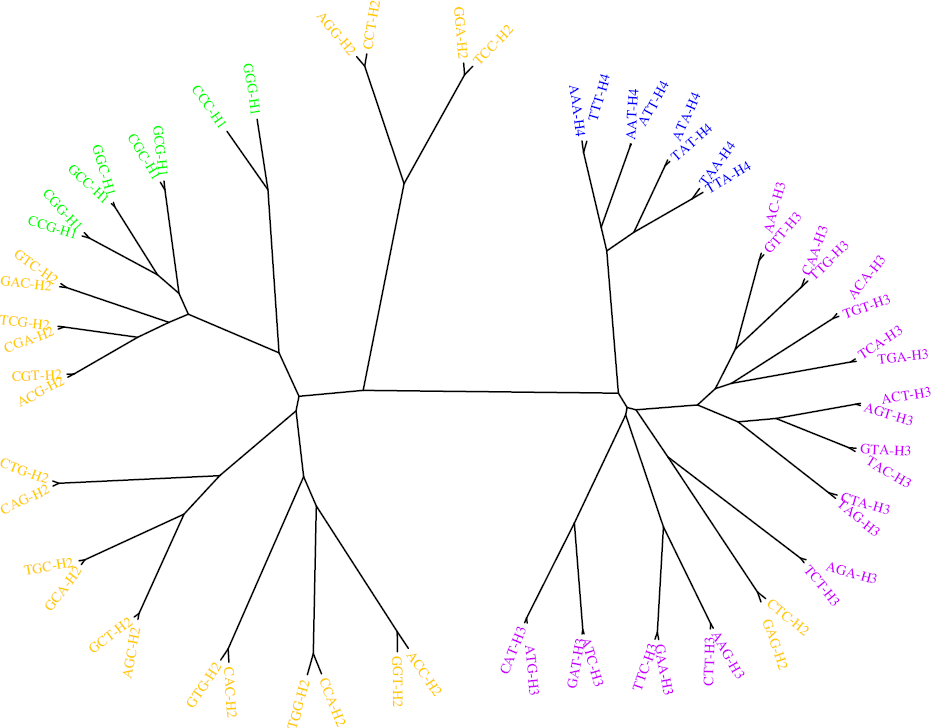
The tree of codons based on the genomic codon distributions, which agrees with the differentiation of the four hierarchies *Hierarchy* 1−4 (H1−4) in the roadmap. This tree of codons is plotted based on the distance matrix for codons by averaging the correlation coefficients of genomic codon distributions among the species. Both the tree of codons in this figure and the tree of species in Fig S3a5 are obtained based on a same set of genomic codon distributions.

**Fig S3a1.**
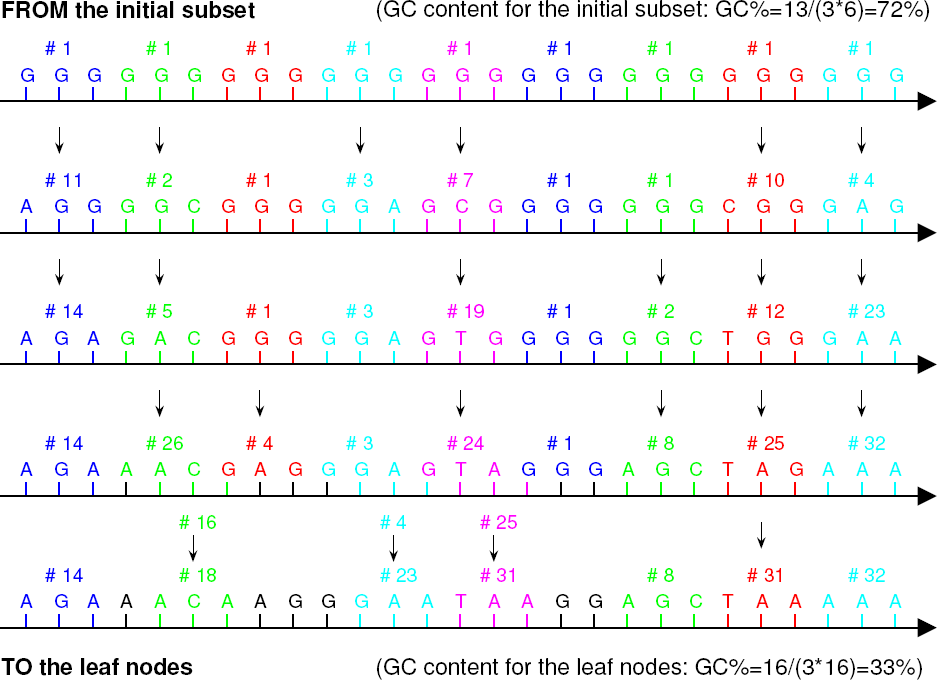
The base substitutions along the roadmap may occur randomly at any places in a strand according to the roadmap Fig 1A. For a simple example, starting from *Poly G* in the present figure, the strand may evolve randomly step by step according to the base substitutions in the triplex DNA along the roadmap. After several steps, a certain sequence of strand is achieved. Such base substitutions essentially determine the genomic codon distributions for species (Fig S3a3). The GC content for the initial subset (Fig 1A) is about 72%, and the GC content for the leaf nodes about 33%. This range of GC content generally agrees with the range of GC content in observations (Fig S3a2).

**Fig S3a2.**
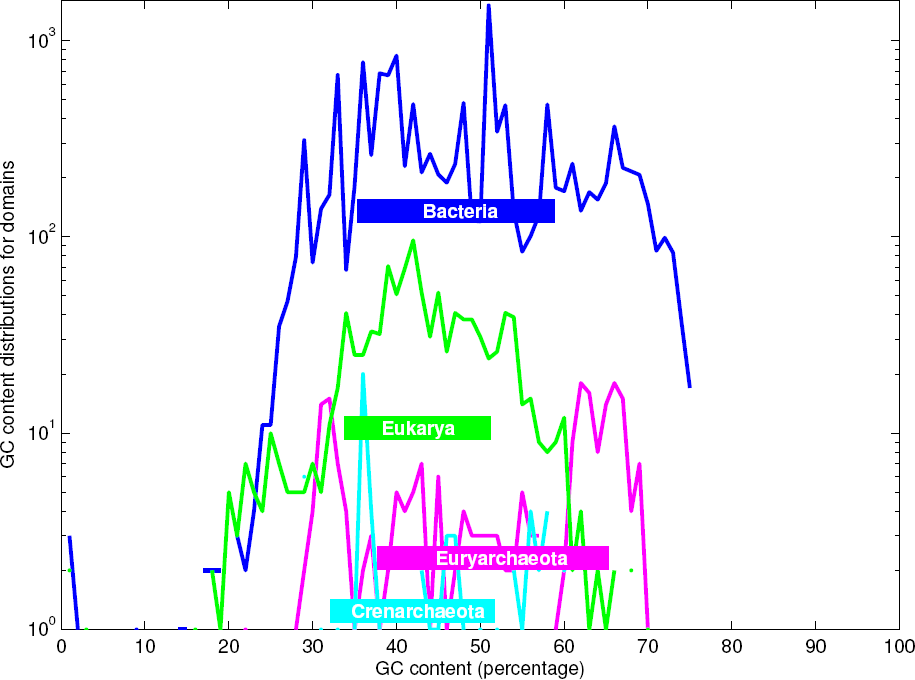
GC content ranges for taxa, which can be explained by the evolution of sequences based on the roadmap (Fig S3a1). The origin order of domains (obtained in Fig 3C) is irrelevant to the order of the average GC contents in taxa here.

**Fig S3a3.**
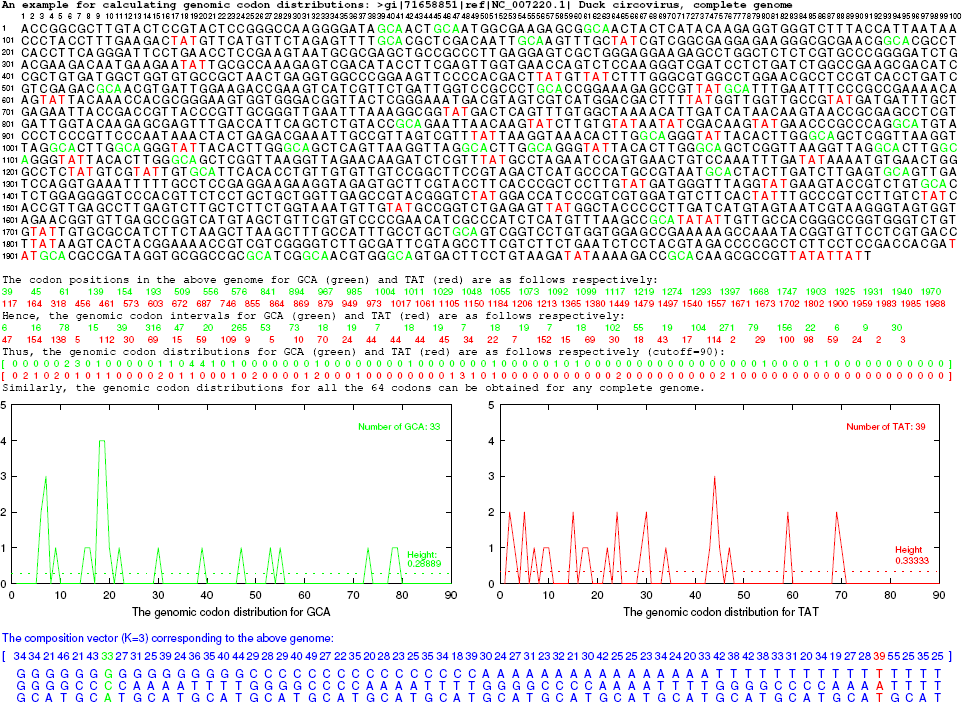
Definition of the genomic codon distribution. Take the complete genome of Duck circovirus for example, and take the codons *GCA* (denoted in green) and *T AT* (denoted in red) among the 64 codons for example. First, mark all the positions of *GCA* and *T AT* in the genome respectively; second, calculate the codon intervals for *GCA* and *T AT* respectively, and obtain the distributions of intervals of *GCA* and *T AT* respectively, where a cutoff 90 is used in this simple case when counting the number of common intervals (a sufficiently large enough cutoff 1000 is used in this paper); and third, obtain the genomic codon distributions for *GCA* and *T AT* by normalising the distributions of intervals of *GCA* and *T AT* respectively. The genomic codon distribution for a species consists of 64 distributions for each of the 64 codons. An example of unnormalised genomic codon distribution (take for example only two distributions for *GCA* and *T AT*) is illustrated in two subfigures of the present figure. Four normalised genomic codon distributions for species in different domains are respectively shown in Fig S3b2. In addition, the summations of the elements of the unnormalised genomic codon distributions (e. g. 33 and 39 for *GCA* and *T AT* respectively) are obtained; consequently the average height of the genomic codon distribution is obtained for each of the distributions of the 64 codons respectively. In a so-called composition vector tree method in the molecular phylogenetics, the elements of the *K* = 3 composition vector for a species (e. g. the elements 33 and 39 for *GCA* and *T AT*) correspond to the summations of the elements of the genomic cordon distributions respectively, which are also proportional to the average heights of the genomic codon distribution.

**Fig S3a4.**
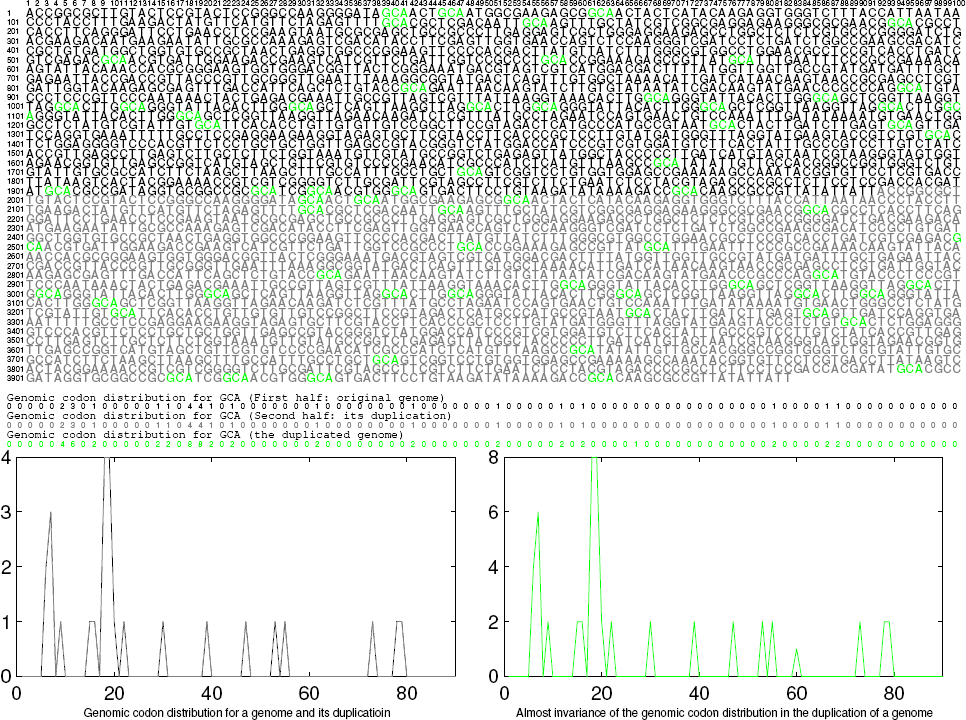
Invariance of the genomic codon distribution after genome duplication. If a genome expands by duplicating itself, the genomic codon distribution does not change due to the normalisation. In the present figure, the distribution in the left subfigure and the distribution in the right subfigure will be same after normalisation. In this sense, the genomic codon distribution is a conservative and intrinsic feature of a species when its genome evolves generally by large scale duplications (Fig S4a2). The trees of species in Fig S3a5 and Fig S3d8 are obtained based on the genomic codon distributions of species with large variation of genome sizes, where the genome sizes themselves do not play a significant role also due to the normalisation in the definition of the genomic codon distribution.

**Fig S3a5.**
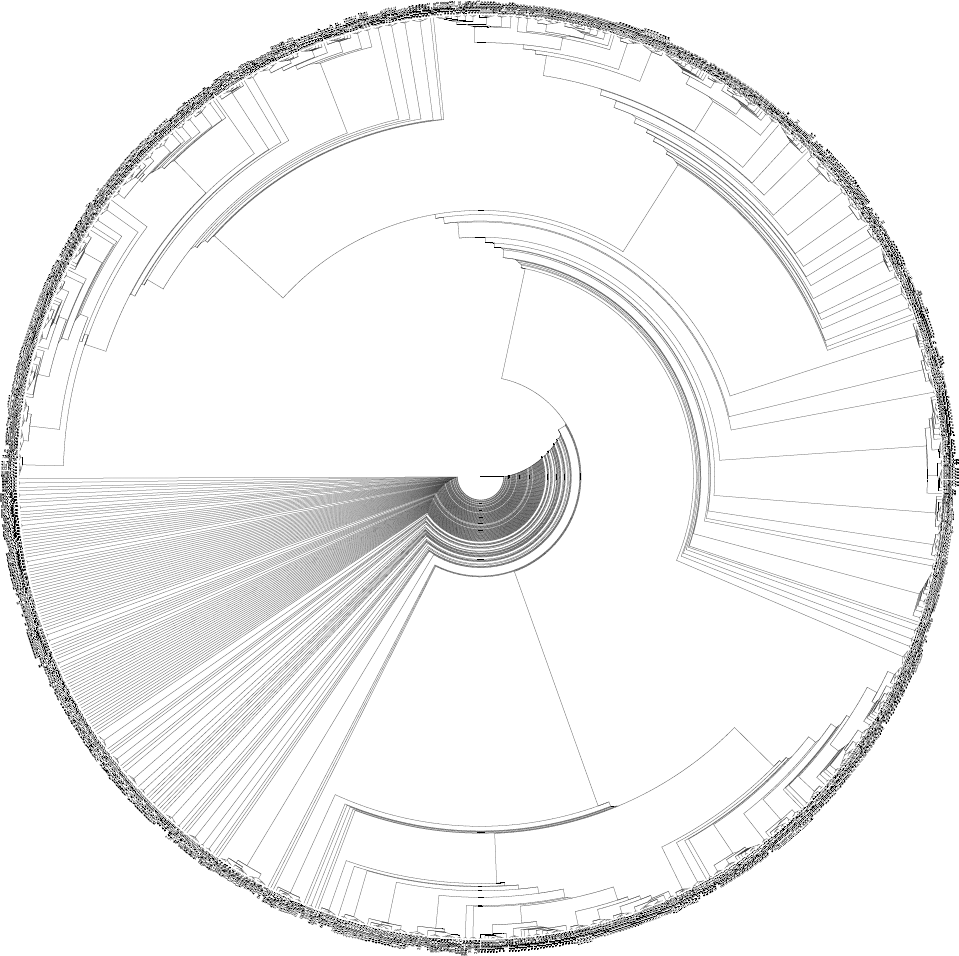
Tree of species according to the average correlation coefficients among genomic codon distributions by averaging for the 64 codons. This tree is reasonable to some extent. Considering the also reasonable tree of codons in Fig 3A that is obtained based on the same data, it is inferred that an intrinsic relationship exists between the tree of life and the evolution of the genetic code. Please enlarge to see details. Refer to the legend of Fig 3D in the appendix to see the abbreviations of the taxa.

**Fig S3a6.**
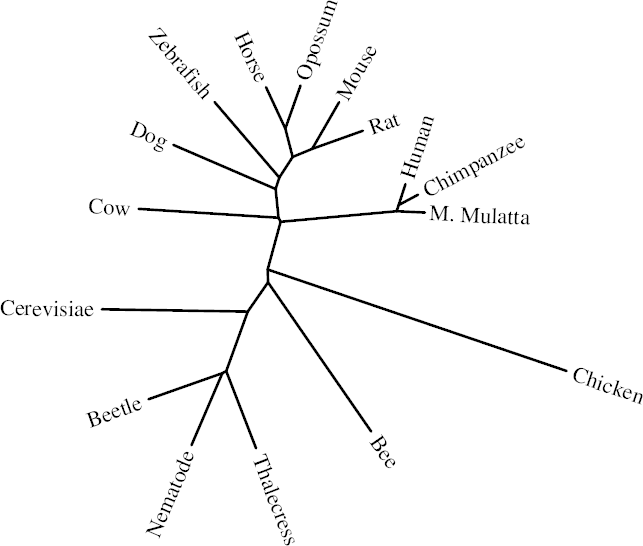
Tree of eukaryotes based on the average correlation coefficients among genomic codon distributions. This tree is reasonable to some extent. Note that most eukaryotic genomes are non-coding DNA. So this reasonable tree is mostly based on the non-coding DNA, which indicates that the non-coding DNA, rather than the coding DNA, plays a principal role in the phylogeny of eukaryotes.

**Fig 3B.**
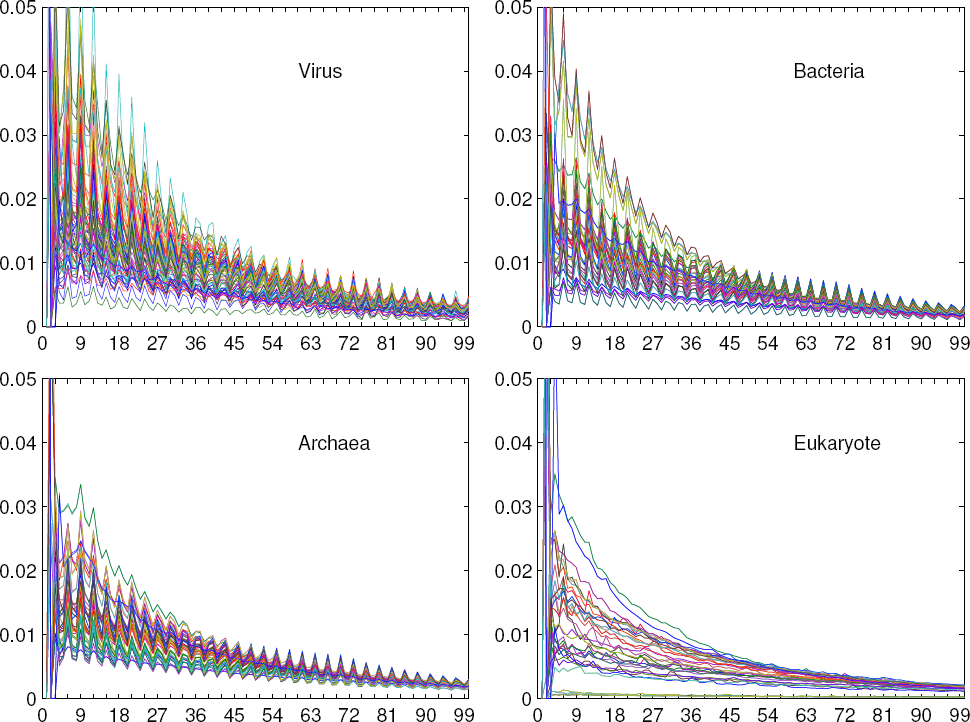
Features of genomic codon distributions for domains based on complete genome sequences. Refer to Fig S3b1 to see the genomic codon distributions of Crenarchaeota and Euryarchaeota; refer to Fig S3b2 to see the genomic codon distributions for species; and refer to Fig S3b3 to see the simulations of the genomic codon distributions.

**Fig S3b1.**
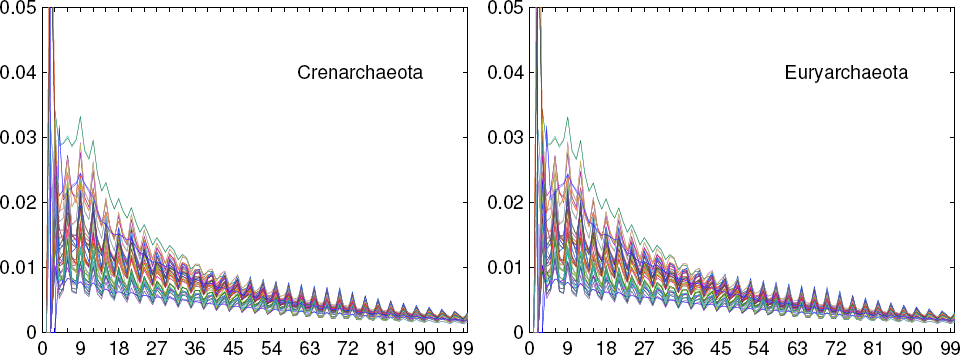
Genomic codon distributions for Crenarchaeota and Euryarchaeota based on genomic data. The features for both Crenarchaeota and Euryarchaeota agree with that for Archaea in Fig 3B, while they are quite different from the features for Eukarya in Fig 3B. So the relationship between Crenarchaeota and Euryarchaeota is closer than with Eukarya, which tends to agree with the three-domain tree rather than the eocyte tree.

**Fig S3b2.**
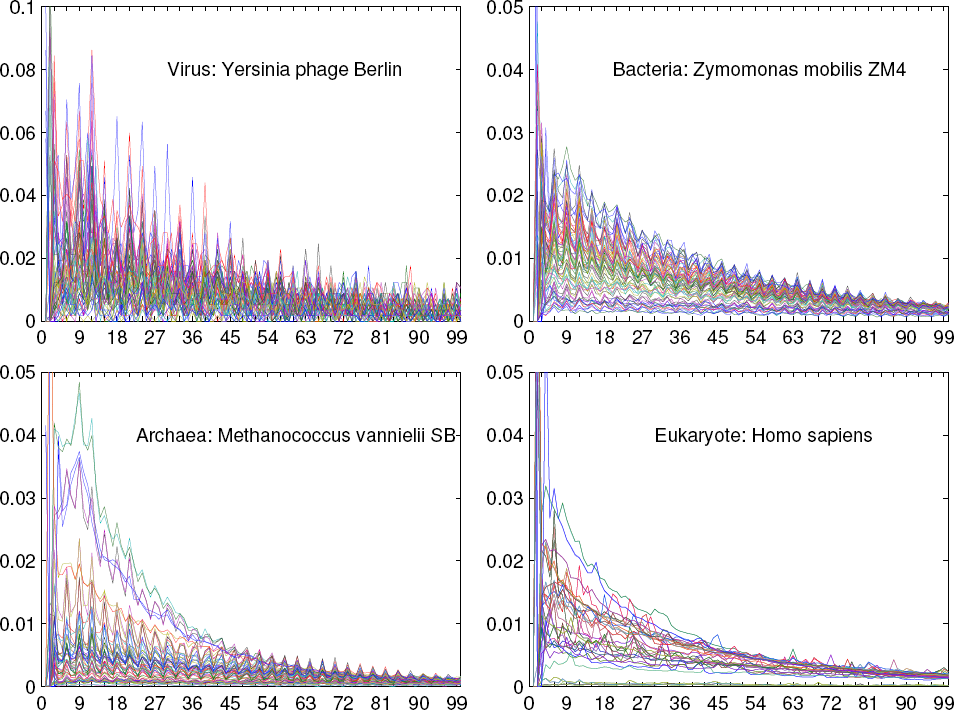
Features of genomic codon distributions of species in different domains based on genomic data. The features for species in the present figures agree with the features for the corresponding domains in Fig 3B respectively. For example, the features for eukaryotes are quite different from the features for the other domains, either for any species in the present figure or for the domain in Fig 3B.

**Fig S3b3.**
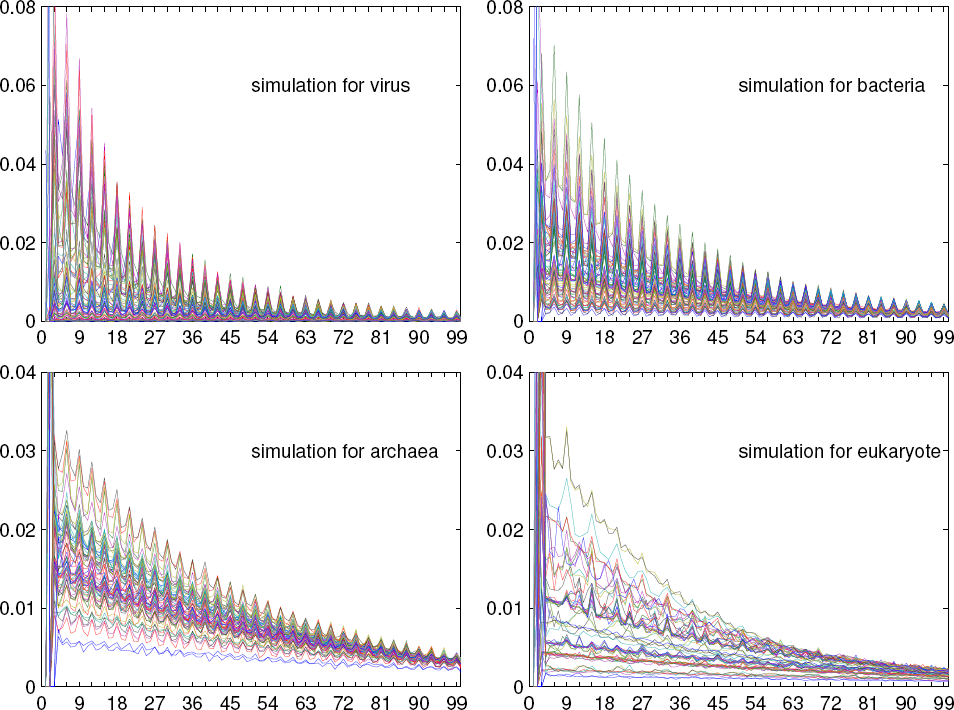
Simulations of features of genomic codon distributions for the different domains. This simulation results for the domains agree with the observations based on genomic data in Fig 2B, respectively. The simulations also reveal the mechanism that results in such features.

**Fig 3C.**
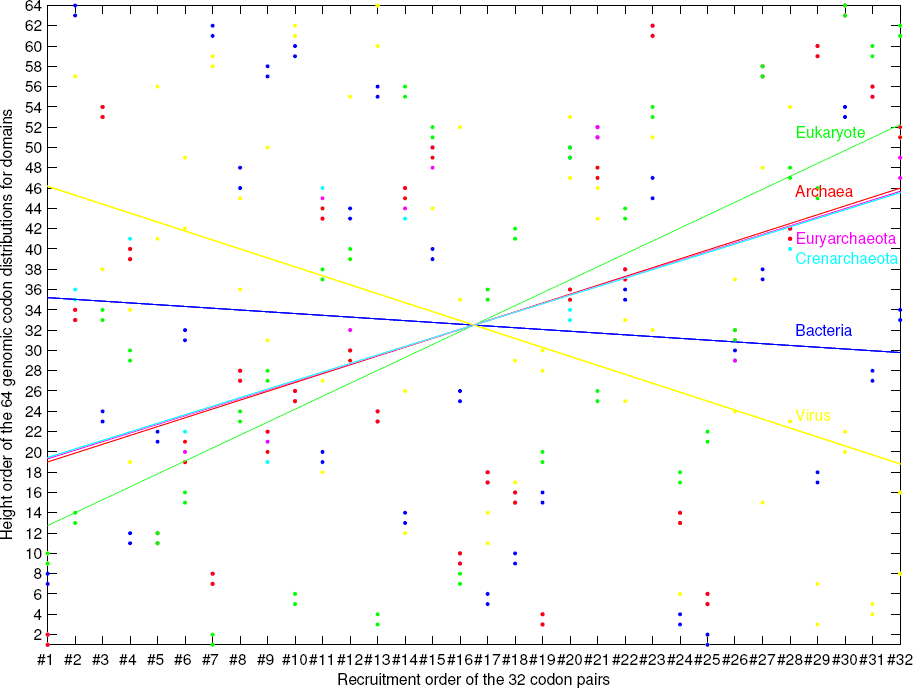
On the origin order of the domains. The recruitment order of the 32 codon pairs indicates an evolutionary direction, which reveals the origin order of the domains.

**Fig S3c1.**
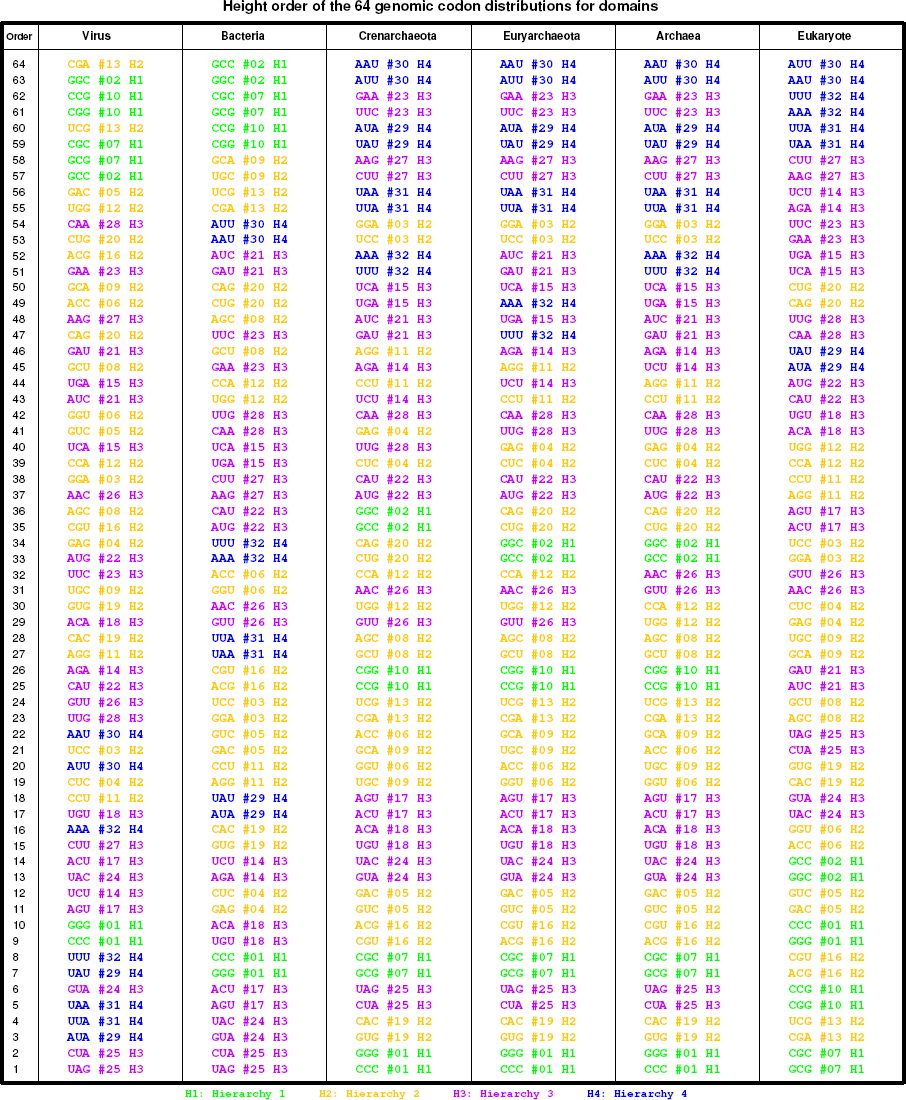
The orders of the average distribution heights for different domains, where the values of genomic codon distributions here are obtained based on genomic data in Fig 3B and Fig S3b1. The orders of the average distribution heights for Archaea and Eukarya are more similar with respect to *Hierarchy* 1−4 than that for Virus and Bacteria.

**Fig S3c2.**
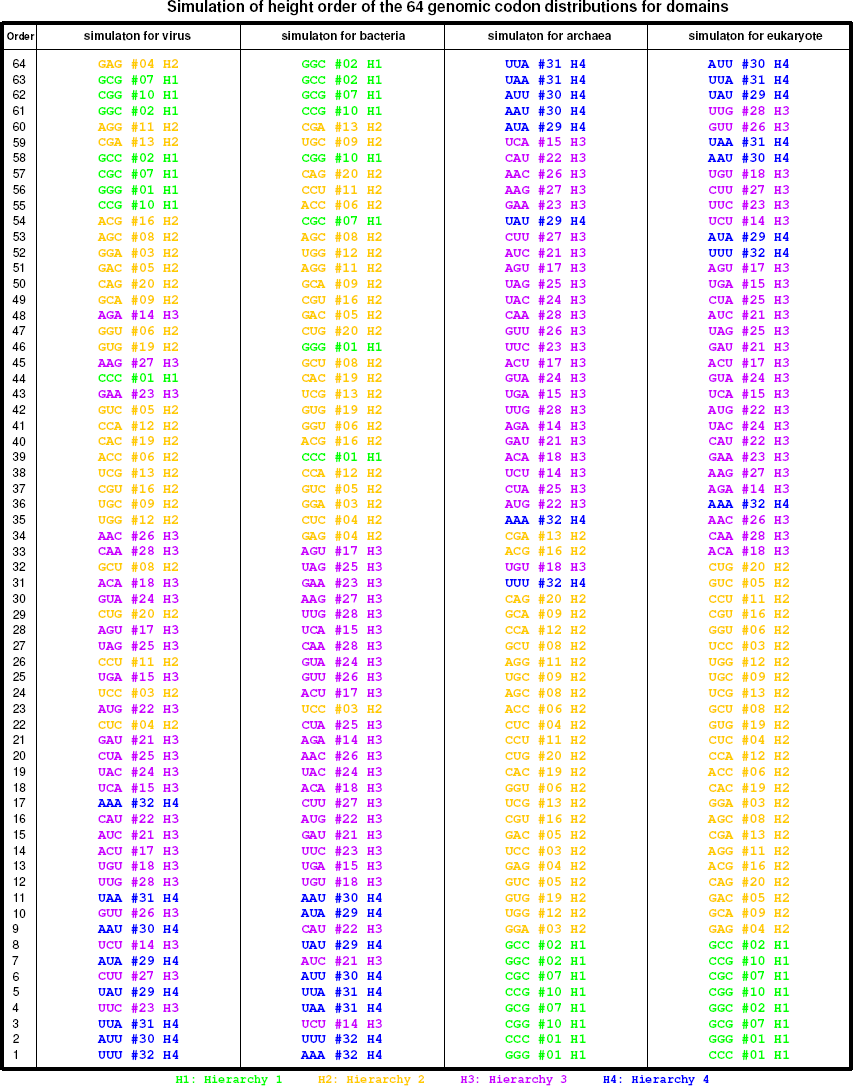
Simulations of the orders of the average distribution heights for different domains, where the values of genomic codon distributions here are obtained from the simulation results in Fig S3b3. These simulation results for the domains agree with the observations in Fig S3c1 based on genomic data, respectively.

**Fig 3D.**
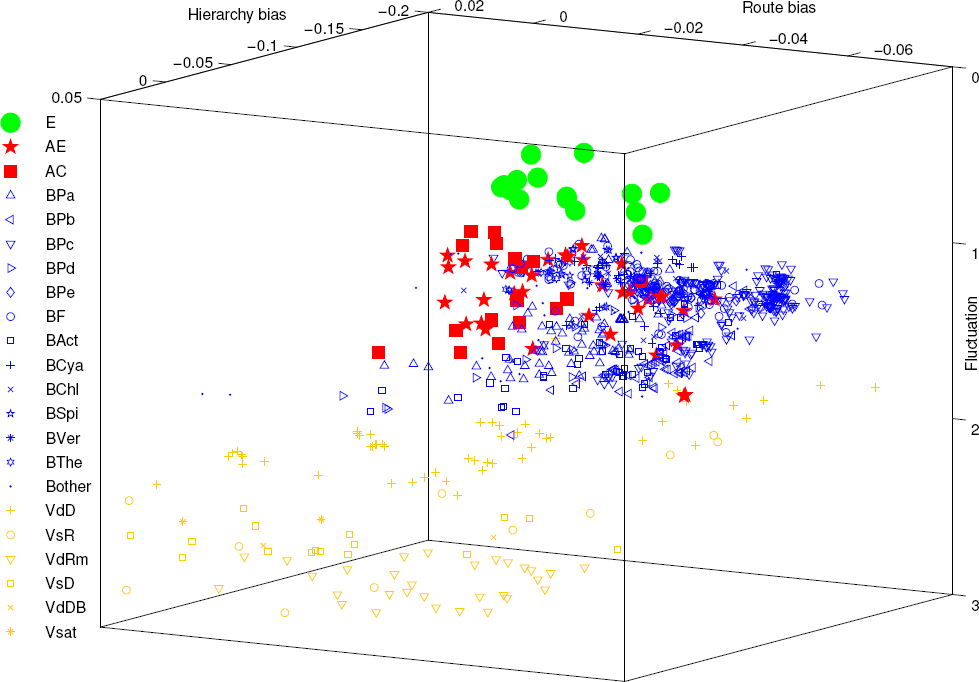
Classification of Virus (yellow), Bacteria (blue), Archaea (red) and Eukarya (green) in the evolution space, which is based on the roadmap and the complete genome sequences. Refer to Fig S3d3, Fig S3d4, S3d5 to see the two dimensional projections of this figure. The abbreviations for the taxa are as follows: Eukarya (E), Euryarchaeota (AE), Crenarchaeota (AC), Alphaproteobacteria (BPa), Betaproteobacteria (BPb), Gammaproteobacteria (BPc), Deltaproteobacteria (BPd), Epsilonproteobacteria (BPe), Firmicutes (BF), Actinobacteria (BAct), Cyanobacteria (BCya), Chlorobi (BChl), Spirochaetes (BSpi), Chlamydiae/Verrucomicrobia (BVer), Thermotogae (BThe), other bacteria (Bother), dsDNA virus (VdD), ssRNA virus (VsR), Mammalian orthoreovirus (VdRm), ssDNA virus (VsD), Blainvillea yellow spot virus (VdDB), satellite virus (Vsat).

**Fig S3d1.**
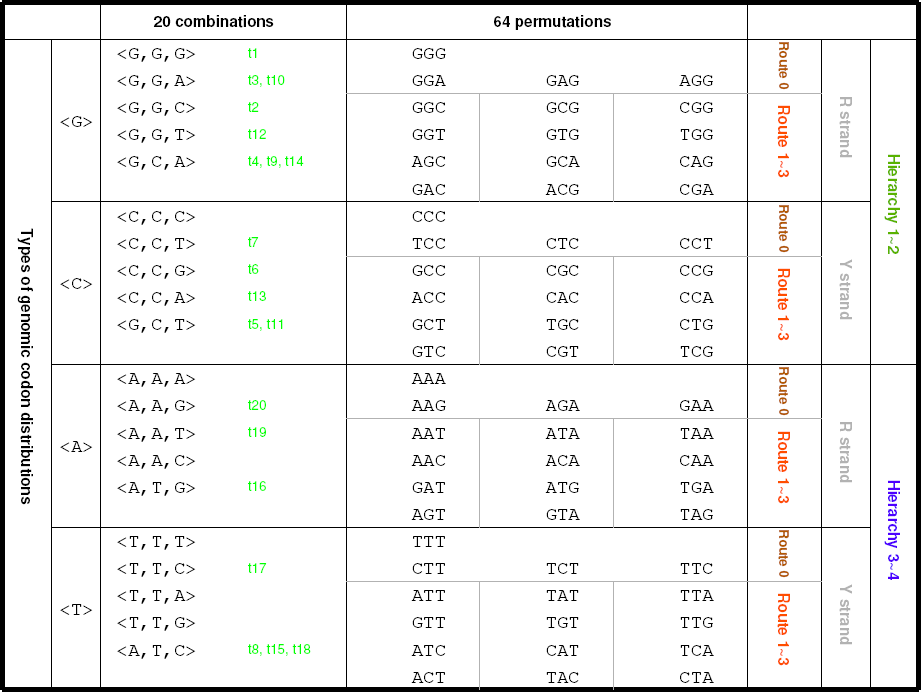
The 20 types of genomic codon distributions. There are respectively 20 combinations and 64 permutations of the 4 bases. There is a coarse correlation between the 20 combinations of codons and the 20 amino acids according to the tRNAs *t*1 to *t*20 (Fig S2a1).

**Fig S3d2.**
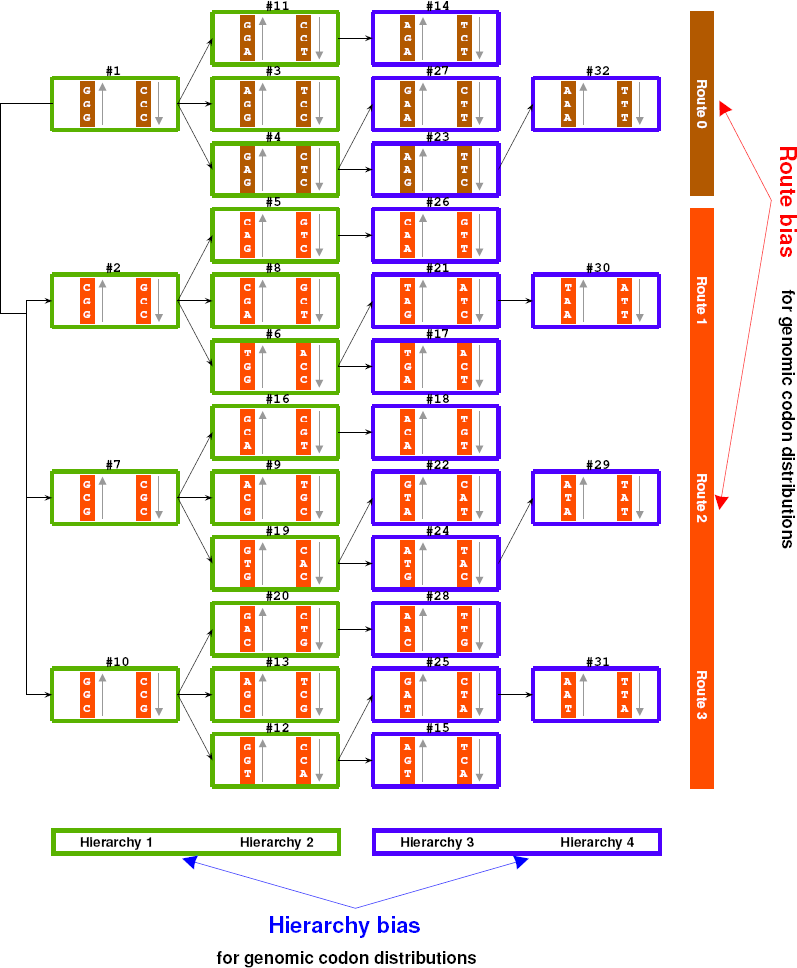
Explanation of the definitions of hierarchy bias and route bias. The hierarchy bias is defined by the difference between the averages of genomic codon distributions for *Hierarchy* 1 − 2 and that for *Hierarchy* 3 − 4. And the route bias is defined by the difference between the averages of genomic codon distributions for *Route* 0 and that for *Route* 1 − 3. By the way, the leaf node bias is defined by the difference between the averages of genomic codon distributions for the leaf nodes in *Route* 0 − 1 and that for the leaf nodes in *Route* 2 − 3.

**Fig S3d3.**
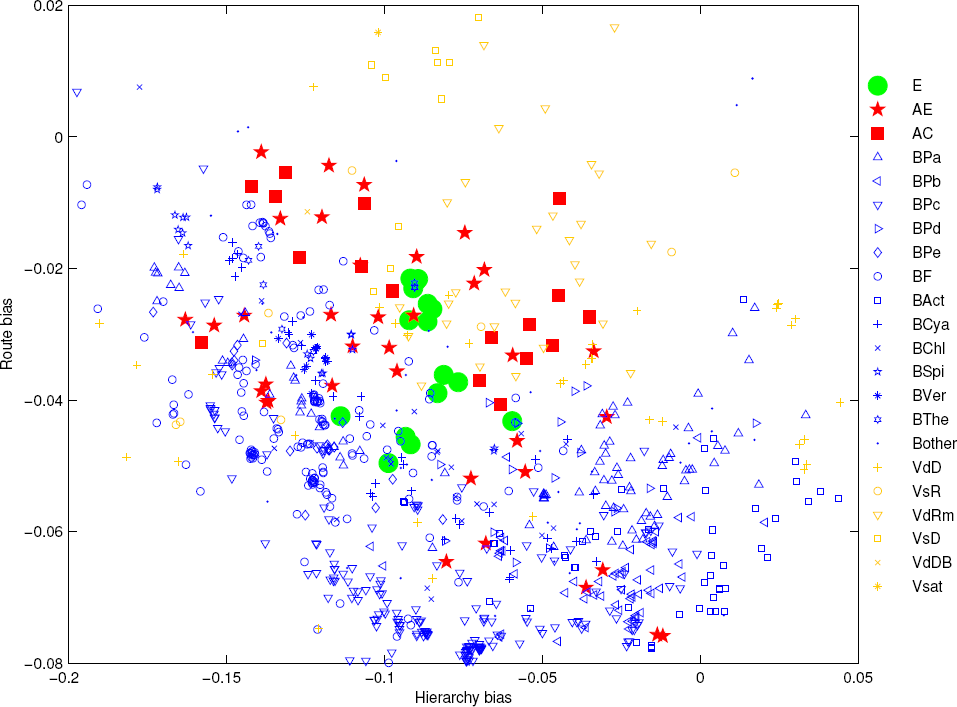
A two dimensional projection of the three dimensional evolution space (Fig 3D): the Hierarchy bias and Route bias plane. Bacteria (blue) and Archaea (red) are distinguished by and large in the present figure.

**Fig S3d4.**
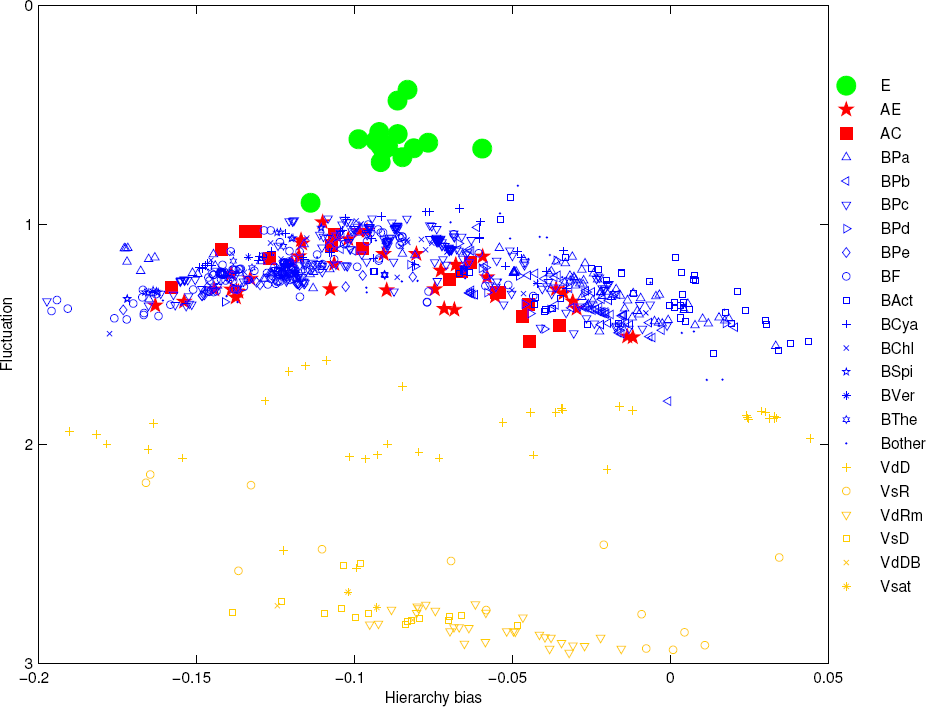
A two dimensional projection of the three dimensional evolution space (Fig 3D): the Hierarchy bias and Genomic codon distribution Fluctuation plane. Eukarya (green), Bacteria/Archaea, and Virus (yellow) are distinguished by and large in the present figure.

**Fig S3d5.**
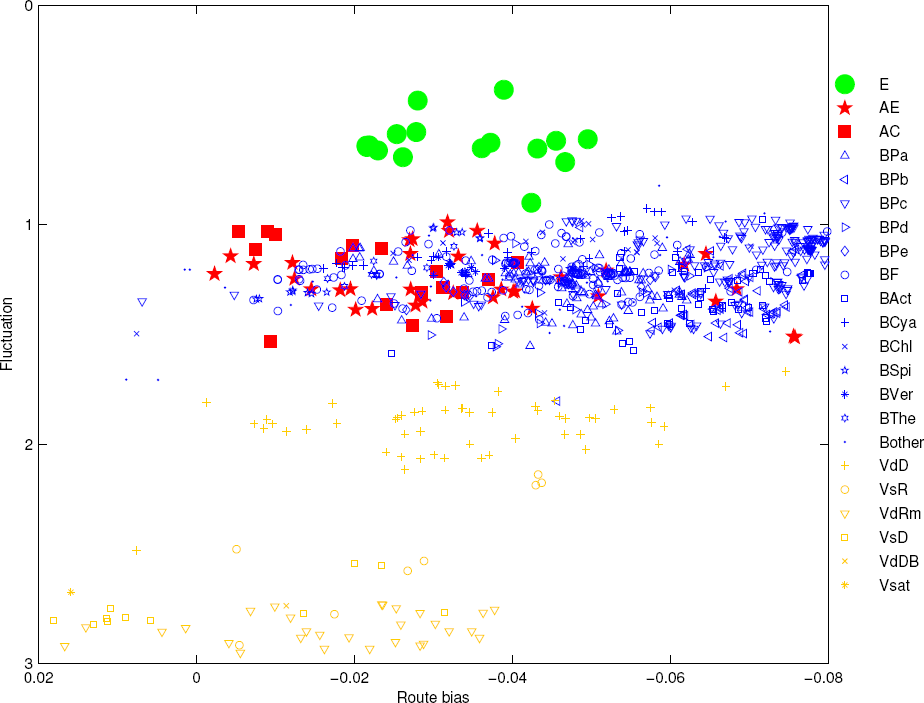
A two dimensional projection of the three dimensional evolution space (Fig 3D): the Route bias and Fluctuation plane. Eukarya (green), Bacteria (blue), Archaea (red), and Virus (yellow) are distinguished by and large in the present figure.

**Fig S3d6.**
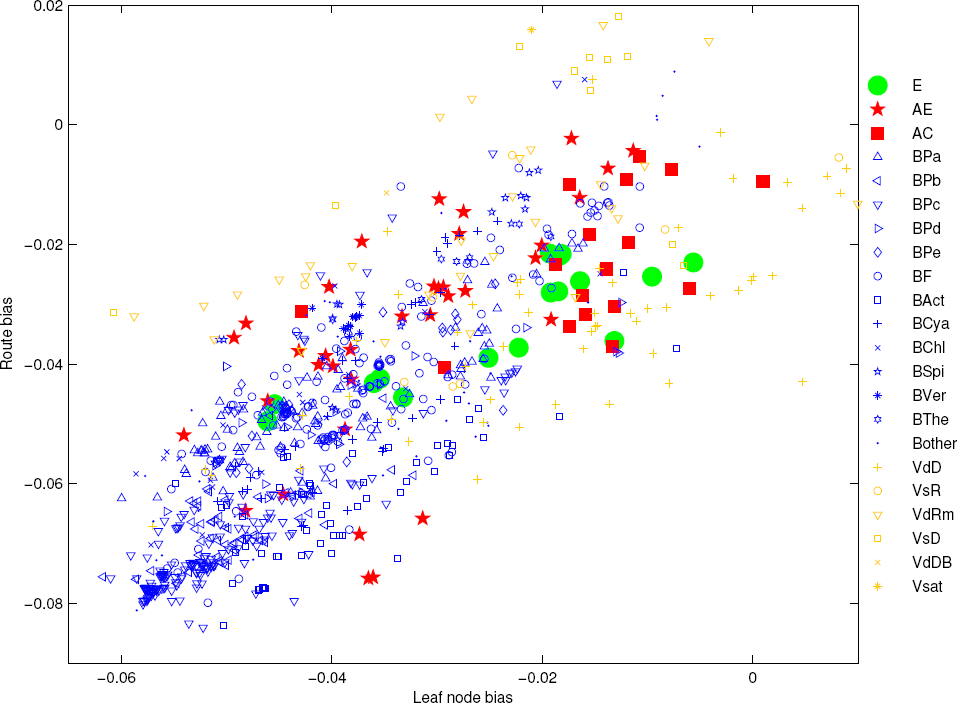
Distinguish Crenarchaeota (AC) and Euryarchaeota (AE) in the Leaf node bias and Route bias plane.

**Fig S3d7.**
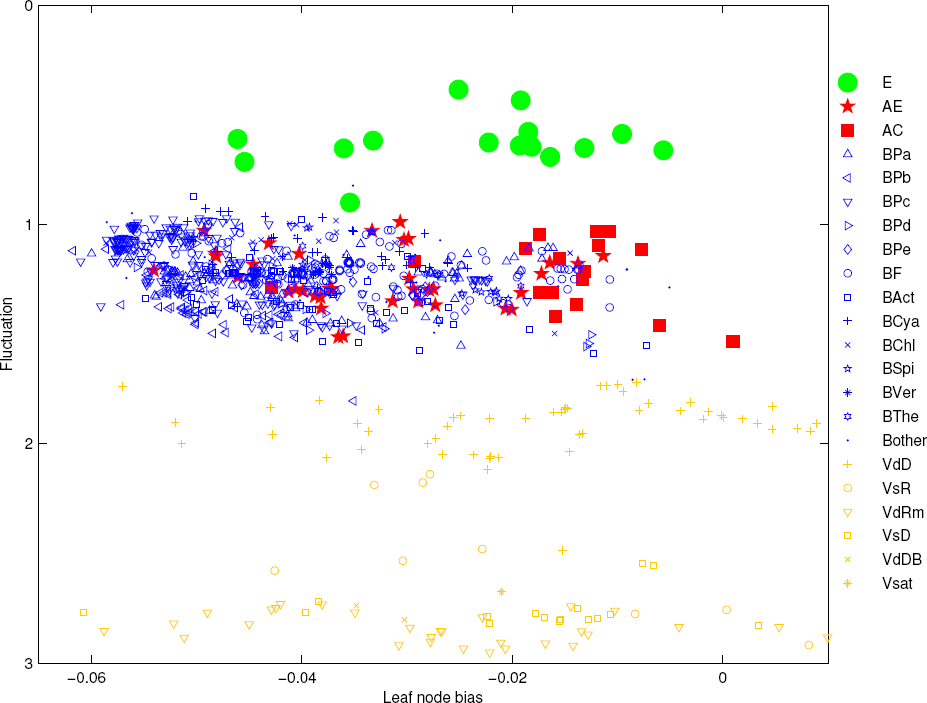
Distinguish Crenarchaeota (AC) and Euryarchaeota (AE) in the Leaf node bias and Fluctuation plane.

**Fig S3d8.**
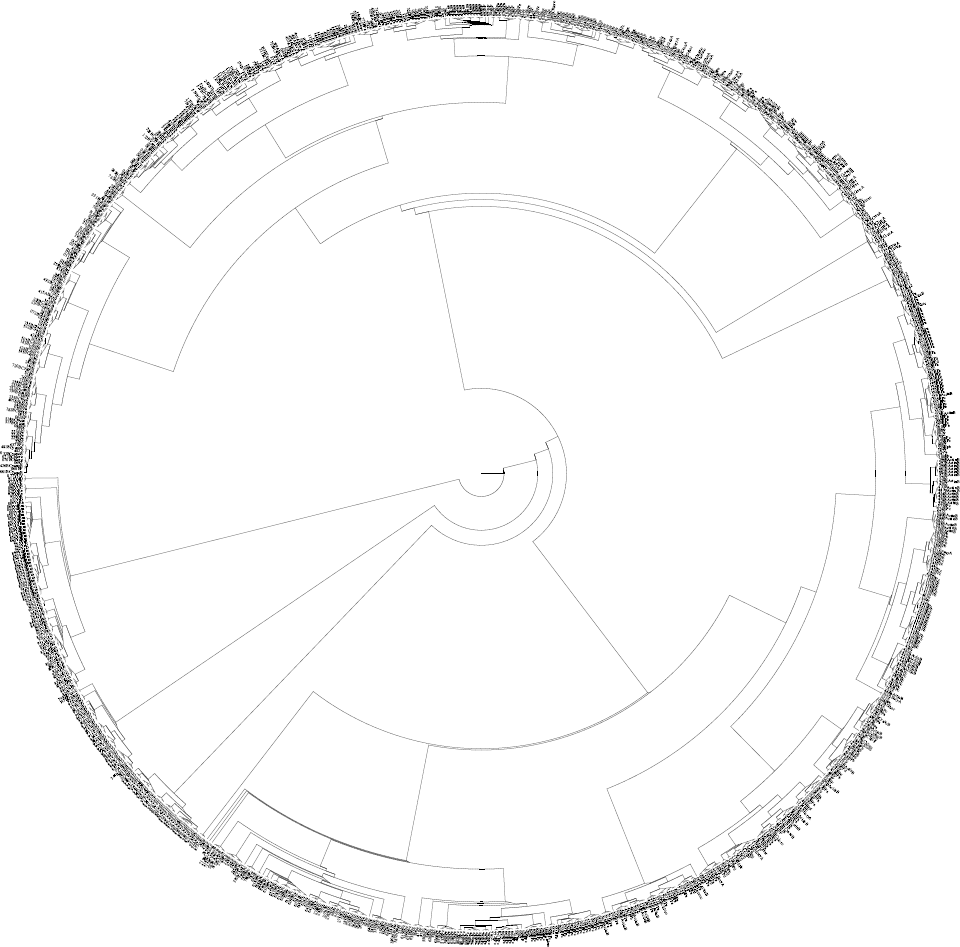
Tree of species based on the distances among species in the nondimentionalized evolution space Fig 3D. This tree agrees with the three-domain tree by and large. Please enlarge to see details. Refer to the legend of Fig 3D in the appendix to see the abbreviations of the taxa.

**Fig S3d9.**
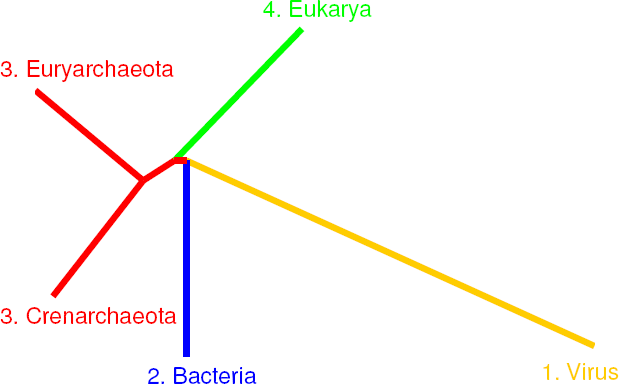
Tree of Bacteria, Crenarchaeota, Euryarchaeota, Eukarya and Virus based on the average distances among these taxa in the nondimentionalized evolution space Fig 3D. The origin order of these taxa is obtained based on the result in Fig 3C. This tree agrees with the three-domain tree rather than the eocyte tree.

**Fig S3d10.**
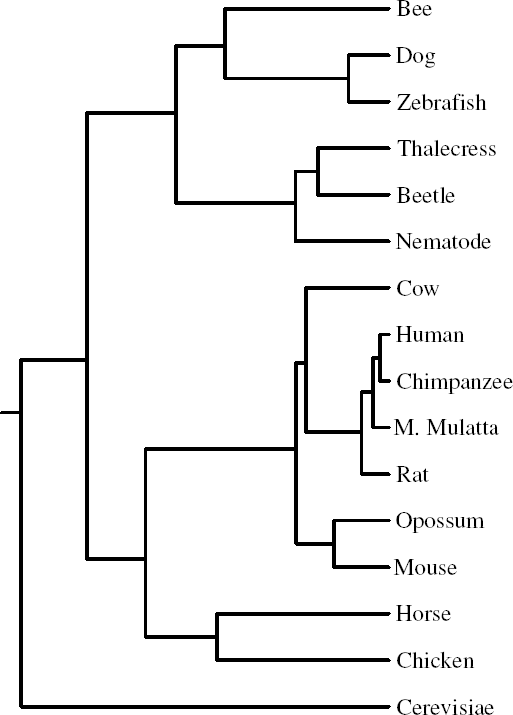
Tree of eukaryotes based on their distances in the nondimentionalized evolution space Fig 3D. This tree is considerably reasonable.

**Fig 4A.**
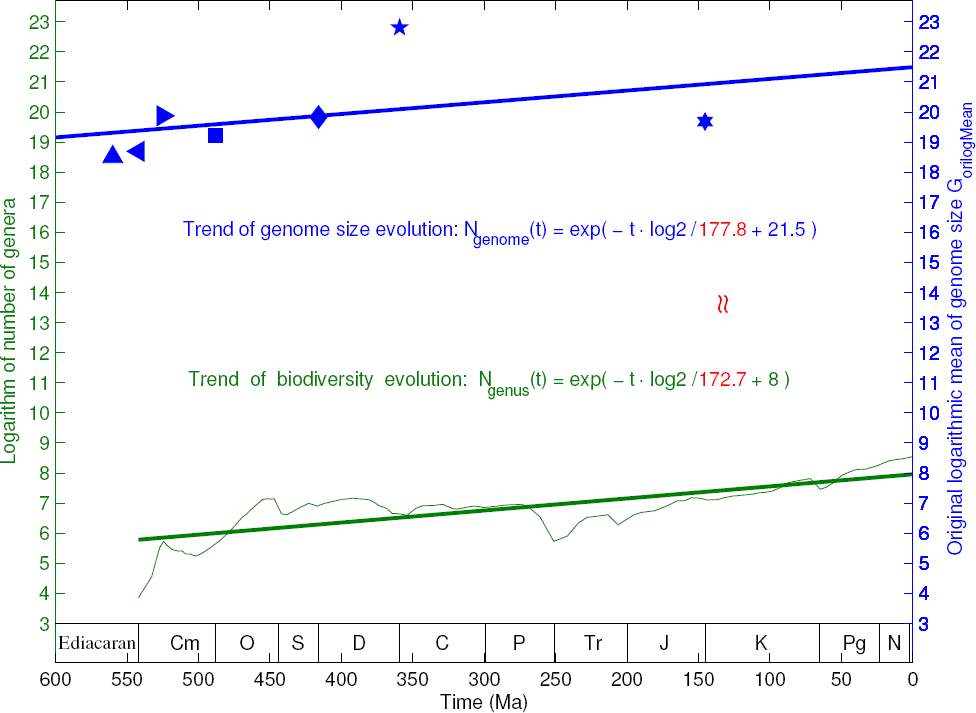
Explanation of the trend of the Phanerozoic biodiversity curve by the trend of genome size evolution. The exponential growth rate of the Phanerozoic biodiversity curve is about equal to the exponential growth rate of genome size evolution (indicated in red). Refer to Fig S4a7 and Fig S4a8 to see the exponential trend of the genome size evolution and the exponential trend of the Phanerozoic biodiversity, respectively.

**Fig S4a1.**
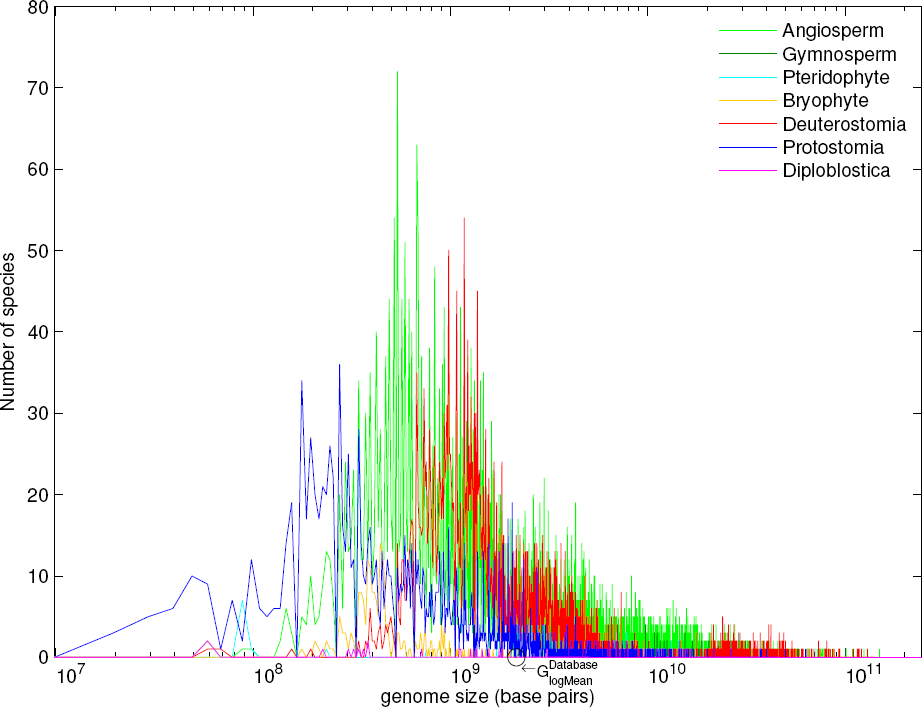
Logarithmic normal distribution of genome sizes for taxa. The genome size distribution for all the species in the two databases satisfies logarithmic normal distribution very well, due to the additivity of normal distributions. The logarithmic mean of genome sizes for all the species in the two databases is marked in the present figure, which represents the intercept of the trend of genome size evolution in Fig S4a7.

**Fig S4a2.**
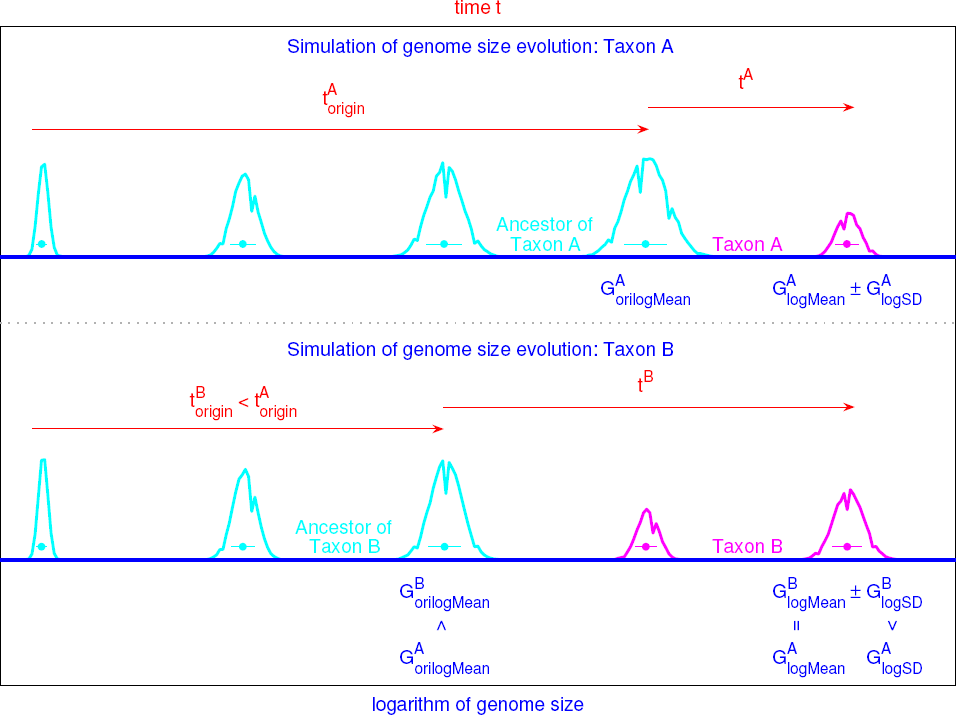
Simulation of the evolution of genome size. In the simulation, the genome size statistically doubles in the stochastic process model, which results in the logarithmic genome size distributions. The invariance of the genomic codon distributions after genome duplication is considered in the model (Fig S3a4). Two taxa *A* and *B* are compared in the simulation. The greater the logarithmic standard deviation of genome sizes *logS D t A* in a taxon is 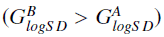, the earlier the taxon originated 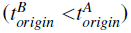. Besides, the less the original *orilogMean* logarithmic mean of genome size is 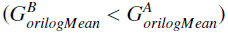, the earlier the taxon originated 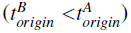.

**Fig S4a3.**
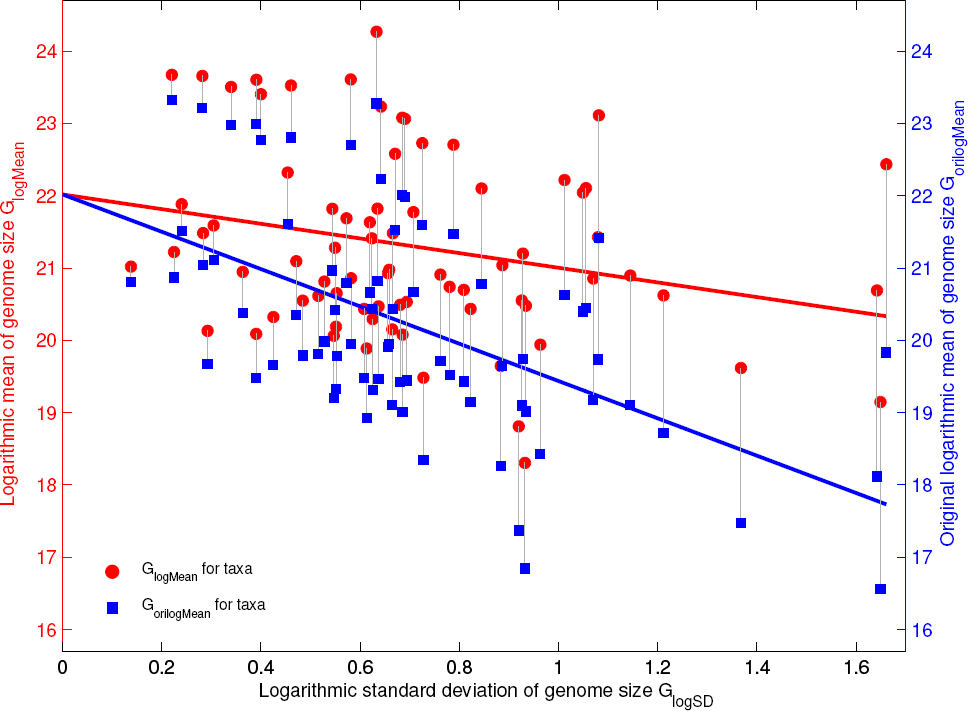
The greater the logarithmic standard deviation of genome sizes *G*_*logS D*_ is, the less the original logarithmic mean of genome sizes *G*_*orilogMean*_ is, and meanwhile also the less the logarithmic mean of genome sizes *G*_*logMean*_ is. These statistical analyses of genome sizes for the taxa are obtained based on the genomic data in the two databases for animals and plants.

**Fig S4a4.**
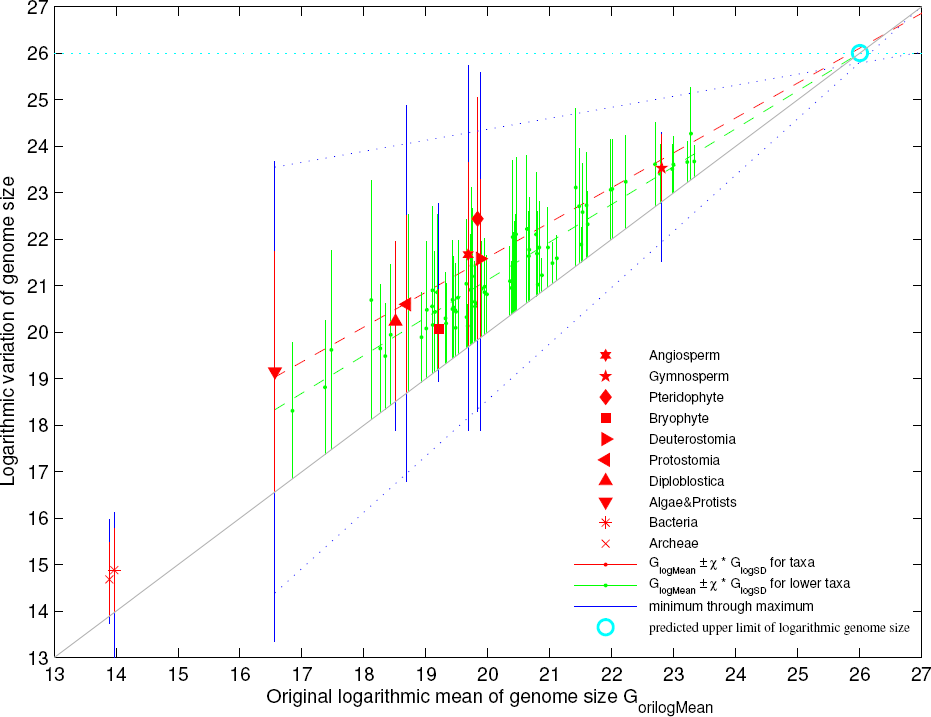
The less the original logarithmic mean of genome sizes *G*_*orilogMean*_ is, the less the logarithmic mean of genome sizes *G*_*logMean*_ is (red), and meanwhile the greater the logarithmic standard deviation of genome sizes *G*_*logS D*_ is (green or blue). The original logarithmic mean of genome sizes *G*_*orilogMean*_ indicate the origin time of taxon (Fig S4a2). The convergent point (namely the upper vertex of the triangle area in the present figure) indicates the upper limit of logarithmic genome sizes at present.

**Fig S4a5.**
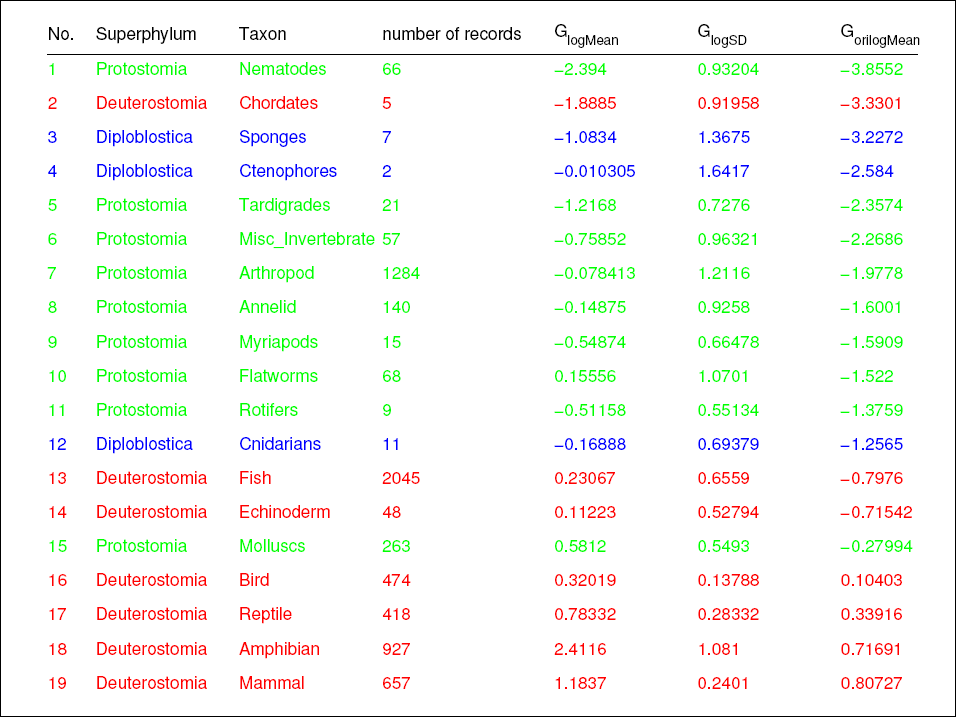
The three animal superphyla Diploblostica, Protostomia and Deuterostomia have been roughly distinguished based on the order of the original logarithmic means of genome sizes *G*_*orilogMean*_ for the animal taxa (Fig S4a4). This supports that the order of *G*_*orilogMean*_ indicate the evolutionary chronology.

**Fig S4a6.**
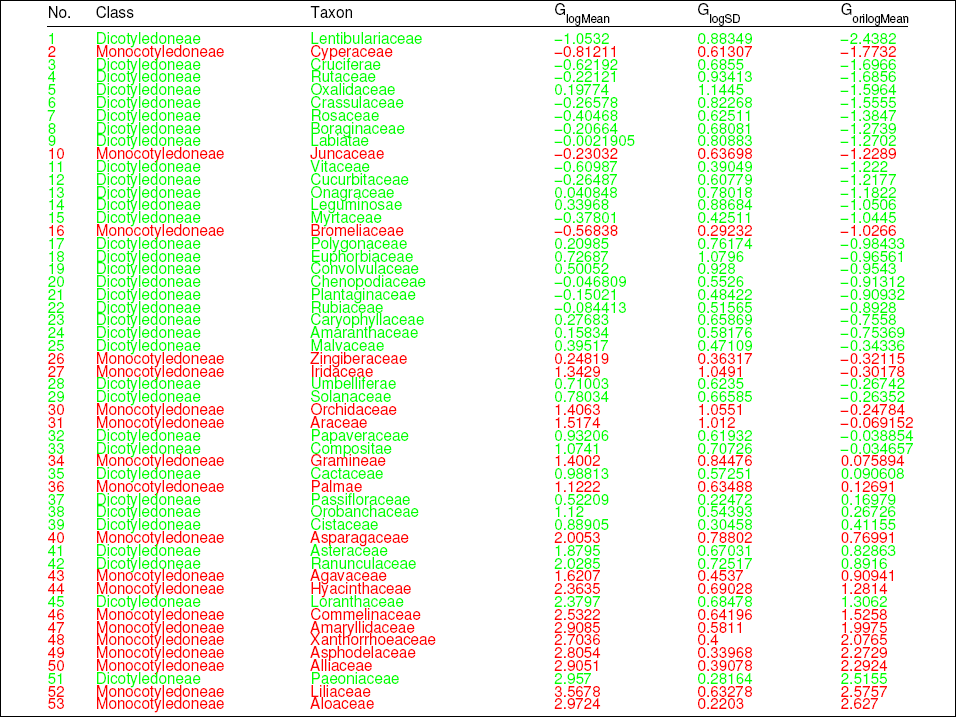
The two angiosperm classes Dicotyledoneae and Monocotyledoneae have been roughly distinguished based on the order of the original logarithmic means of genome sizes *G*_*orilogMean*_ for the angiosperm taxa (Fig S4a4). This also supports that the order of *G*_*orilogMean*_ indicate the evolutionary chronology.

**Fig S4a7.**
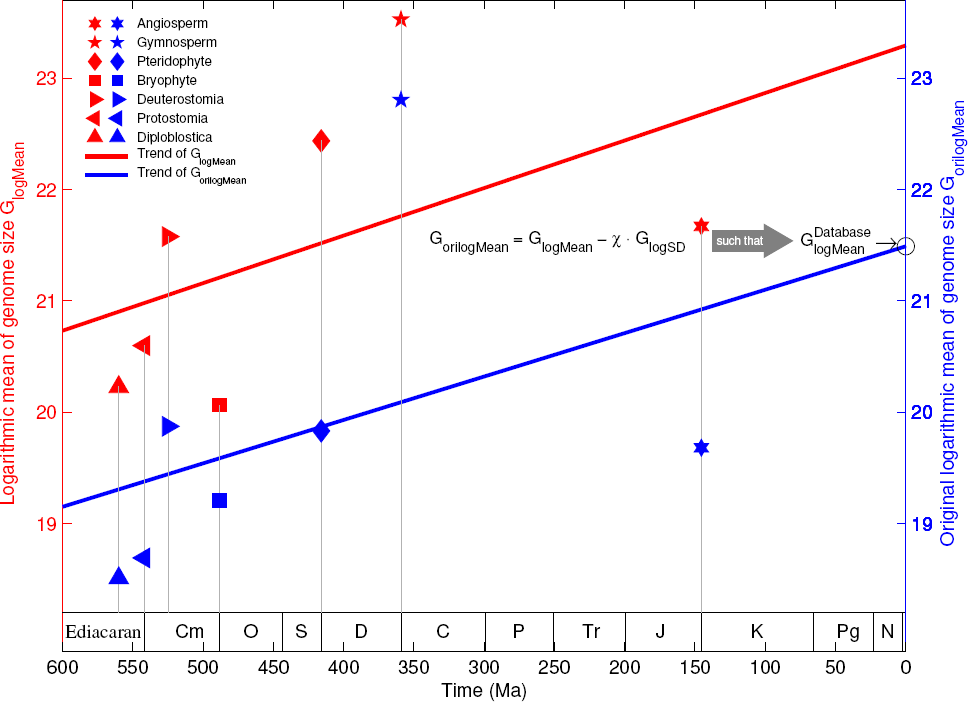
The exponential trend of genome size evolution based on the relation between the origin time of taxa and their original logarithmic means of genome sizes. The red regression line of *G*_*logMean*_ against the origin time should be shifted downwards to the blue regression line of *G*_*orilogMean*_ against the origin time, by considering the inverse relationship between *G*_*logS D*_ and the origin time (Fig 4a2). The parameter χ in the definition of *G*_*orilogMean*_ is determined by letting the intercept of the blue regression line be the average logarithmic genome size *GDatabase* (Fig 4a1).

**Fig S4a8.**
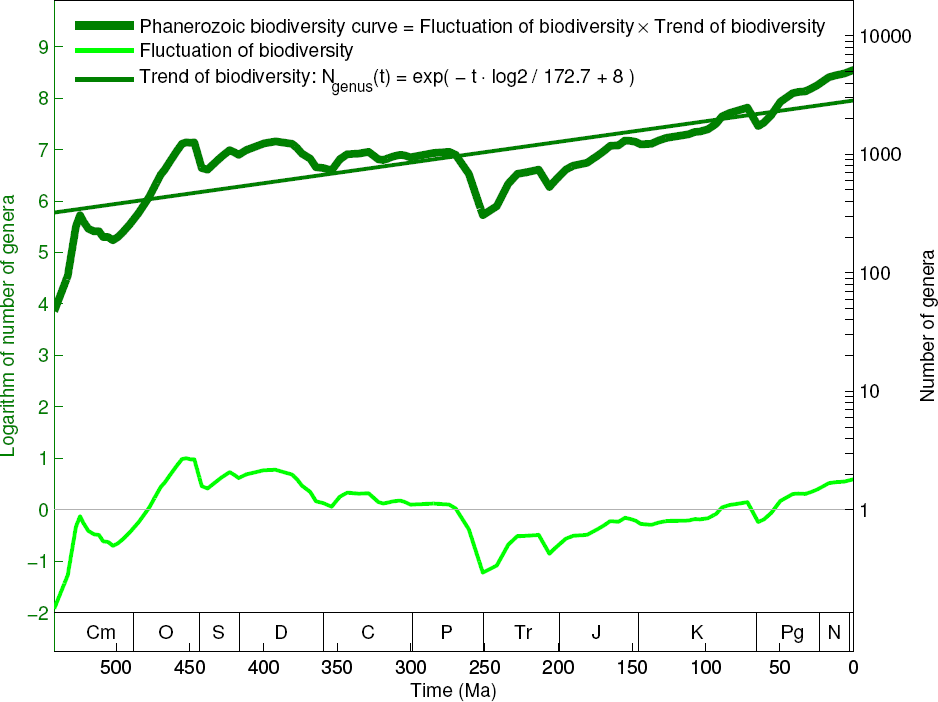
The exponential trend of Bambach et al.’s Phanerozoic biodiversity curve. The unnormalised biodiversity net fluctuation curve (thin light green) is obtained by subtracting the linear trend of logarithmic biodiversity curve (thin dark green) from the logarithmic biodiversity curve (thick dark green).

**Fig 4B.**
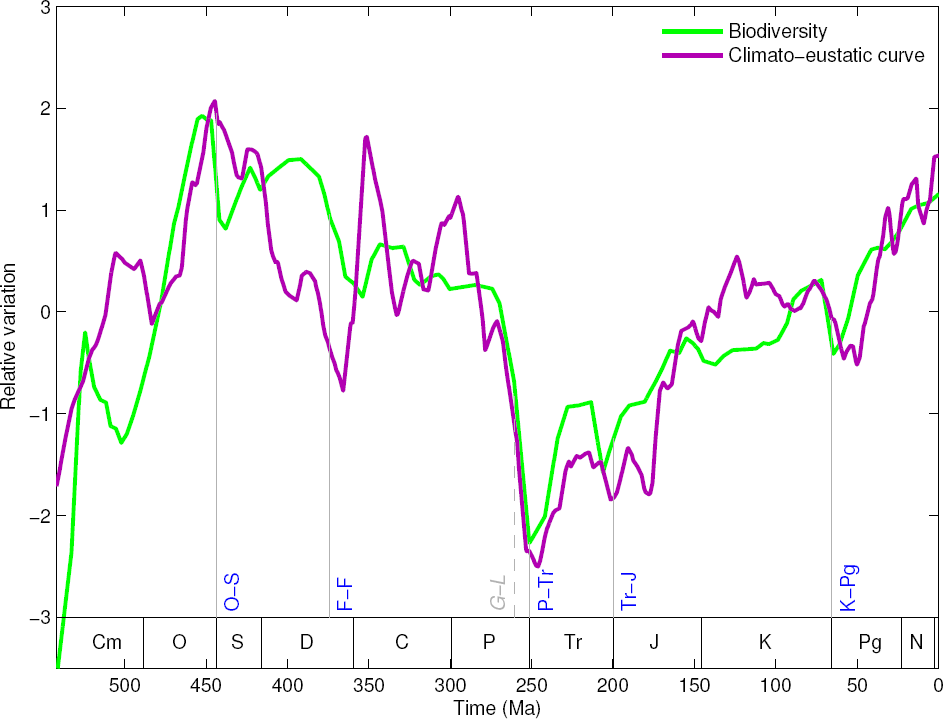
Explanation of the fluctuations of the Phanerozoic biodiversity curve based on climate change and sea level fluctuations, which indicates the tectonic cause of the mass extinctions. Refer to Fig S4b4 and Fig S4a8 to see the climato-eustatic curve and the biodiversity net fluctuation curve, respectively.

**Fig S4b1.**
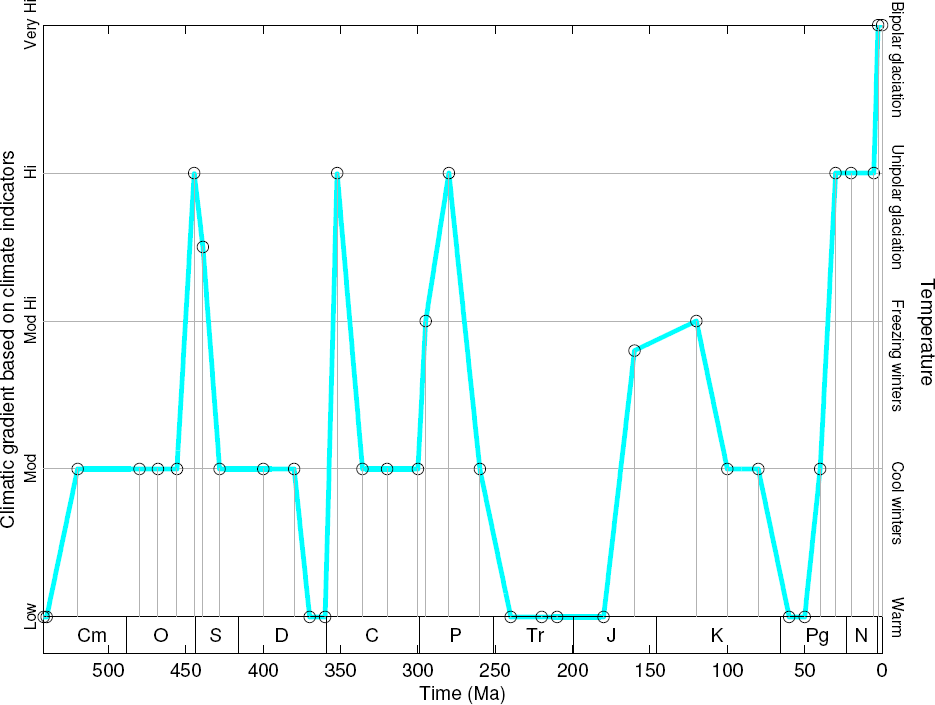
Climatic gradient curve in the Phanerozoic eon based on climate indicators. The climatic gradient curve is opposite to the climate curve.

**Fig S4b2.**
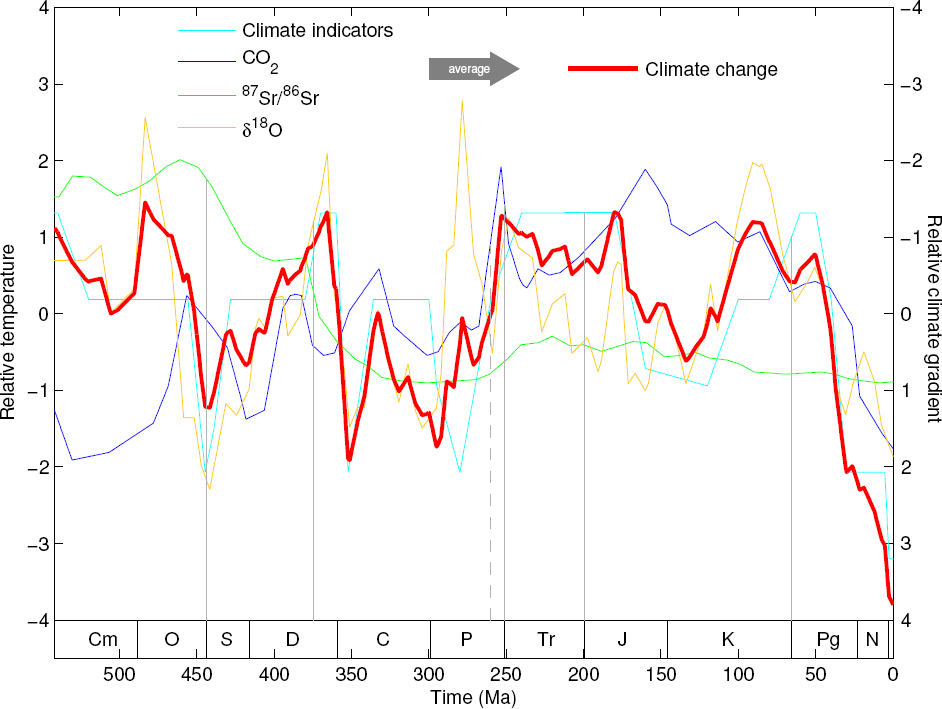
The Phanerozoic climate curve is obtained by averaging the four independent results (climate indicators, Berner’s *CO*_2_, marine ^87^*S r*/^86^*S r* and marine *δ*^18^*O*).

**Fig S4b3.**
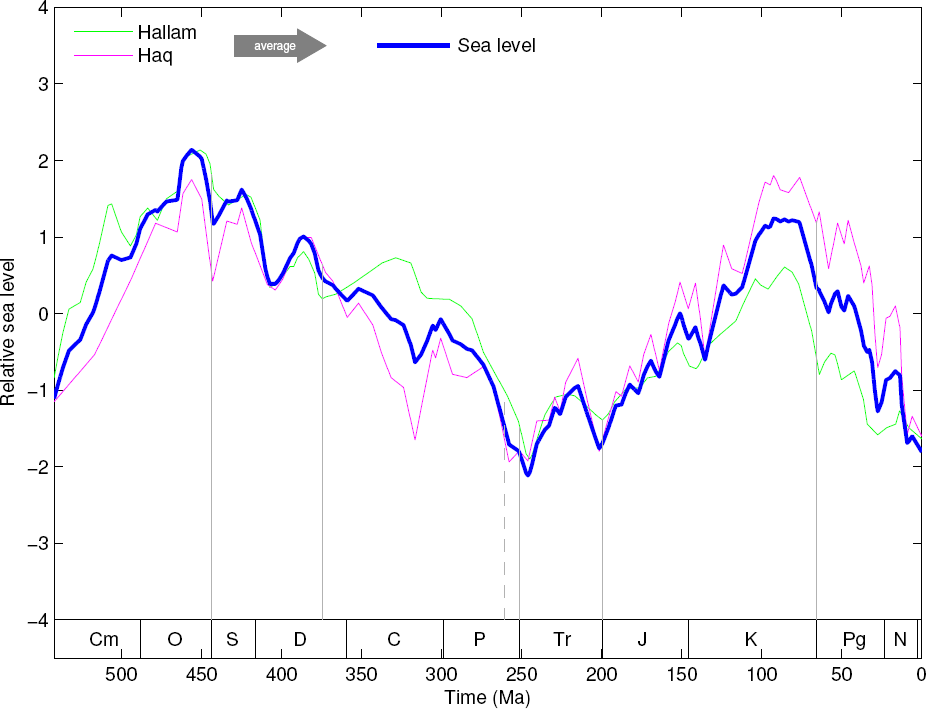
The Phanerozoic eustatic curve is obtained by averaging the Hallam’s and Haq’s results on sea level fluctuations.

**Fig S4b4.**
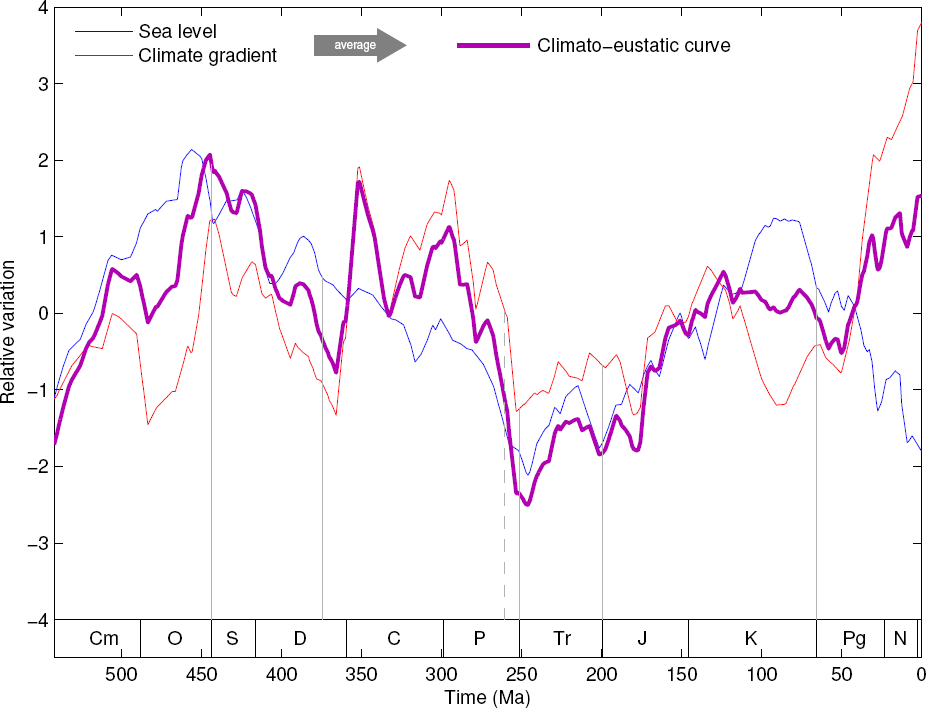
The climato-eustatic curve is obtained by averaging the Phanerozoic climate gradient curve (Fig S4b2) and the Phanerozoic eustatic curve (Fig S4b3).

**Fig S4b5.**
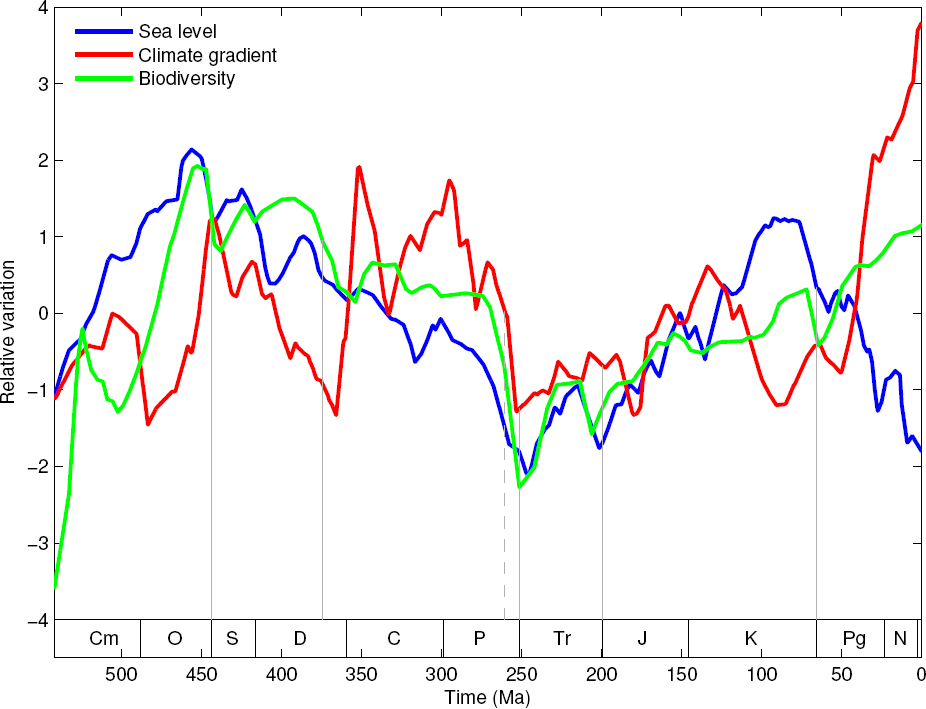
Relationships among the eustatic curve (Fig S4b3), the climate gradient curve (Fig S4b2) and the biodiversity net fluctuation curve (Fig S4a8) in the Phanerozoic eon.

**Fig S4b6.**
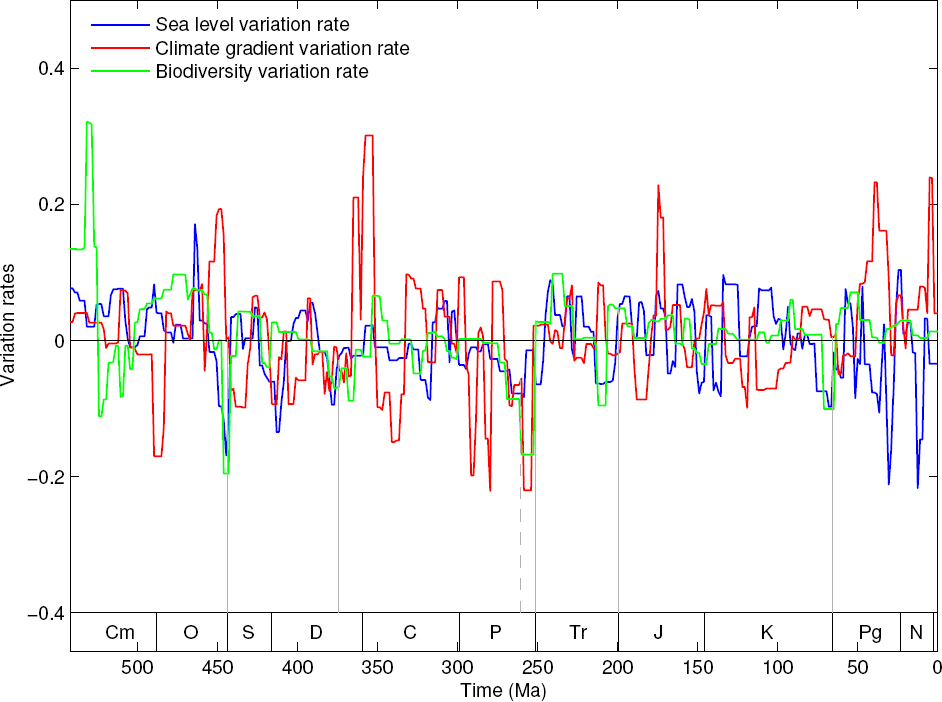
Variation rate of the eustatic curve, variation rate of the climate gradient curve and variation rate of the biodiversity net fluctuation curve in the Phanerozoic eon (the derivative curves of the three curves in Fig S4b5).

**Fig S4b7.**
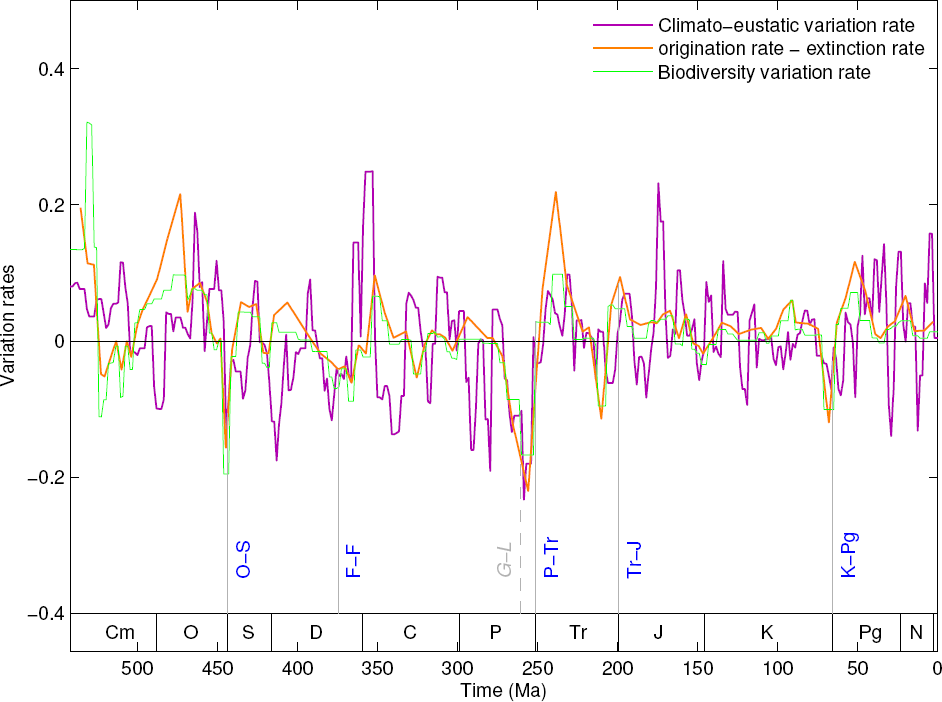
Variation rate of the climato-eustatic curve and variation rate of the biodiversity net fluctuation curve in the Phanerozoic eon (the derivative curves of the two curves in Fig 4B). The five mass extinctions (including two phases in the P-Tr extinction) are marked in the present figure (Fig 4B), and the minor extinctions can also be observed in the variation rate of the climato-eustatic curve. Additionally, the difference between the origination rate and the extinction rate is plotted (Fig S4d3).

**Fig 4C.**
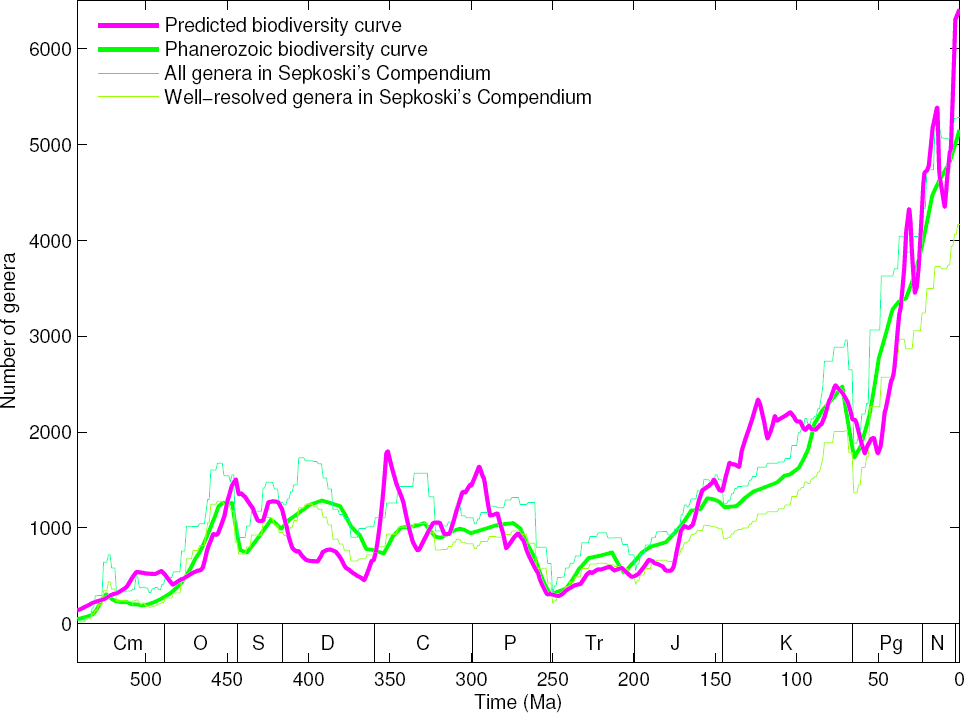
Reconstruction of the Phanerozoic biodiversity curve based on genomic, climatic and eustatic data, whose result agrees with the biodiversity curve based on fossil records. The Bambach et al.’s Phanerozoic biodiversity curve (Fig S4a8) based on fossil records is in green, and the predicted biodiversity curve in pink is based on the climato-eustatic curve in Fig S4b4 and the exponential trend of genome size evolution in Fig S4a7. Refer to Fig S4c1 for details.

**Fig S4c1.**
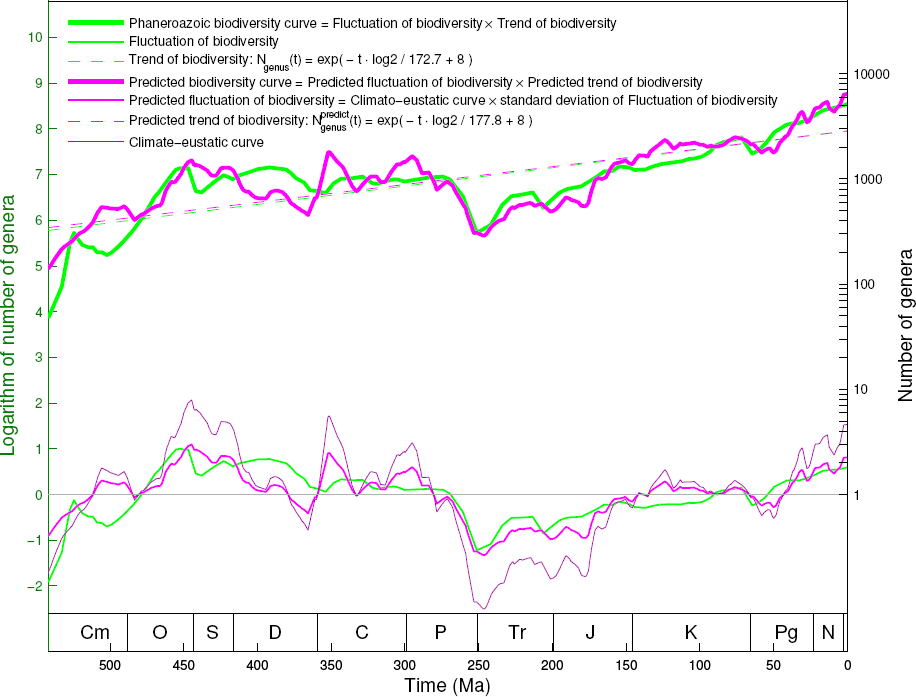
Detailed explanation of the reconstruction of the Phanerozoic biodiversity curve based on genomic, climatic and eustatic data (Fig 4A, 4B, 4C). The exponential growth trend of the predicted biodiversity curve (obtained based on the exponential trend in genome size evolution in Fig S4a7) corresponds to the exponential growth trend in the Phanerozoic biodiversity curve (obtained based on the fossil records in Fig S4a8); The fluctuations of the predicted biodiversity curve (obtained based on the climato-eustatic curve in Fig S4b4) corresponds to the biodiversity net fluctuation curve (obtained based on the fossil records in Fig S4a8).

**Fig 4D.**
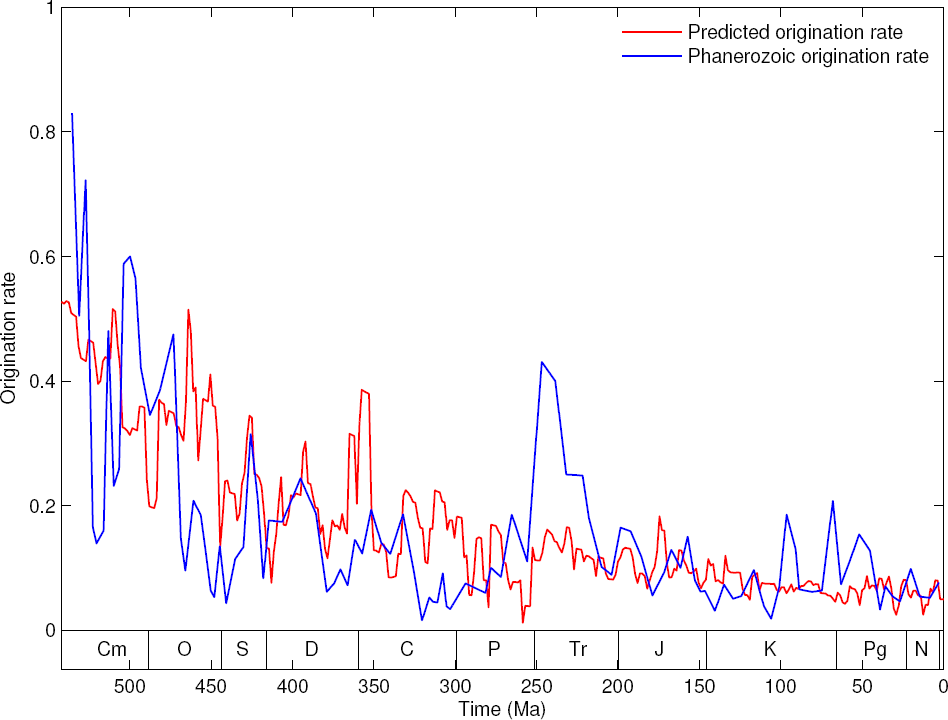
Explanation of the declining origination rate through the Phanerozoic. Refer to Fig S4d3 to see the declining origination rate and extinction rate based on fossil records; and refer to Fig S4d2 to see the explanation of these declining rates.

**Fig S4d1.**
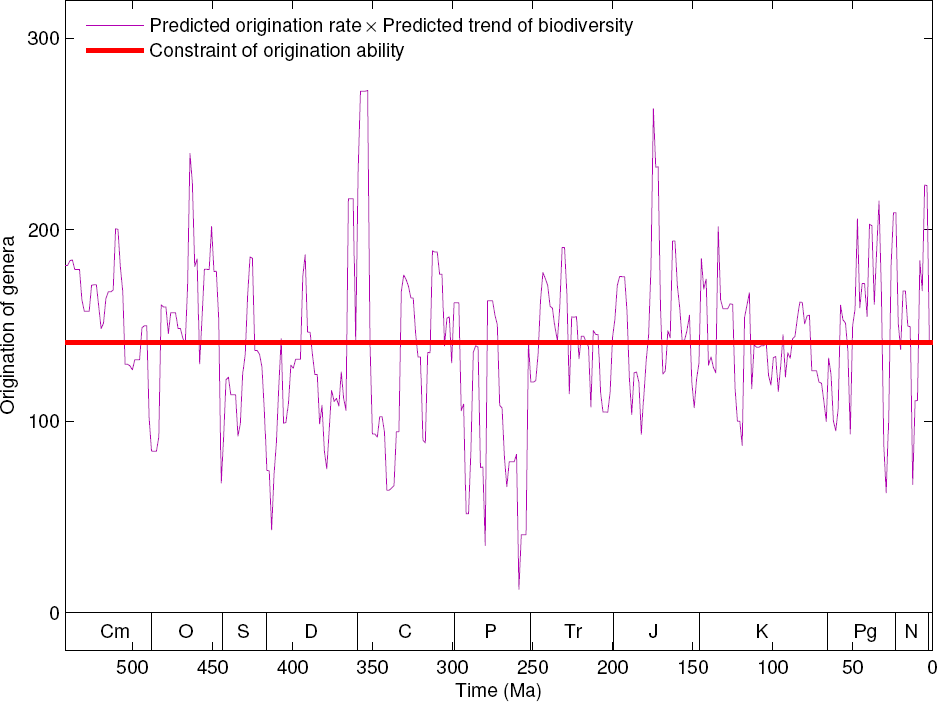
Constraint of origination ability of life on Earth, which results in the declining origination rate through the Phanerozoic eon (Fig 4D).

**Fig S4d2.**
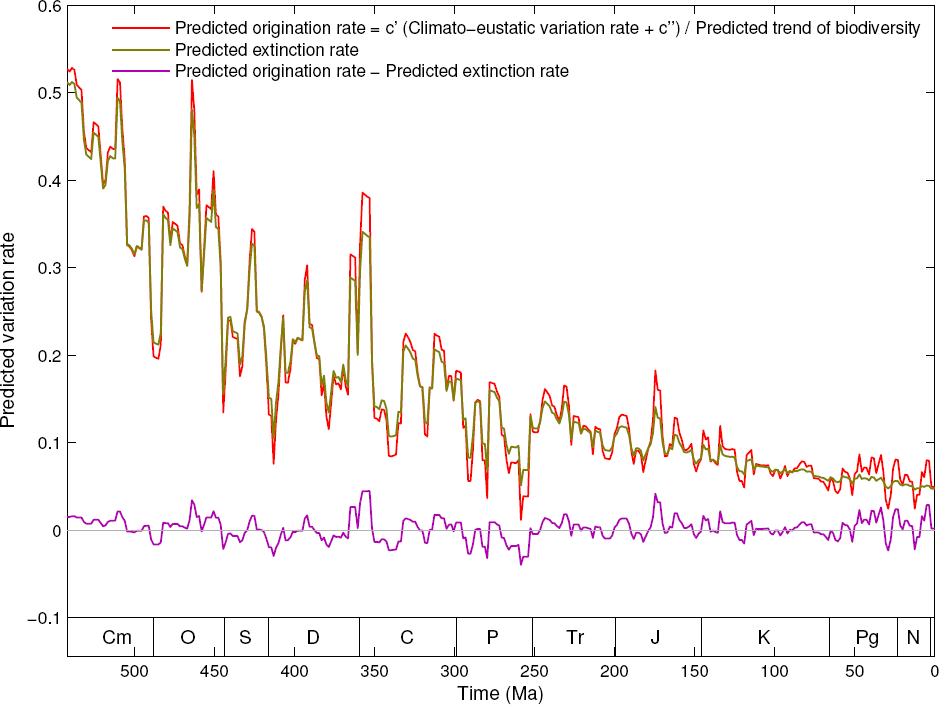
Explanation of the declining origination rate and extinction rate in the Phanerozoic eon, according to the reconstruction of the Phanerozoic biodiversity curve (Fig S4c1). The declining rates are owing to the exponential growth trend driven by the genome size evolution.

**Fig S4d3.**
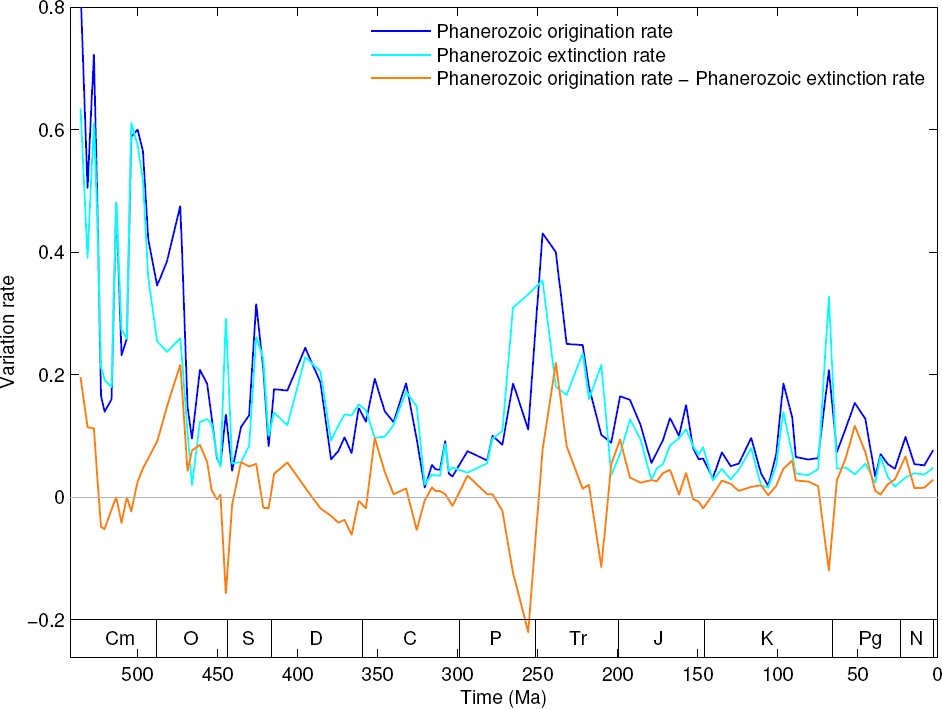
The declining origination rate and extinction rate in the Phanerozoic eon based on fossil records, which can be explained in Fig 4D and Fig S4d2.

